# High-plex spatial RNA imaging in one round with conventional microscopes using color-intensity barcodes

**DOI:** 10.1101/2024.06.29.601330

**Authors:** Tianyi Chang, Shihui Zhao, Kunyue Deng, Zhizhao Liao, Mingchuan Tang, Yanxi Zhu, Wuji Han, Chenxi Yu, Wenyi Fan, Mengcheng Jiang, Guanbo Wang, Dongfang Liu, Jirun Peng, Yuhong Pang, Peng Fei, Jianbin Wang, Chunhong Zheng, Yanyi Huang

**Author notes:** Correspondance should be addressed to (C.Z.) and (Y.H.). These authors contributed equally to this work (Tianyi Chang, Shihui Zhao, Kunyue Deng, Zhizhao Liao, Mingchuan Tang).

## Abstract

Spatial RNA imaging has not been widely adopted because conventional fluorescence microscopy is limited to only a few channels, and the cyclic reactions needed to increase multiplexing in techniques such as sequential fluorescence in-situ hybridization (FISH) require sophisticated instrumentation. Here, we introduce ‘Profiling of RNA In-situ through Single-round iMaging’ (PRISM), a method that expands coding capacity through color intensity grading. Using a radius vector filtering strategy to ensure the distinguishability of codewords in color space, PRISM achieves up to 64-plex color-barcoded RNA imaging in a single imaging round with conventional microscopes. We validate PRISM’s versatility across various tissues by generating a 3D atlas of mouse embryonic development from E12.5 to E14.5, a quasi-3D tumor-normal transition landscape of human hepatocellular carcinoma (HCC), and a 3D cell atlas and subcellular RNA localization landscapes of mouse brain. Additionally, we show the critical role of cancer-associated fibroblasts (CAFs) in mediating immune infiltration and immune response heterogeneity within and between tumor microenvironments.

## Main

The spatial arrangement of RNA transcripts is essential for revealing biological interactions at cellular and sub-cellular levels, exposing the diverse heterogeneity and structural compositions across various tissues. As demand for spatially resolved methods within the scientific community continues to grow^1–6^, existing methods face significant challenges regarding accessibility due to complex instrumentation, high costs, and low throughput^7,8^, particularly with imaging-based approaches^9^. Standard fluorescent microscopes, for instance, are generally limited to resolving no more than five channels due to spectral overlap^10,11^. Consequently, increasing multiplexity typically requires multiple rounds of labeling-stripping reactions or the addition of fluorescence channels in existing FISH-based methods^12–18^ and in-situ sequencing approaches^19–27^.

Although multiplexing potential can theoretically be increased exponentially, fluidics-dependent methods have limited widespread adoption because they rely on sophisticated, delicate equipment that combines fluidics, optics, and temperature regulation, which are typically costly and time-intensive^9^. Ensuring precise registration of signal spots across reaction cycles is further complicated by sample deformation or displacement over extended experiment durations. Additionally, three-dimensional RNA profiling presents unique challenges, such as slow reagent diffusion in thick tissues and the alignment of three-axis imaging in fluidics-dependent techniques.

Researchers are increasingly turning to fluidics-free methodologies. To exceed traditional channel limitations and encode more molecular species, some additional microscopic imaging modalities, such as super-resolution^28^, fluorescence lifetime^29^, and coherent Raman scattering microscopy^30^, have been employed. Recently, pseudo-thermal-plex channels^31^ and binding-affinity labeling^32^ were used to expand multiplexity on traditional microscopes. While these efforts achieve higher coding capacities without fluidics, the improvement in multiplexing remains modest. Furthermore, the requirement for specialized microscopy setups and temperature control equipment restricts broader application in biological laboratories.

Here we present PRISM (**P**rofiling of **R**NA **I**n-situ through **S**ingle-round i**M**aging), a high-multiplex FISH method that employs a multi-channel color barcoding on rolling circle products to distinguish dozens of RNA transcript species in a single imaging round. As a fluidics-free approach, PRISM addresses accessibility challenges by requiring only a standard fluorescence microscope, with the entire workflow from sample preparation to data acquisition completed within a day. We tested PRISM across various tissue samples, including mouse brains, mouse embryos and human hepatocellular carcinoma (HCC) samples. Using a 30-gene panel, we developed a 3D cell atlas of mouse embryos at developmental stages E12.5, 13.5, and 14.5, encompassing over 4.2 million cells categorized into 12 types, showcasing cellular compositions and organizations in different organs at various developmental phases. Additionally, using a 31-gene panel on 20 consecutive sections from an HBV-positive patient, we created a quasi-3D HCC tumor-normal transition landscape with 1.2 million annotated cells across 32 cell types, uncovering heterogeneities in the tumor microenvironment and demonstrating how cancer-associated fibroblasts (CAFs) construct a physical barrier against immune infiltration. PRISM’s compatibility with intact 3D imaging was further validated using 100-µm-thick mouse brain slices, where a 30-gene panel effectively identified over 20 cell types within the cortex, hippocampus, thalamus, and hypothalamus regions in a single staining round using conventional confocal microscopy. With a coding capacity reaching up to 64-plex, PRISM has been validated across different tissue types and thickness. It stands as a robust and accessible high-multiplex FISH method for in-situ imaging of RNA transcripts, relying solely on standard laboratory equipment.

## Results

### Coding principles of PRISM

The PRISM workflow begins by coding the targeted gene transcripts using padlock probes, each containing multiple segments of specific sequences. The barcode of each probe is a combination of segments, with each segment corresponding to a fluorescence channel (color). Although the number of available channels is limited due to spectral indistinguishability among fluorophores, each channel can accommodate multiple unique segment sequences, each encoding specific intensity information (Fig. 1a). By using this spectral barcoding scheme, the coding capacity can greatly exceed the channel limit through the combination of segmental sequences.

**Fig. 1.**
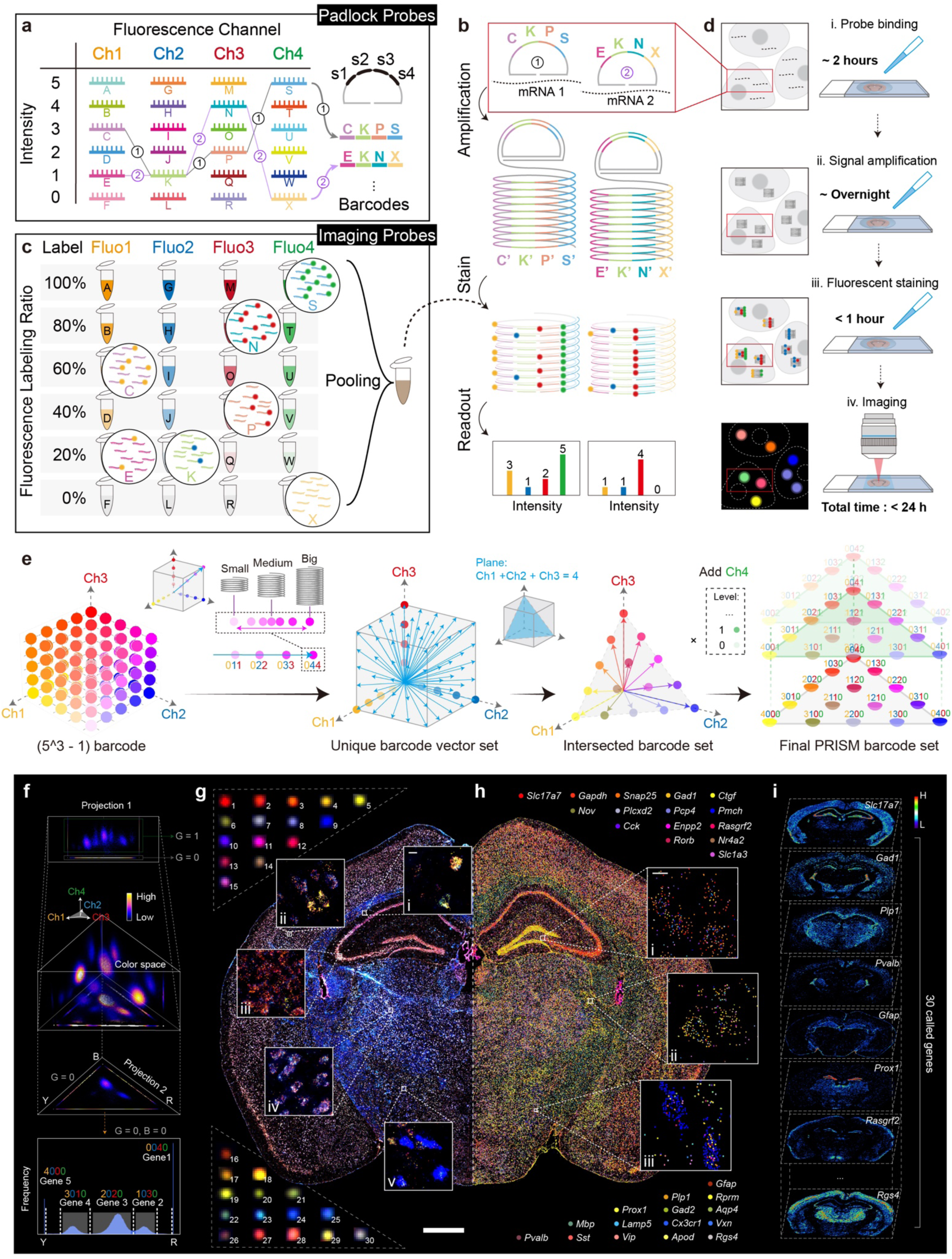
Workflow and principle of PRISM encoding. **a**, PRISM barcoding on padlock probes. Barcode on padlock probe consists of four segments, with each segment corresponding to one spectral channel. Each segment can accommodate various sequences (denoted as A, B, C,… in Ch1), with each sequence represents a distinct intensity level. The barcode on padlock probe is a combination of different segment sequence (intensities) under each channel, e.g. C-K-P-S, signifies level 3 in Ch1, level 1 in Ch2, level 2 in Ch3 and level 5 in Ch4. Therefore, each barcode is spectroscopically unique, visualized as a color. **b**, Molecular events in the PRISM workflow. Gene transcripts are targeted using a set of padlock probes with unique barcodes. Following ligation, each padlock probe undergoes rolling circle amplification (RCA) to produce hundreds of identical barcode sequences. These barcodes are then read out in-situ through fluorescent staining and imaging. **c**, Decoding through staining with pooled imaging probes. Each kind of imaging probe is designed with sequences identical to each segment sequence, with two types: fluorescent labeled and unlabeled. Two types are pre-mixed in ratios according to the designed intensity level for that segment sequence. During staining, competitive binding ensures that the desired fraction of fluorophore binds to the rolling circle product, thereby exhibiting the intended intensity. All imaging probes are pooled together for one-pot staining. **d**, Experimental workflow of PRISM. Major steps include padlock probe binding, signal amplification (probe ligation and rolling circle amplification), fluorescent staining, and imaging. The entire experiment can be completed within 24 hours using standard equipment. **e**, Radius vector filtering to ensure ample barcode discriminability. Barcode candidates result from combinations of intensity levels and spectral channels. Due to the amplicon size heterogeneity during isothermal RCA, the absolute signal intensity varies, causing the spot from a certain barcode to shift along its radial ray in the color space, leading to indistinguishability. To address this, all barcodes were represented as their radius vector, forming a unique set of barcode vectors. Barcode vectors with small neighbor angles are difficult to distinguish. To maximize the dispersion of vectors, the barcode set intersected by plane (Ch1 + Ch2 + Ch3 = K) was selected. By quantizing three channels into quarters and setting K to 4, 15 highly distinguishable barcodes can be generated. The addition of a fourth channel, Ch4, multiplies the coding capacity by Ch4 quantities. For example, with a 2-intensity-level, which is 0 (No) or 1 (Yes), the total available barcodes are 30 or 31 (including only Ch4 itself). **f**, Gene decoding in color space. The signals from four colors were converted into three-dimensional coordinates, and each spot was plotted in a “color space”. 30 clusters located in different positions, each representing a unique barcode. Each barcode was identified by its specific location within the color space. **g**, Spots in the raw image corresponded to 30 different barcodes. Each spot consists of four channels, shown within the same dynamic range. Scale bar: 10 µm, zoom-in image; 1 mm, below. **h**, Decoded genes. Each gene was assigned a pseudo-color corresponding to its barcode. Scale bar: 10 µm. **i**, Spatial distribution of 30 decoded genes.

Each transcript-bound padlock probe is then clonally amplified in situ via rolling circle amplification (RCA), creating a large number of identical barcodes for each clone (Fig. 1b). An array of imaging probes is also prepared, each containing a pair of identical single-stranded oligos, one with and one without fluorescent labels (Fig. 1c, Supplementary Fig. 1). The ratio of labeled to unlabeled twin-probes is carefully determined, and all probes are pooled together as a single staining mix. When this mix is applied to hybridize with the RCA amplicons in situ, each segment binds both labeled and unlabeled probes in the preset ratio, forming a spectral barcode for each clone through competitive hybridization. Quantitative assessment of fluorescence intensity across all channels in each clone then reveals its barcode (Fig. 1c,d).

Theoretically, an array with *m* channels and *n* intensity-level segments can provide a coding capacity of *n^m^*−1. For example, with three spectral channels, each quantized into quarters (0, 1/4, 2/4, 3/4, 4/4), it is possible to write codes with 3-digit quinary numbers, enabling encoding of up to 5^3^ − 1 = 124 genes. However, in practice, the number of usable barcodes is reduced due to variations in the sizes of amplified nanoball clones, which can cause certain codes indistinguishable in spectral color space (Fig. 1e). For instance, a small amplicon barcoded as (020) would be indistinguishable from a twice-sized amplicon barcoded as (010), as would (033) and (022).

To better describe barcode discriminability under these conditions, each barcode is conceptualized as a ‘radius vector’ in color space. Indistinguishable clones share the same vector direction but differ in magnitude; therefore, only angularly distinct barcode vectors are considered legitimate (Fig. 1e, Extended Data Fig. 1, Supplementary Fig. 2). To ensure ample dispersion of barcode vectors, we constrained the coding space to the plane where the sum of intensities across three channels (Ch1+Ch2+Ch3) equals 4. This approach optimizes the angles between barcode vectors, minimizing crosstalk and maximizing the distinguishability of barcodes. As a result, with three quarterly quantized channels, we achieve 15 highly distinguishable barcodes.

With a 3-channel setup that minimizes amplification bias, the coding capacity can be further expanded by adding a fourth channel. In our demonstration, we selected common fluorophores, Texas Red (TR, Ch1), Alexa Fluor 488 (AF488, Ch2), Cy5 (Ch3), and Cy3 (Ch4), due to their wide availability. To address interference from tissue autofluorescence, the intensity level of Ch4 (Cy3) is limited to 2 or 3, effectively doubling or tripling the coding capacity to more than 30 (Supplementary Fig. 2).

Using conventional equipment and a single round of staining, we conducted cell typing on a 10-µm-thick mouse brain coronal section with a 30-gene panel^33^ (Fig. 1f-i). Fluorescent puncta were processed to determine the spectral code of each amplicon, revealing the barcode of each transcript in situ (Extended Data Fig. 2). The barcodes formed into 30 distinct clusters in color space, accurately reflecting the original design. Clusters were identified using manually adjustable Gaussian fitting for probability prediction, achieving over 80% spot-preserving efficiency (Fig. 1f, Extended Data Fig. 3). By mapping each barcode back to the sample, we reconstructed the submicron-resolution spatial distribution of all 30 genes (Extended Data Fig. 4). High overall accuracy was demonstrated with amplification fidelity exceeding 95% and a PRISM decoding accuracy of 95.7%, as validated by orthogonal methods (Supplementary Figs. 3-5, Extended Data Fig. 5). With a minimal false-targeting rate (Supplementary Fig. 6), the overall accuracy was further validated through benchmarking spatial expression patterns against RNAscope results and previously reported data^27^ (Supplementary Figs. 7-9).

Cell segmentation and typing were completed by integrating 30 RNA spatial images with DAPI-stained nuclei territories (Supplementary Fig. 10). In this brain section, we identified 71,435 cells, categorizing them into five major types: excitatory neurons (39,757, 55.6%), inhibitory neurons (13,134, 18.4%), and non-neuronal cells, including oligodendrocytes (8,244, 11.5%), astrocytes (7,062, 9.9%) and microglia (3,238, 4.5%). Further differentiating into subtypes revealed the architectural organizations of the mouse brain at single-cell resolution (Supplementary Fig. 11).

### Spatial profiling of mouse embryos enables intracellular interaction analysis

We used PRISM with a specific panel of 30 marker genes to explore the intricate structural dynamics and cellular interactions within various tissues and organs during embryonic development^34,35^ (Fig. 2a-d, Supplementary Figs. 12-14). Spatial expression patterns are well-correlated with previously reported atlas^35^, offering finer details of tissue morphological structures (Supplementary Figs. 15, 16). A 10-µm whole-mount section of an E13.5 mouse embryo revealed 12 primary cell types, including the epidermis, bone, and gastrointestinal tract, distinguished by the most abundantly expressed genes (Fig. 2c,d). Notably, the nervous system, constituting 39.3% of the total cell count, can be further classified based on marker gene expression, revealing spatial organization within the embryonic brain at E13.5, spanning the forebrain (pallium, subpallium), midbrain, hindbrain, and spinal cord (Fig. 2e). Gene co-expression analyses were conducted to determine the co-localization of genes within the nervous system (Fig. 2f), with significant co-localization observed for *Hoxb8*, *Robo2*, and *Stmn2* in the spinal cord region. Additionally, the co-presence of *Gad2* and *Dlx1* in the subpallium supported the hypothesis that DLX1 transcription factors directly influence *Gad2* expression, a crucial interaction for the development of GABAergic neurons in the forebrain and illustrated a key aspect of neuronal differentiation and specialization^36^.

**Fig. 2.**
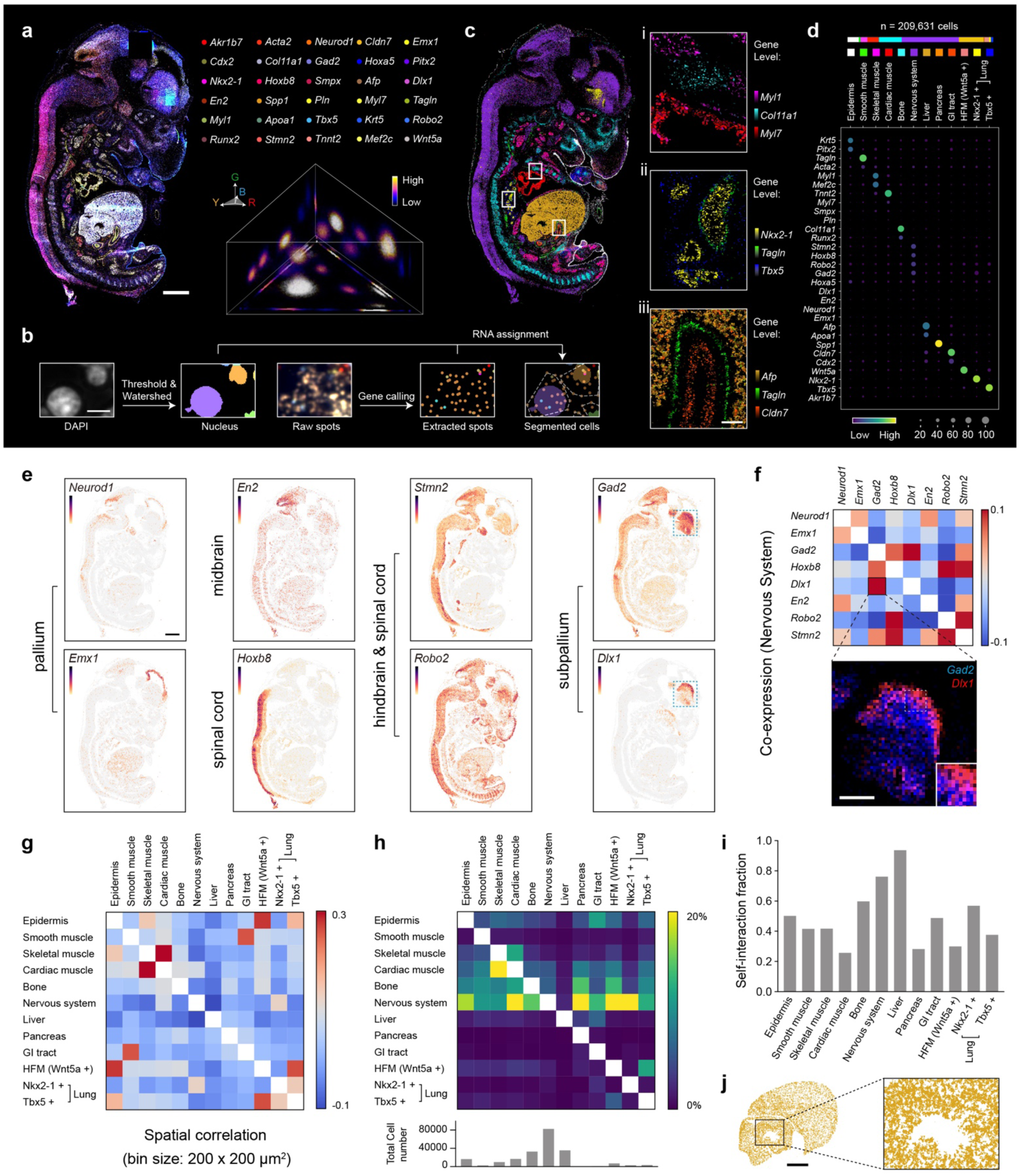
Spatial profiling of mouse E13.5 embryo by PRISM. **a**, A composite raw image of four channels of the mouse embryo E13.5 (sagittal section) by PRISM and the distribution of 30 clusters of spots in color space. Scale bar: 1mm. **b**, Cell segmentation. Cells were segmented through a DAPI image, and each identified gene spot was assigned to the nearest nucleus through Euclidean distance. Scale bar: 5 µm. **c**, Annotated cell type distribution of the mouse embryo. Cell type was determined by the most abundant gene within each cell. Zoom-in figs (i - iii) were colored based on gene abundance. Scale bar: 100 µm. **d**, Dot plot of each cell type. **e**, Heterogeneity in the nervous system can be revealed through gene expression patterns. The cells were colored based on the abundance of their respective gene expressions and grouped according to the organ’s location. Scale bar: 1 mm. **f**, Co-expression analysis revealed the developmental role of transcription factor DLX1 in forebrain GABAergic-neuron. Scale bar: 500 µm. **g**, Coarse-grain spatial correlation analysis between general cell types. The spatial binning size is 200 x 200 µm^2^. **h**, Direct intercellular interaction between general cell types. The interaction degree was defined as the proportion of cells directly adjacent to another cell. The figure below shows the total cell number of each general cell type. **i**, Direct intercellular interaction between each cell type itself. **j**, Spatial distribution of liver cells. The zoom-in image shows numerous structures of hepatic lobes, each composed of several liver cells. Scale bar: 500 µm.

A coarse-grained spatial correlation analysis was performed to understand the relationship between various cell types across tissue locations (Fig. 2g, Supplementary Fig. 17). The co-occurrence of skeletal and cardiac muscle cells, as well as GI tract and smooth muscle cells, indicates their collaborative roles in specific developmental functions. Both Wnt5a^+^ and Tbx5^+^ cells play critical roles in coordinating limb development. In our analysis, Wnt5a^+^ Limb mesenchymal cells are primarily distributed in the distal regions of the limb and the terminal mouth area but spatially proximal to epidermis cells. Additionally, Tbx5^+^ limb fibroblasts demonstrate a high spatial correlation with Wnt5a^+^ cells. They exhibit a greater abundance in the limb and are primarily distributed in the proximal regions, exemplifying their joint contribution to the formation and functioning of the limbs.

The further refined analysis assessed direct intercellular interactions (Fig. 2h), revealing a complex network of developmental interactions between the nervous system and other organs at E13.5. Notably, pancreatic cells marked by *Spp1* exhibited significant interactions with chondrocytes and osteoblasts. The presence of islands of Spp1^+^ cells within the skeletal structure suggests their possible involvement in the early formation of the intervertebral disc^37^ (Supplementary Fig. 18). The previously noted spatial correlation between Wnt5a^+^ and Tbx5^+^ cells was confirmed to be a direct interaction. Additionally, Nkx2-1^+^ cells, highly expressed in certain neural precursors^38^, were found predominantly in the subpallium and demonstrated extensive interactions with the nervous system, including the spinal cord. Notably, numerous Nkx2-1^+^ cell clusters, with their distinctive disk-like structure, are scattered across the alar plates of the spinal cord, suggesting a role in cell differentiation and migration within the spinal cord (Supplementary Fig. 18). Not surprisingly, interactions within the same cell type are considerable (Fig. 2i). Particularly, 94% of hepatocytes interact with each other in close proximity (Fig. 2j), indicating early multicellular structure formation within the hepatic lobe and the establishment of basic hepatic architecture by E13.5.

### Spatial-temporal atlases illustrate embryonic organ development

Single-round 30-plex imaging has enabled rapid spatial profiling on a large scale. We created a sampled 3D cell atlas of an E13.5 mouse, utilizing 11 whole-mount sections spaced 280 µm apart, capturing a total of 2,331,513 annotated cells (Fig. 3a). This comprehensive experiment, from sample preparation to imaging, was completed within three days to generate an annotated cell atlas, allowing for the visualization of various early developmental structures.

**Fig. 3.**
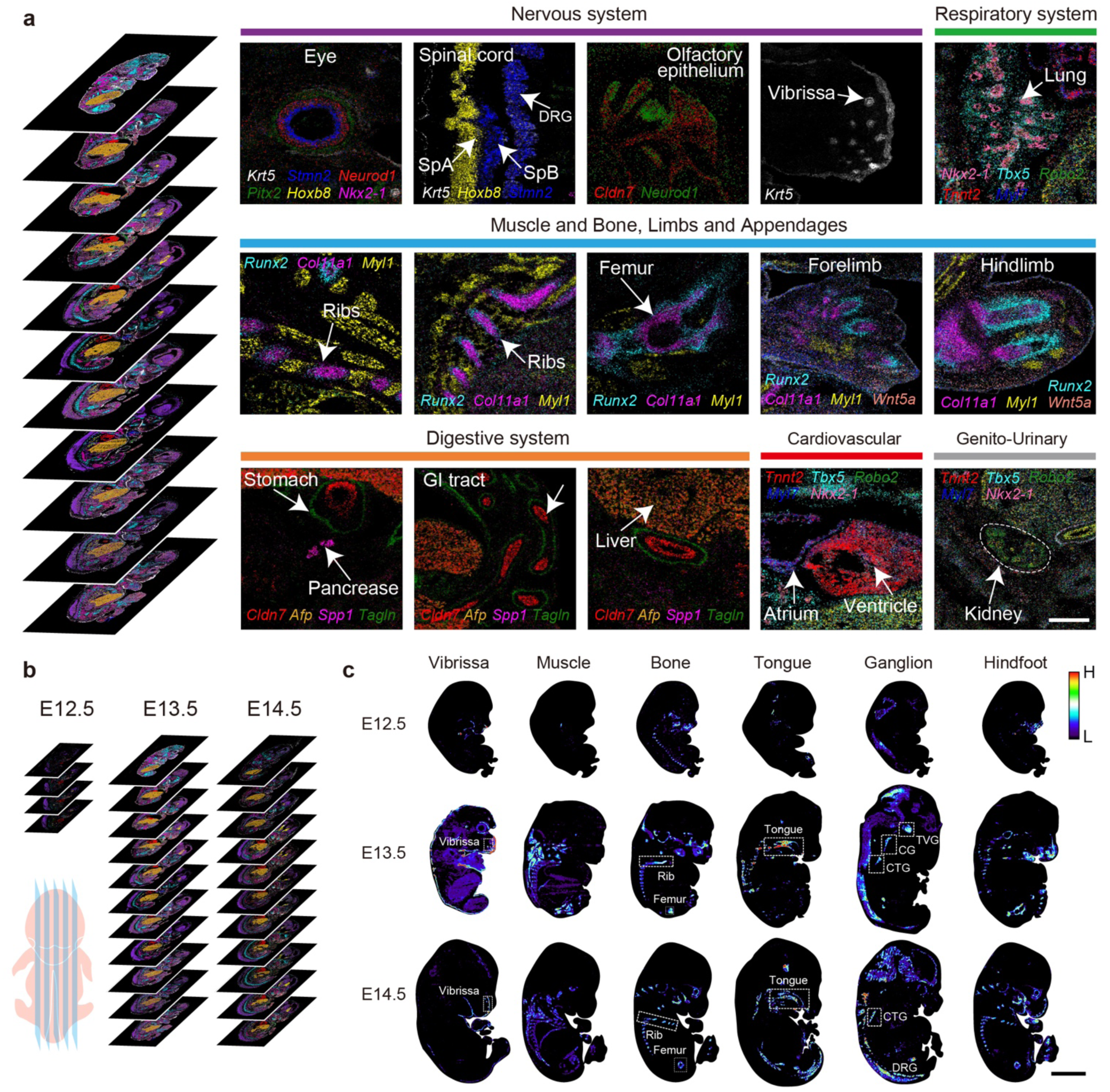
High-throughput spatial profiling of mouse embryos E12.5, E13.5, and E14.5. **a**, Developmental structures across different systems from a sampled-3D spatial atlas of the E13.5 embryo, constructed from 11 sagittal sections at an average interval of 280 µm. Fine structures from the nervous, respiratory, motion, digestive, cardiovascular, and genitourinary systems are collected from different sections. Cells within these structures are colored based on the abundance of their respective marker genes. Scale bar: 500 µm. **b**, Sampled-3D spatial atlas of E12.5 (4 sections), E13.5 (11 sections) and E14.5 (11 sections) embryo. **c**, Developmental progression at different stages is revealed by annotated cell distribution. DRG (Dorsal Root Ganglion); CG (Cervical ganglion); TVG (Trigeminal V ganglion); CTG (Cervico thoracic ganglion). Scale bar: 2.5 mm.

At the E13.5 stage, the nervous system showcases distinct structures such as the multi-layered composition of the eye (marked by Neurod1^+^, Pitx2^+^, Stmn2^+^), the spinal cord’s alar and basal plates (Hoxb8^+^, Stmn2^+^), and the olfactory epithelium (Neurod1^+^, Cldn7^+^). Notably, *Pitx2* (cornea), *Neurod*1 (outer layer), and *Stmn2* (inner layer) of the optic cup exhibit a gradient of expression from outer to inner layers, contributing to eye development. Conversely, in the pallium, an inverse pattern of expression was observed, likely due to the invagination process during the optic cup formation^39^ (Supplementary Fig. 19). The respiratory system at this stage, indicated by *Nkx2-1* and *Tbx5* expression, reveals lung-lobe and mesenchymal cells’ distribution. The early motion system, including muscles, bones, and appendages, can be observed with cellular resolution from various angles, laying the groundwork for future movement and coordination. Skeletal development is marked by specific gene expressions that present the differentiation direction of cell types: *Runx2* for osteoblasts and *Col11a1* for cartilage. The digestive system begins to differentiate into the gastric and intestinal primordia, with *Cldn7* marking this early form. The circulatory system, particularly the heart, shows a clear chamber structure and atrium/ventricle differentiation (Myl7^+^, Tnnt2^+^), while the renal system’s fine structure is delineated by Robo2^+^ and Nkx2-1^+^ expressions. PRISM’s high resolution and accuracy effectively reveal mouse embryo’s intricate structures and cell types at E13.5.

Expanding the atlas to include E12.5 and E14.5 mouse embryos offers valuable insights into their developmental progression, creating a temporal-spatial atlas with 26 whole-mount sections, 4,257,418 cells, and 107,655,795 transcripts from 30 RNAs at sub-micron resolution (Fig. 3b, Supplementary Fig. 20). The period from E12.5 to E13.5 witnessed significant transformations in various tissues (muscle, skeleton, and ganglia) and organs (such as vibrissa and tongue) (Fig. 3c). By E12.5, the pancreas was identifiable as a cluster of Spp1^+^ cells, which subsequently enlarged and underwent bifurcation by E14.5 (Supplementary Fig. 21). Also, a group of Wnt5a^+^ cells in the hindbrain has a significant migration behavior, suggesting morphogenic behavior of the dorsal hindbrain during development^40^. Lung development is characterized by volume reduction and increased branching and bifurcation within the lung structure (Nkx2-1^+^, Tbx5^+^), and a more developed trachea is observed (Nkx2-1^+^). At the same time, the GI tract shows rapid development (Cldn7^+^, Tagln^+^), highlighting the dynamic nature of embryonic growth and organ formation.

### PRISM exhibits complex histological profiling of liver tumor microenvironment

We conducted a detailed analysis of a hepatitis B virus (HBV) positive hepatocellular carcinoma (HCC) sample (around 6 mm x 5 mm) using a 31-gene panel, comprising 30 selected markers associated with cell proliferation, liver tumor cells, cancer-associated fibroblasts (CAFs), endothelial cells, and immune cells, in addition to a probe targeting a 28-bp sequence encoding the HBV core protein^41,42^ (Fig. 4a, Supplementary Figs. 22-24). This comprehensive panel facilitated the spatial exploration of the tumor microenvironment at single-cell resolution.

**Fig. 4.**
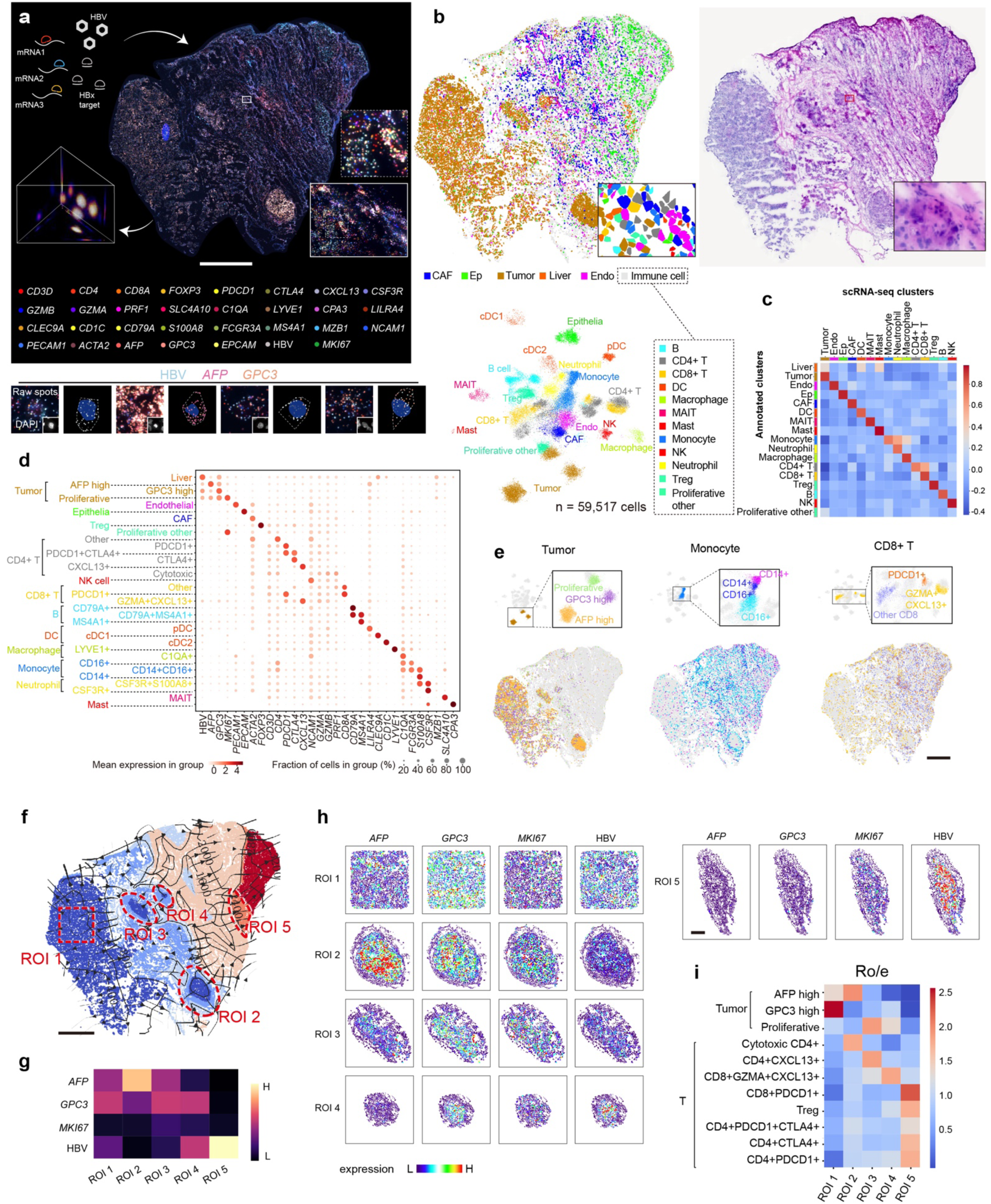
Spatial profiling of human hepatocellular carcinoma cancer microenvironment by PRISM. **a**, Workflow and signal results of HCC sample profiled by PRISM 31-gene panel, targeting 30 marker genes and one HBV core-protein-encoding RNA. Scale bar: 1 mm. **b**, Cell type classification in the HCC sample. Upper left: cell annotations derived via Harmony embedding and Leiden clustering. Bottom left: UMAP projection of all cells. Upper right: H&E staining of an adjacent tissue section. **c**, Pearson correlation between PRISM-based cell classifications and single-cell transcriptomic data. **d**, Dot plot displaying the expression of marker genes across each cell subtype. **e**, Spatial projection of cell subtypes across the tissue section. Scale bar: 1 mm. **f**, GASTON isodepth and cluster plot with annotated ROIs. Scale bar: 1 mm. **g**, Expression levels of *AFP*, *GPC3*, *MKI67*, and HBV core protein RNA across 5 ROIs. **h**, Spatial gene expression levels of *AFP*, *GPC3*, *MKI67*, and HBV RNA content within 5 ROIs. Scale bar: 200 µm. **i**, Ro/e (ratios of observed cell numbers to random expectations) cell type enrichment analysis across the five ROIs.

Our analysis identified 1,695,418 transcripts and assigned them to 60,329 cells, predominantly classified into 12 immune cell populations—including B cells, T cells, natural killer cells, mast cells, neutrophils, monocytes, and macrophages—and 5 non-immune cell types, such as CAFs, tumor cells, epithelial cells, endothelial cells, and normal liver cells (Fig. 4b, Supplementary Figs. 25, 26). This classification was achieved using Harmony embedding and unsupervised clustering analysis based on distinct PRISM gene marker profiles. The accuracy of cell identification by PRISM closely aligned with findings from prior single-cell RNA-sequencing studies^41–43^ (Fig. 4c), validating the effectiveness of this single-round imaging technique.

By mapping single cells to their original spatial contexts, we found that the spatial distributions and positions of cells, as revealed by PRISM, well correlated with those derived from conventional fast-diagnostic hematoxylin and eosin (H&E) and immunohistochemistry (IHC) staining methods (Fig. 4b, Supplementary Fig. 27). Further analysis delineated dominant cell populations into subtypes based on their unique PRISM profiles (Fig. 4d, Supplementary Fig. 28). For instance, tumor cells were subdivided into three distinct clusters characterized by the high expression of *AFP*, *GPC3,* and *MKI67*, indicating intrinsic heterogeneity. Immune cells, such as monocytes, were further categorized into classical (CD14^+^CD16^-^), non-classical (CD14^dim^CD16^+^), and intermediate (CD14^+^CD16^+^) subtypes. Additionally, cytotoxic, exhausted and CXCL13^+^ tumor-reactive CD8 T cells were identified as unique subpopulations among CD8 T cells (Fig. 4e). To probe cellular crosstalk, Squidpy neighborhood enrichment analysis revealed that AFP-high tumor cells preferentially co-localized with specific cell clusters, such as plasmacytoid dendritic cells (pDCs), and CD4⁺CXCL13⁺ T cells (Supplementary Fig. 29), suggesting their potential spatial interaction. This nuanced classification and the investigation of spatial colocalization among individual cell types, especially between different subtypes, enrich our understanding of the complex dynamics within the tumor microenvironment.

Using the GASTON method^44^, we profiled spatial features across the tissue (Fig. 4f). Four distinct tumor regions and one normal region of interest (ROIs) were selected based on their GASTON-defined altitudes. Tumor ROIs exhibited pronounced heterogeneity in cellular composition and spatial distribution that ROI1 was enriched for GPC3-high tumor cells, whereas ROI2 was dominated by AFP-high cells (Fig. 4g). Notably, the normal liver ROI (ROI5) exhibited elevated HBV core protein expression, suggesting early HBV activity preceding HCC development (Fig. 4h). The immune landscape also varied by regions, with greater immune cell infiltration observed in ROI5 (Fig. 4i). Exhausted T cells expressing *PDCD1* and/or *CTLA4* were more abundant in the HBV-positive ROI5 than other ROIs. In contrast, CXCL13^+^ CD4^+^ and CD8^+^ T cells, previously characterized as tumor-reactive T cells^45^, were preferentially enriched at the tumor periphery (ROI3 and ROI4), potentially reflecting distinct immune responses to HBV infection versus tumor antigens. This detailed investigation underscores the spatial heterogeneity of HCC, offering insights that could significantly enhance our understanding of tumor microenvironment.

### Quasi-3D atlas exhibits microscale stromal heterogeneity in liver tumor

To comprehensive understand the tumor microenvironment in HBV-associated HCC, we extended our spatial analysis beyond traditional 2D projections to a quasi-3D reconstruction. By stacking PRISM data from 20 consecutive tissue sections using the same 31-gene panel, we generated a thick slab-like structure (Fig.5a). Single-cell transcriptomic analysis identified 1,218,279 cells across 32 clusters via our 3D PRISM approach, demonstrating high concordance with 2D PRISM annotations (Supplementary Fig. 30).

The 3D reconstruction unveiled architectural complexities obscured in 2D analyses. Tumor nests appearing isolated in 2D interconnected in 3D, highlighting the intricate spatial organization within the native tumor microenvironment (Supplementary Fig. 31). This spatial continuity also captured more realistic patterns of heterogeneity in cell distribution across the tissue (Supplementary Fig. 31). Immune cells were largely confined to non-tumor regions, with minimal infiltration into tumor cores (Supplementary Fig. 32). CAFs appeared to form physical barriers that impeded the entry of multiple types of immune cells (Fig. 5b, Supplementary Fig. 33), including CD8^+^ T cells and neutrophils, into the tumor core. Notably, NK cells were observed to traverse this CAF-defined boundary and accumulated at the tumor front, suggesting a distinct migratory behavior and potential antitumor roles (Fig. 5c).

**Fig. 5.**
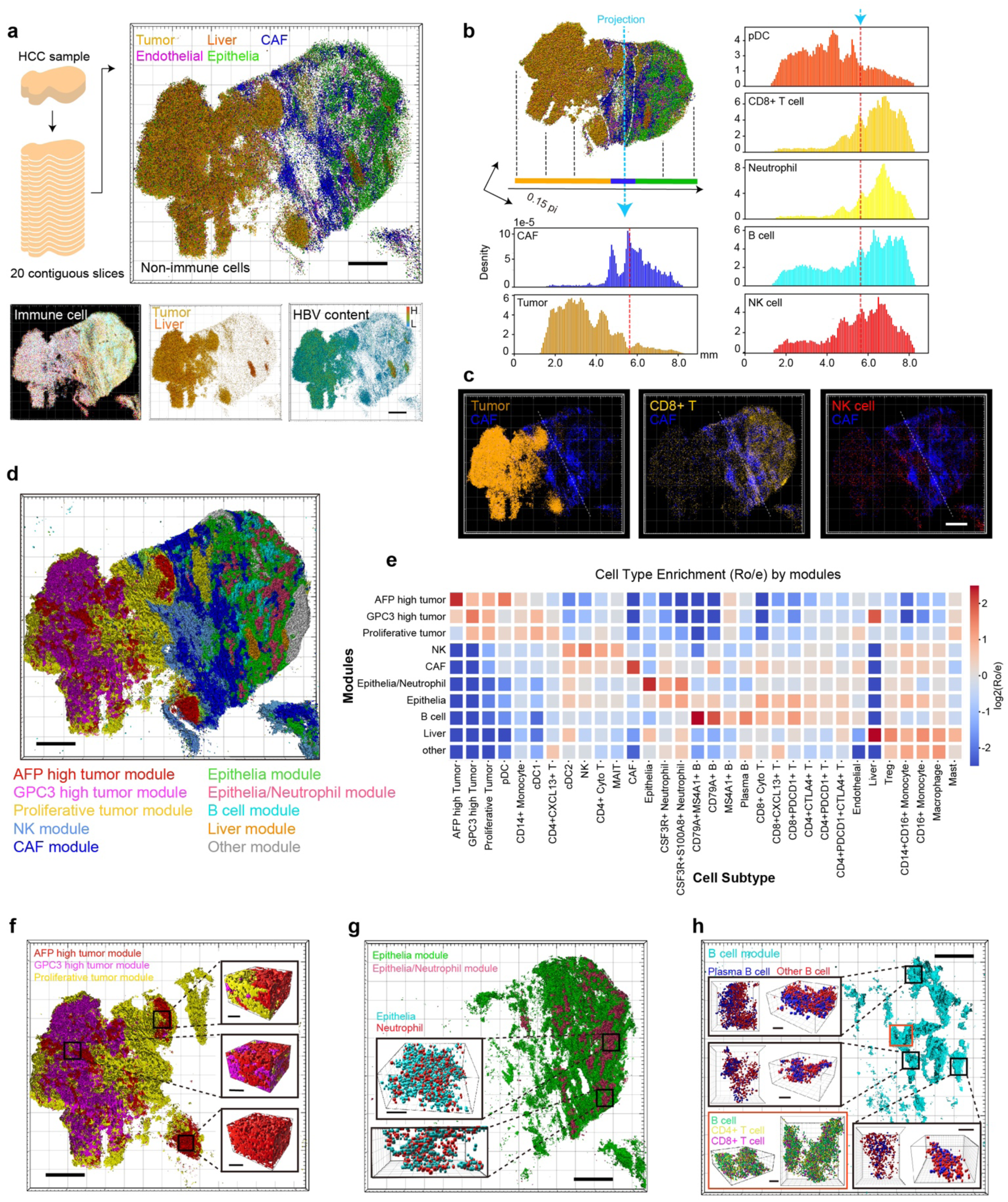
Quasi-3D atlas reconstitution of HCC microenvironment through PRISM consecutive profiling. **a**, Reconstituted 3D cellular atlas from 20 consecutive tissue sections. Scale bar: 1 mm. **b**, CAF barrier effect on immune cell infiltration through cell density projections. The sample was rotated 27° to horizontally align its ‘long-axis’. All cells were projected onto this long axis, and distribution histograms for selected cell types were plotted. The red dashed line indicates the peak distribution of CAFs. **c**, 3D projection of various types of cells. Scale bar: 1 mm. **d**, 3D projection of spatial domains annotated by STAGATE. Scale bar: 1 mm. **e**, Ro/e (ratios of observed cell numbers to random expectations) cell type enrichment analysis for STAGATE-defined regions shown in panel d. **f**, Intra-tumor heterogeneity analysis. Spatial distributions of GPC3-high, AFP-high and proliferative tumor cell modules exhibit distinct aggregation patterns. Scale bar: 1 mm (overview); 100 µm (zoom-in). **g**, Spatial projection of cell types in the Epithelia/Neutrophil module. Scale bar: 1 mm (overview); 100 µm (zoom-in). **h**, Spatial projection of cell types in the B cell-enriched module. Scale bar: 1 mm (overview); 100 µm (zoom-in).

To further interrogate the finer spatial architecture of the tumor microenvironment, we applied STAGATE^46^ analysis to our PRISM spatial transcriptomic data. Using a 40-μm neighborhood threshold to segment the tissue into spatial domains, we identified ten discrete cellular modules (Fig. 5d, e) with distinct cell compositions, including three tumor-enriched, one normal liver, one CAF, one epithelia, and three immune cell modules. 3D reconstruction revealed pronounced heterogeneity. Within tumor regions, distinct tumor-enriched modules interpenetrated in 3D space (Fig. 5f), reflecting complex clonal structures and invasion patterns. In non-tumor areas, we observed dense 3D niches of neutrophils co-localized with EPCAM^+^ epithelial cells (Fig. 5g), consistent with previous reports that EPCAM⁺ epithelial progenitors recruit neutrophils to facilitate tissue repair and modulate tumor invasion^47,48^.

Notably, structures that appeared as isolated B-cell clusters in 2D projections coalesced into interconnected lymphoid networks in 3D (Fig. 5h, Supplementary Fig. 31d). These B-cell-dominated niches harbored CD4⁺ and CD8⁺ T cells, with plasma cells enriched along tumor-oriented axes, consistent with previously reported chemokine-guided lymphocyte recruitment patterns in tumor-adjacent regions^49^. However, these aggregates lacked fully developed follicular dendritic cell networks or germinal center, resembling deviating tertiary lymphoid structures (TLSs) prevalent in anti-PD-1 non-responders^50^, implicating their roles in HCC progression. Taken together, these 3D spatial transcriptomic data provide a systematic and comprehensive view of the tumor’s internal architecture, with potential implications for understanding HBV-associated HCC.

### Large-scale 3D in-situ profiling unveils subcellular RNA distribution heterogeneities in intact thick tissues

We expanded the application of PRISM to analyze the intact 3D cell composition of 100-µm thick mouse brain tissue, demonstrating its inherent compatibility with the processing and imaging of thick tissues (Fig. 6a). This approach preserves the three-dimensional structure with high precision and accuracy. By employing single-round imaging, we avoid repetitive macromolecule penetration within thick tissues and effectively circumvent the challenges of fluorescent spot alignment and registration across multiple imaging sessions, which are common issues in existing multiplex RNA imaging techniques.

**Fig. 6.**
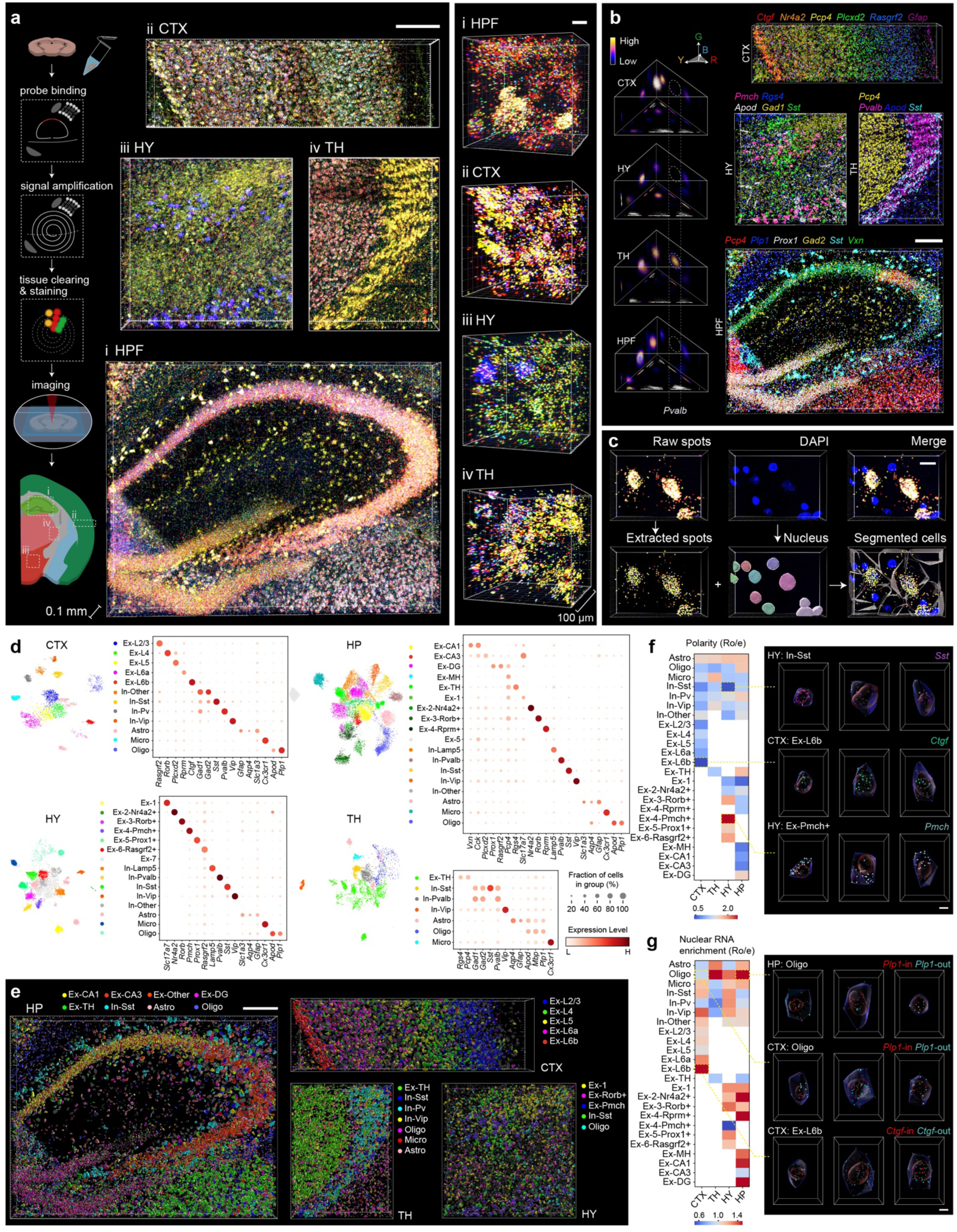
Three-dimensional RNA in-situ staining in intact mouse brain tissue. **a**, Workflow of PRISM 3D staining and the result. A liquid removal step is added after rolling circle amplification. RI matching was performed after imaging probe staining to enhance optical transparency. Resulting images depict tissues from four brain regions (HPF: hippocampus formation, CTX: cortex, HY: hypothalamus, and TH: thalamus). The images are composited using four channels (488nm, 543nm, 594nm and 647nm). Scale bar: 200 µm, upper left; 10µm, right. **b**, Gene calling in color space for four brain regions. The distribution of selected called genes in the four regions is shown on the right side. Scale bar: 200 µm. **c**, Three-dimensional cell segmentation. Identified gene spots were assigned to nearest nucleus centroid through Euclidean distance. Scale bar: 15 µm. **d**, Cell classification by Harmony embedding and Leiden clustering. The color of cell type is unified in UMAP plot and dot plot. **e**, Projection of selected cell type in the four brain regions. Scale bar: 200 µm. **f**, Subcellular RNA polarity analysis across brain regions and cell types. The polarity analysis was based on the scaled distance of RNAs’ centroid and nucleus centroid within one cell. The top 20% was designated as high polarity, and the bottom 20% as low polarity. The values in heatmap were calculated through Ro/e analysis using the ratio of high to low polarity. Polarity values from 30 genes were averaged in each cell type. On the right: examples of cells with different RNA polarity level. Scale bar: 5 µm. **g**, Nuclear RNA enrichment analysis using each cell’s Ro/e, calculated from the nuclear RNA fraction, as the indicator. On the right: examples of cells with different levels of nuclear RNA enrichment. Scale bar: 5 µm.

To improve optical transmission through thick tissues, we incorporated a post-amplification lipid-removal step and used iohexol for refractive index matching to ensure a deep light penetration (Supplementary Fig. 34). PRISM decoding process used signal ratios rather than absolute intensities, offering greater tolerance to optical attenuation (Extended Data Fig. 6). Utilizing a 30-gene panel, we profiled large-scale ROIs within the brain, including the cortex (1330 µm x 400 µm x 100 µm, 207,115 RNA transcripts), hypothalamus (770 µm x 770 µm x 100 µm, 214,671 RNAs), thalamus (585 µm x 775 µm x 100 µm, 156,368 RNAs), and hippocampus (1520 µm x 1150 µm x 100 µm, 548,220 RNAs) (Fig. 6a). The detailed morphology and distribution of marker genes within these areas align with those observed in 2D experiments, indicating consistency across dimensions (Fig. 6b).

Cell segmentation and classification within each ROI were performed using StarDist^51,52^ for 3D nucleus segmentation(Fig. 6c) (resulting in 4,792 cells in the cortex ROI, 9,838 cells in the hypothalamus ROI, 5,370 cells in the thalamus ROI, and 20,721 cells in the hippocampus ROI, respectively) and Harmony embedding coupled with Leiden cell clustering, respectively, allowing for the differentiation of glial cells and neurons into subtypes (Fig. 6d). These classifications show high correlation with single-cell transcriptome data, affirming the robustness of our approach (Supplementary Fig. 35). Furthermore, we projected the cells back to their original locations to construct a 3D neuron-architecture landscape (Fig. 6e).

We then characterized the subcellular RNA distribution (Supplementary Fig. 36). RNA polarity within each cell type was quantified by averaging the distances from RNA centroids to the corresponding cell centroid for each gene (Fig. 6f). This analysis revealed notable heterogeneity in RNA polarity among cells. For example, In-Sst and Ex-L6b cells exhibited minimal polarity with given genes, whereas Pmch^+^ cells in the hypothalamus displayed pronounced polarity, with most RNA gathering on one side of the cell body, indicative of unique cell orientations. These polarity patterns were consistent with previously reported in situ sequencing data^26^ (Supplementary Fig. 37a). Additionally, we assessed nuclear RNA enrichment across cell types and brain regions to examine subcellular transcript localization (Fig. 6g). Oligodendrocytes in the hippocampus and thalamus, and those particularly located within fiber tracts, demonstrated higher nuclear RNA enrichment compared to their cortical counterparts. This regional pattern was corroborated by RNAscope validation (Supplementary Fig. 37b-d). Together, these subcellular insights add another dimension to our understanding of cell type identity and state, further enriching the comprehensive profile provided by PRISM.

## Discussion

PRISM leverages multi-channel color barcoding, a strategy that uses specific spectral combinations to enhance information density within a single round of fluorescent in situ hybridization staining and imaging, while facilitating size-independent barcoding of clonal RCA products. PRISM’s coding space can be expanded by increasing the levels of division without the need to lengthen barcode sequences. This scalability is achieved by setting more divisions and reallocating each quantity of level. Each level can also be fine-tuned by adjusting the mixing ratio of fluorescent probes to non-fluorescent probes corresponding to each barcode segment in the padlock probes (Supplementary Fig. 38). This adjustability of the fluorophore proportion for each level and channel also allows for optimal differentiation in color space upon decoding, making PRISM adaptable to various optical imaging systems regardless of their specific configurations and performance characteristics.

Furthermore, considering the inherent autofluorescence of many tissues, PRISM’s design accommodates quantization of each spectral channel to different levels to ensure distinguishability within a fluorescence-background environment (Supplementary Fig. 39). We demonstrated its effectiveness in diverse sample types—including mouse brain, mouse embryo, and human tumor sections—and the ability to detect viral genome fragments as short as 28 bp, highlighting PRISM’s broad applicability.

With PRISM, encoding capacity is enhanced by creating higher intensity levels within respective channels, achieving an experimentally verified encoding capability of up to 64 genes in a single imaging round using four channels (Extended Data Fig. 7). Specifically, three spectral channels Ch1, Ch2, and Ch3, were quantized into fifths (0, 1/5, 2/5, 3/5, 4/5, 5/5), and the channel (Ch4) was divided into three levels: high (2), low (1), and none (0). This capacity allows for detailed spatial mapping, as demonstrated in an E14.5 stage mouse embryo, mouse brain and 100-µm thick tissue (Extended Data Fig. 8, Supplementary Figs. 40, 41). The decoding accuracy in a 64-plex PRISM experiment was evaluated to be 95.8% and 97.6% in mouse brain and embryo tissue, respectively (Extended Data Figs. 9, 10). With its ability to resolve dozens of identities in a single shot, PRISM offers a much higher information entropy of imaging than other multi-cycle methods, which are typically limited to only four identities. While cyclic strip-and-rehybridization methods or adding more optical channels could theoretically further increase multiplexity for PRISM, they would necessitate extensive instrumentation and image processing, compromising the ease of use. Challenges such as signal crowdedness, another common issue in amplification-based RNA in-situ detection technologies, are acknowledged as multiplexity increases. This could be addressed through expansion microscopy or introducing pseudo-channels via light-activable/switchable fluorescent dyes or fluorescence resonance energy transfer (FRET). Moreover, although PRISM demonstrates high in situ specificity, its sensitivity (about 10%) is notably lower than that of smFISH^53^, primarily due to the use of RCA-based amplification. This limitation aligns with previous findings^54^ and likely arises from suboptimal binding efficiency of single probes^55^, and this constraint may be alleviated by increasing the number of probes per target gene.

PRISM distinguishes itself through cost efficiency, simplicity, and rapid experimental turnaround. Using standardized reagents and conventional benchtop setups, it serves as a valuable tool for diverse applications, from molecular pathology to cell atlas construction, while minimizing reliance on specialized equipment and extensive resources. With an open-source design, PRISM aims to simplify high-multiplex RNA imaging, making it as accessible as standard immunostaining techniques.

## Acknowledgments

The authors thank Dr. Ye Liang of the State Key Laboratory of Membrane Biology and Dr. Liqin Fu of the School of Life Sciences and National Biomedical Imaging Center at Peking University for assistance with confocal microscopy imaging. The authors thank Lei Wang for experimental assistance. This work was supported by grants from National Natural Science Foundation of China Grants T2188102 (Y.H.), T2225005 (J.W.), Beijing National Laboratory for Molecular Sciences BNLMS-CXTD-202401 (Y.H.), Noncommunicable Chronic Diseases-National Science and Technology Major Project 2023ZD0520400 (C.Z.), Beijing Municipal Science and Technology Commission Grant Z221100007022003 (Y.H.), Ministry of Science and Technology Grant 2018YFA0800200 (J.W.), and Clinical Medicine Plus X - Young Scholars Project of Peking University, the Fundamental Research Funds for the Central Universities (C.Z.).

## Author contributions

Conceptualization: T.C., M.J. and Y.H. Experiment: T.C., S.Z., K.D., Z.L., M.T., Y.Z., C.Y., W.F., G.W., D.L., J.P., Y.P., P.F. and J.W. Data analysis: T.C., M.T., Y.Z., W.H., Z.L., S.Z., K.D., C.Z. and Y.H. Writing: T.C., S.Z., K.D., Z.L., M.T., C.Z. and Y.H.

## Materials and Methods

### Cell culture

HEK293T cells (ATCC) were cultured in Dulbecco’s Modified Eagle Medium (DMEM; Gibco, Cat. No. 11965092) supplemented with 10% fetal bovine serum (FBS; Gibco, Cat. No. 16000044) and maintained at 37°C in a humidified 5% CO₂ incubator.

### Mice

Animals were housed at 22-24 °C in 40%-60% relative humidity under a 12-h light/12-h dark cycle. Animal handling and tissue harvesting methods were conducted in strict adherence to local animal welfare laws and approved by the Laboratory Animal Care and Use Committee at Peking University, ensuring alignment with recognized ethical standards for laboratory research. Wild-type C57BL/6J mice, aged over 8 weeks (sex information was not related to our study), were euthanized to collect brain tissue, while embryos at 12.5-, 13.5-, and 14.5-days post-fertilization were also harvested. The mouse brains and embryos were embedded in Tissue-Tek O.C.T. Compound (Sakura), snap frozen, and stored at-80 °C. The frozen tissues were sectioned into 10-μm-thick sections using a CM1950 Cryostat (Leica) and affixed to a Superfrost Plus glass slide (Epredia). The mounted sections can be stored for one year at-80 °C.

For 100-µm-thick tissue collection, C57BL/6J mice (> 8 weeks) were perfused with pre-chilled 4% (w/v) RNase-free paraformaldehyde (PFA, Shanghai Yuanye). Subsequent to the brain harvest, the samples were immersed in 4% PFA for overnight fixation at 4 °C. The brain was sectioned into 100-µm slices using a VT1200 Vibratome (Leica). These slices were preserved in 70% ethanol at 4 °C and can be stored for more than 3 months.

### Human

A human hepatocellular carcinoma sample was placed in RPMI 1640 medium after being obtained from the patient and then processed following the aforementioned protocol (O.C.T embedded, snap frozen and sectioned into 10-μm-thick sections). This study was approved by the Ethics Committee of Beijing Shijitan Hospital, Capital Medical University. The patient in this study provided written informed consent for sample collection and data analyses.

### Probe and barcode design

For each gene of interest, a 40-nucleotide (nt) sequence window was identified to serve as a binding target, with 2 binding targets designated for mouse brain and embryo samples and 3 for human hepatocellular carcinoma samples. The selection of these 40 nt sequences adhered to stringent criteria: a melting temperature (Tm) in the range of 65-75 °C, the absence of 5 consecutive guanine (G) bases, coverage of all transcript variants (if possible), no blast results matching other genes in the NCBI database and no significant DNA secondary structure (ΔG >-10). Each 40nt sequence was divided into two segments of 20nt each for the construction of a padlock probe.

A “selective amplification” strategy was employed to mitigate signal crowdedness issues^26^. Briefly, padlock probes targeting highly expressed genes were doped with irreplicable ones at proper ratios during probe pooling. This approach can dilute the clonal populations by partially suppressing the amplification of abundant transcripts. During analysis in a gene-by-count expression matrix, the quantities of those transcripts can be recalibrated by multiplying respective dilution ratios. The dilution ratios we used for all samples were listed in **Supplementary Table 1**.

Each gene was assigned by a specific barcode (Fig. 1a,b). The barcode consists of four segments (20nt for each), with segments specifically binding to fluorescent probes labeled with Texas Red (TR), Alexa Fluor 488 (AF488), Cy5, and Cy3, respectively. For 30-or 31-plex coding, each segment variation allowed for 0-4 intensity levels per channel except for Cy3, which was compromised to two levels due to autofluorescence background. Color Intensity grading is realized by mixing fluorophore-labeled probes with unlabeled probes in a specific ratio. Mixing ratio for all imaging probes was listed in **Supplementary Table 2**. The ratio for each probe pair can be finetuned based on actual optical setups. All padlock probes and imaging probes were ordered from Sangon Biotech (Shanghai).

### Sample treatment for thin tissue

For a 10-µm-thick frozen tissue sections, they were fixed in 4% PFA with 0.1% Glutaric dialdehyde at room temperature for 15 min after taking out from-80 °C refrigerator, and then rinsed with PBST (0.05% Tween in 1X PBS). The sample was then permeabilized with 0.01% pepsin in 0.1M HCl at 37 °C for 2 min (10 min for mouse embryos), followed by PBST washing. After that, the samples were dehydrated in a series of ethanol washes (10 min in 80% ethanol followed by 2 min in pure ethanol) and rehydrated with three-time PBST washes. The samples were then blocked with oligo-dT (100 nM oligo-dT [Sequence: AAGCAGTGGTATCAACGCAGAGTACT_30_VN], 50 mM KCl, 20% formamide, 200 μg/mL BSA, 200 μg/mL Yeast tRNA (AM7119, Invitrogen), and 1U/μL Ribolock RNase inhibitor (Thermo Scientific) in Ampligase buffer (Lucigen)) for 10 min at room temperature, thereby suppressing non-specific probe binding. The probes were hybridized to sample by incubating them in a hybridization mix (200 nM padlock probe for each, 50 mM KCl, 20% formamide, 200 μg/mL BSA, 200 μg/mL Yeast tRNA, and 1U/μL Ribolock RNase inhibitor in Ampligase buffer) at 55 °C for 15 min and then 45 °C for 2 h. The samples were placed in a humidified chamber during incubation. After hybridization, the samples underwent three 10-min washes with washing buffer (10% formamide, 2X SSC buffer) to remove non-specifically bound probes, then rinsed twice with PBST. The samples were then incubated in a ligation mix (2.5 U/μL SplintR ligase (NEB), 200 μg/mL BSA, and 1U/μL Ribolock RNase inhibitor in SplintR buffer) at 37 °C for 2 h in the humidified chamber to ligate the nick on the padlock probe, and then the samples were rinsed twice with PBST. Then rolling circle amplification (RCA) was performed using 0.25 U/µl Phi29 polymerase in Phi29 polymerase buffer (Thermo Scientific) with 250 μM dNTP (Thermo Scientific), 50 μM aminoallyl-dUTP (Thermo Scientific), 10% glycerol, 200 μg/mL BSA, and 300 nM RCA primer at 30 °C overnight. Following RCA, the samples were rinsed twice with PBST, then fixed with 10 μg/μL BS(PEG)_9_ (Thermo Scientific) in PBST at room temperature for 15 min, and subsequently rinsed three times with PBST. Finally, the samples were washed three times (5 min per wash) with 65% formamide at room temperature, followed by two PBST rinses.

### Sample treatment for thick tissue

Tissue slices were rehydrated through three 20-min incubations in PBST after taking out from ethanol, then blocked with oligo-dT (the reagent was the same as mentioned above) for 30 min at room temperature. After that, the probes were hybridized to samples as mentioned above at 45 °C for 24 h with gentle shaking, followed by washing three times for 20 min each in washing buffer, then three times in PBST, then overnight incubation at 37 °C in ligation mix as mentioned above with gentle shaking. After two PBST washes, RCA was performed at 30 °C for 12-24 h, followed by two gentle PBST washes and a 30-min fixation in BS(PEG)_9_ as mentioned above. After being washed three times with PBST and 65% formamide at room temperature, the samples were treated with Cubic L (TCI, T3740) for lipid removal at room temperature for 1 h, followed by three-time washes with PBST.

### Imaging probe staining

The samples were incubated with a mixture of 17 fluorescence-labeled probes (each paired with an unlabeled probe at a predetermined ratio, with a final concentration of 120 nM per pair, 20% formamide, 2X SSC buffer) at 50 °C for 9 min, then 37 °C for 21 min. Following incubation, the samples were washed three times with washing buffer, stained with DAPI (5 μg/μL, Beyotime) solution at room temperature for 5 min, and washed twice with PBST. Finally, the samples were mounted for imaging with a cover slip.

For staining 100-µm-thick samples, the imaging probe incubation was performed at 37 °C for 2 h, followed by three washes with washing buffer, a DAPI staining at room temperature for 30 min, and two PBST washes. The samples were then immersed in 0.9 M Iohexol (2X SSC buffer) for 1 h at room temperature to adjust the refractive index. Finally, the sample was mounted for imaging with a cover slip and sealed with varnish to avoid evaporation.

### Imaging

Imaging can be performed using any proper instrument. Our experiment used both home-built setups and conventional equipment in the core facilities. For our thin tissue imaging, the sample was placed on a home-built epifluorescence microscope, using MIM microscopic imaging framework (Applied Scientific Instrumentation: RAMM-Basic, RAMM-URKIT2, MIM4-OSM25, TN200-MMC, LLG3-E100, MS4, S551-2201B, C60-3WMS-Mx) equipped with an S551-2201B motorized stage (Applied Scientific Instrumentation), an ATF6.5 SYS 785 automated focusing module (Wise Device Inc.), a CFI S Plan Fluor ELWD (40X NA 0.60) objective lens (Nikon), an X-Cite Turbo LED light source (Excelitas Technologies), and an Orca Fusion BT scientific CMOS camera (Hamamatsu Photonics). To image a 10-µm thick tissue, we collected a z-stack of 9 planes at ∼1 µm intervals for each tile. 4% overlap was preserved for image registration and stitching. Cy3 / Cy5 (532/647 nm) and AF488 / TxRed / DAPI (488/594/405 nm) channels were serially imaged. The sampling rate was 0.1625 µm x 0.1625 µm per pixel. For a tissue section similar in size to a coronal section of a mouse brain (∼1 cm x 0.6 cm), approximately 600 tiles were imaged, and it took around 90 min for imaging. The imaging can also be performed with a commercialized fluorescent microscope or a slide scanner with appropriate channels. Conventional confocal microscopes, such as LSM880 and LSM980 (Zeiss), were used for thick tissue imaging. Fast DAPI scanning across the whole slice was performed to find regions of interest. 647 nm, 594 nm, 543 nm, 488 nm, and 405 nm were used for imaging. The sampling rate is 0.207 x 0.207 x 0.700 µm/voxel for a 100 µm-thick brain slice. The imaging software was Zen or Zen Blue for LSM880 and LSM980, respectively.

### Image processing

The thin tissue imaging processing was conducted as described in SPRINTseq^26^. Briefly, images from multiple planes were stacked along the z-axis through an all-in-focus algorithm, followed by shade correction with CIDRE^56^ and inter-channel image registration (Fast Fourier Transform and Maximum Cross-Correlation). We resized the image from different channels to correct color aberration by re-sampling the image tile-by-tile in respective channel (github: PRISM_Code/Image_process/image_process_after_stack.py/resize_batch). The tiles were stitched using the Microscopy Image Stitching Tool^57^. For the alignment of 20 consecutive sections of HCC, we manually identified and marked 12 anchor points on each slide. The initial slide served as the template, and affine transformations were applied to the subsequent 19 slides, utilizing the established anchor points. This procedure facilitated the creation of a comprehensive, albeit artificial, three-dimensional representation of the HCC sample. For thick tissue, registration from each channel and stitching from each tile were performed in ImarisStitcher (Oxford Instruments). Given that dehydration, refractive index (RI) matching during sample preparation can cause a slight tissue shrinkage in the z-direction. Z-length was rescaled to 100 µm during image processing.

### Gene decoding

Signal spots were extracted from all four channels, using local maxima on tophat-filtered images for thin tissue (Extended Data Fig. 2). 3D local maxima and Gaussian fitting (Airlocalize)^58^ were used for spot extraction for thick tissue. Spot signal coordinates across all channels were combined to get 4-dimensional (from Ch1 to Ch4) intensity information. Spectral crosstalk was corrected according to optical set up condition. In our imaging platform, we observed approximately 10% signal bleed-through from the Cy3 channel into AF488 channel. Thus, we subtracted corresponding proportion of the AF488 channel intensity from the Cy3 channel on a pixel-by-pixel basis. Four intensity values were normalized through the mean values of respective channels. Spots with low sum-intensity values were filtered out.

In barcode design, Ch1, Ch2, and Ch3 sum into a fixed value (for a 30/31-barcode set, this sum is 4), corresponding to 15 intersected barcodes (Fig. 1e). Correspondingly, all spots from real data made up 15 rays in color space (x=Ch1, y=Ch2, z=Ch3, Extended Data Fig. 2). Due to experimental noise, such as amplification variations and the fluorophores’ chemical environments, spots belonging to one ray may exhibit behavior indicative of a joint distribution. This joint distribution resembles a Poisson-like distribution along the direction of the ray and a Gaussian-like distribution in the direction perpendicular to the ray. For the ease of following analysis, we projected all spots onto the plane (x+y+z=4); this was accomplished by scaling the sum value of each spot to 1 (L1-normalization). Therefore, the values in each spot were transformed to their respective proportions: Ch1/(Ch1+Ch2+Ch3), Ch2/(Ch1+Ch2+Ch3), Ch3/(Ch1+Ch2+Ch3). Thus, in color space, the Poisson-like distribution along the ray can be omitted, and spots on the plane can be seen as the results of 15 Gaussian distributions with different means and variances or 15 clusters. The channel Ch4 was also scaled to Ch4/(Ch1+Ch2+Ch3), set as a new z-axis for the projected 2-D plane. The new color space axes were defined as x = (Ch1-Ch2)/(Ch1+Ch2+Ch3), y = 2*Ch3/(Ch1+Ch2+Ch3)-1, and z = Ch4/(Ch1+Ch2+Ch3). As a result, 15 clusters (determined by Ch1, Ch2, and Ch3) x Ch4 (Yes or No) make up a new color space containing 30 clusters. Barcode 31 (Ch4 only) was not in this color space, but it could be easily extracted since it has a high-scaled Ch4 value.

The position of each cluster in the color space corresponds to a specific barcode by design. Gaussian fitting was used to evaluate the degree of separation between clusters (Extended Data Fig. 3c). Since the barcode at endpoint “0” and “1” (*e.g.*, ‘4000’, ‘0040’) was not a cluster but a spot in color space, we introduced a small variance to coordinates from all spots to make the endpoint cluster a pseudo-Gaussian distribution. This way, an equal assessment can be achieved between these endpoint clusters and other clusters. The initial gene calling step for all signal spots was based on thresholding their confidence level of belonging to a certain cluster. For a refined gene calling confirmation, three-dimensional boundaries were manually delimited for barcodes in color space. Direct gene calling (all spots within such delimited boundary were assigned to a specific barcode) or iterative Gaussian fitting within delimited boundary before thresholding confidence level could be performed to ensure decoding accuracy (Fig. 1f, Extended Data Fig. 3c).

### Experimental validation

Signal detection in the PRISM workflow consists of two key steps: (1) rolling circle nanoball (amplicon) generation through padlock probes targeting mRNA and amplification, and (2) PRISM color decoding on nanoballs. The overall detection accuracy is jointly determined by the performance of both steps, and we validated independently through smFISH experiment and nanoball decoding verification experiment. We also benchmarked overall performance by comparing PRISM 30-plex spatial expression pattern with RNAscope on adjacent tissue sections. Beyond accuracy, smFISH and RNAscope experiments were also used to characterize amplification sensitivity and overall sensitivity. Together, these experiments-smFISH, RNAscope, and strip-rehybridization-based nanoball decoding verification-comprehensively confirmed the fidelity and sensitivity of the PRISM detection workflow.

1) smFISH (Stellaris RNA FISH). Three experiments (a-c) were performed in this manuscript, (a) and (b) were for amplification accuracy and (c) was for amplification sensitivity.

a) smFISH and RCA simultaneous staining and imaging on the same sample (Supplementary Fig. 3). Since smFISH signal is susceptible to RNA degradation, smFISH probe hybridization step was prioritized before RCA. However, it is reported that RCA can degrade smFISH fluorescence signals due to the 3’-5’ exonuclease activity of Phi29 polymerase^59^. Therefore, we adopted a two-stage smFISH probe, where the primary smFISH probe includes a single-stranded end to improve signal preservation during RCA^60^.

Fresh cultured cells were rinsed with PBS twice and then fixed in 4% PFA for 15 min at room temperature, and then rinsed with PBS again. Then the cells were permeabilized for 20 min at 4°C in PBSR (8 U/ml Ribolock RNase Inhibitor) with 0.1% Triton X-100. After three-time PBSR washes, the cells were further permeabilized in a graded ethanol series (30%-50%-70%-100%, 3 min for each step). After drying in 100% ethanol, the cells on glass slide are stored at-80°C refrigerator overnight. Then, cells were taken out from-80°C refrigerator and rehydrated with three-time PBSR washes. After incubating for 20 min at room temperature with prehybridization buffer A (30% formamide and 2X SSC buffer), the cells were stained overnight at 37°C with hybridization buffer consisting of 10% dextran sulfate, 30% formamide, 2X SSC, 1U/μL Ribolock RNase inhibitor, 200 μg/mL Yeast tRNA and 100 nM primary probe. After that, the cells were washed using prehybridization buffer A for 20 min at 37°C for three times, and then were post-fixed in 4% PFA for 15 min at room temperature, with three-time PBSR washes.

After smFISH primary probe hybridization, PRISM amplification steps (including blocking, padlock hybridization, ligation, RCA and post-fix) were performed as mentioned above. Then the cells were stained with 10% dextran sulfate, 10% formamide, 2X SSC, 1U/μL Ribolock RNase inhibitor, 200 μg/mL Yeast tRNA and 10 nM smFISH secondary probe (TxRed-labeled) and 10 nM RCP imaging probe (AF488-labeled) for 3 h at 37°C. The cells were then washed three-time for 10min in prehybridization buffer B, followed by three-time washes in PBSR.

Finally, the cells were mounted in Anti-Fade Mounting Medium (Sangon, E675011-0010) and imaged with confocal microscopy (LSM 880). The result showed that more than 90% of RCP signal is co-localized with smFISH signal (Supplementary Fig. 3). In this experiment, we observed an inverse correlation between RCA and smFISH signal intensity, suggesting a certain degree of potential physical mutual exclusivity from two detection. While these co-localization results can be used to validate the specificity and accuracy of amplification (> 90%, the rest 10% includes RNA degradation, primary smFISH probe degradation and RCA false detection) this experiment cannot be used to assess amplification sensitivity due to the exclusivity.

b) Serial detection using Stellaris smFISH and RCA on the same sample (Supplementary Fig. 4). smFISH experiment was also prioritized before RCA since it is susceptible to RNA degradation.

In this experiment, we first used fluorophore-labeled smFISH probe to directly hybridize sample and perform imaging, after which we strip the smFISH probe and then perform ‘Padlock & RCA’ (PRISM amplification) as well as imaging probes staining and imaging.

In detail, after two-time PBS rinsing, fresh cultured cells were fixed in 4% PFA at room temperature for 15 min, and then rinsed with PBS. Then the cells were permeabilized for 20 min at 4°C in PBSR (8 U/ml Ribolock RNase Inhibitor) with 0.1% Triton X-100. After three-time PBSR washes, the cells were further permeabilized in a series of ethanol washes (30%-50%-70%-100%, 3 min for each step). After drying in 100% ethanol, the cells on glass slide are put in-80°C refrigerator for stock. The cells were taken out from-80°C refrigerator and rehydrated with three-time PBSR washes. After incubating for 20 min at room temperature with prehybridization buffer A (30% formamide and 2X SSC buffer), the cells were stained overnight at 37°C with hybridization buffer consisting of 10% dextran sulfate, 30% formamide, 2X SSC, 1U/μL Ribolock RNase inhibitor, 200 μg/mL Yeast tRNA and 10 nM TxRed-labeled smFISH probe. Then the cells were washed using prehybridization buffer A for 20 min at 37°C for three times, followed by three-time PBSR washes.

The cells were mounted in Anti-Fade Mounting Medium and imaged using wide-field microscopy with 60X objective lens. We collected a z-stack of 8-9 planes at ∼1 µm intervals and used all-in-focus algorithm to achieve focal stack as described in ‘imaging process’ section. The field-of-view positions were recorded.

After acquiring smFISH signals, the coverslip on cells was unmounted and mounting medium was washed away using PBSR. Then, the smFISH probe was washed away in 60% formamide at 50°C for 10 min, repeated for three times and then washed with PBSR for three times. Then, PRISM amplification steps (including blocking, padlock hybridization, ligation, RCA and post-fix) were performed as described above. After staining with imaging probes for amplification product, the cells were mounted again in Anti-Fade Mounting Medium. Imaging was performed with the same condition and at the same positions as recorded during 1st smFISH-imaging round, and RCA signal at these positions were acquired.

Signals from smFISH and RCA are registered during analysis. As the result in Supplementary Fig. 4 shows, more than 95% of RCA signals are co-localized (position shift < 0.3 µm) with smFISH signals. The sub-micron position displacement between two signals is mainly due to two reasons: the RCP has a typical size around hundreds of nanometers, and the inter-steps stripping.

Notably, both experiment (a) and (b) demonstrate that ‘Padlock + RCA’ signal amplification exhibits high accuracy and specificity, but neither of these two colocalization experiments can be used to characterize sensitivity (detection efficiency) of RCA. Mutually exclusive pattern was also observed in experiment (b), where smFISH signals were dense and strong, RCA signals were notably weak or sparse. This is particularly pronounced in the nucleus, where RCA signals become nearly undetectable. Such exclusion may be attributed to interference from the co-localization experiment workflow such as steric hindrance, RNA degradation, or buffer incompatibility. To fairly evaluate sensitivity of RCA in a standalone condition, we performed the experiment (c).

c) Parallel smFISH and PRISM tests to evaluate detection sensitivity (Supplementary Fig. 5). In this experiment, we performed smFISH and PRISM on different sample and directly compared the mRNA count. The cell culture condition and pre-treatment procedures were kept consistent in two groups.

For smFISH-group, the workflow was consistent with (b) until the completion of smFISH-imaging. For RCA-group, the workflow was consistent with direct PRISM experiment from PFA fixation to imaging.

Cells with similar sizes were selected for mRNA quantification during analysis. Results showed that the overall sensitivity of RCA (one RCA probe) is around 21% (Supplementary Fig. 5). It should be noted that the sensitivity may vary depending on multiple factors, including padlock probe numbers, target gene, and sample type.

2) RNAscope^TM^ (ACD). To systematically benchmark PRISM overall accuracy and sensitivity in tissues, 30-plex PRISM (one RCA probe per gene) were benchmarked with RNAscope. We used one brain section to perform PRISM and its adjacent three sections to perform RNAscope for three different genes (*Snap25*, *Slc17a7* and *Apod*). The probes and reagents were ordered from ACD, and experiment was performed strictly according to ACD protocols (UM 323100), with one gene per section. Imaging procedure followed the wide-field microscopy conditions as mentioned above. Signal extraction was also performed as described in ‘Gene decoding’ section (tophat and local maximum). Sensitivity for the three selected genes was quantified by comparing signal count ratios between PRISM and RNAscope, yielding average values of 9.4% (total gene counts), 15.8% (median per cell with non-zero expression), or 11.5% (mean per cell with non-zero expression).

3) Nanoball decoding verification. While the amplification accuracy (corresponding to mRNA-to-nanoball process) has been validated by co-localization experiment (1-a) and (1-b), the nanoball decoding accuracy (corresponding to PRISM color-coding) was assessed through a strip-rehybridization based nanoball-specific FISH experiment following a PRISM experiment.

The experiment began with either a 30-plex or 64-plex PRISM assay. Tissue sections were assembled into a flow cell by stacking the slide, a strip of double-sided adhesive tape (20106, ARcare) with a cut-opening to form a flow channel, and a blank slide containing two holes for in-and-out reagent flow, as we described in the SPRINTseq protocol^26^.

All post-imaging reactions were performed within the flow cell to facilitate accurate image registration. After PRISM imaging, the imaging probes were stripped away using 60% formamide at 50°C.

To obtain the ground truth nanoball positions from a specific gene, we designed a fluorescently-labeled nanoball-check probe targeting the mRNA-binding region on padlock probe, which is also amplified along with the barcode region during RCA. These nanoballs of a specific gene were selectively stained using corresponding nanoball-check probe (50 nM, in 2X SSC and 20% formamide), generating nanoball-check image for that gene (Extended Data Fig. 5a). This process was performed iteratively for different genes. Prior to hybridizing a new nanoball-check probe, the previous check probe (already imaged) was stripped away using 60% formamide at 50°C.

We then aligned the PRISM decoded coordinates for each gene with its corresponding gene’s nanoball-check image. Fluorescent intensity value was obtained by reading the nanoball-check image using coordinates decoded by PRISM. To eliminate the potential registration-induced bias on accuracy calculation, we selected fields of view with perfect inter-round registration for such intensity reading.

The frequency distribution of the intensity was plotted to characterize the decoding accuracy (Extended Data Fig. 5b). Nanoballs correctly decoded in the PRISM assay will be fluorescently stained in the check-image (overlapping part), showing high fluorescence intensity and forming a right-skewed Gaussian-like peak (gaussian like: since the size of nanoballs is heterogeneous). In contrast, misidentified coordinates exhibited low intensity (non-overlapping), forming a sharp left peak. The proportion of “non-overlapping” coordinates represented the false decoding rate. Different barcodes / genes were used for such decoding accuracy calculation. For the 30-plex PRISM assay on mouse brain tissue, the average false decoding rate was around 4.33%.

Further validation of decoding accuracy on nanoball was performed on 64-plex PRISM datasets from mouse brain and embryo tissues, yielding false decoding rates of 4.20% and 2.40% respectively, suggesting robust gene-calling performance for 64-plex multiplexity (Extended Data Figs. 9, 10).

Notably, we found that some barcodes may be prone to false calling: barcodes with only one channel signal in the first three channel (‘single color’, such as 0040: 5.85%, and 0052: 7.20%) showed increased susceptibility to cross-talk from neighboring barcodes. We attribute this to minor color aberrations or imperfect signal registration that may cause a small fraction of ‘spot splitting’. As such, we recommend avoiding the use of single-color barcodes for critical targets, particularly when imaging or signal extraction conditions are suboptimal.

### Cell segmentation

Cell segmentation and RNA assignment for thin tissue were performed as described in SPRINTseq^26^ (Supplementary Fig. 10, Fig. 2b). Briefly, nuclei were segmented using adaptive thresholding of DAPI-stained images to generate binary images. The binary image underwent Euclidean transform and was further segmented using the watershed algorithm. Identified RNA spots were assigned to the nearest nucleus centroid using a K-D tree, with a “Cell Index” assigned to each RNA in the result table to create an expression matrix for the cells. For 3D data from thick tissue, nuclei segmentation was achieved using the StarDist algorithm with a pre-trained model.

### Annotating and mapping cell types

Different cell-type classification methods were applied. For data derived from mouse embryos, due to the large number of cell types and the lack of comprehensive reference single-cell transcriptome datasets, cell types were directly annotated based on marker gene expression levels. For other datasets, we primarily used Harmony^61^ algorithm to integrate our data with published single cell transcriptome datasets. After integration, a graph-based structure was created, enabling the division of cells into multiple clusters using Leiden algorithm at high resolution, followed by manual annotation of cell type based on marker genes. This process was applied to the 2D mouse brain section (Supplementary Fig. 11), 3D mouse brain data (Fig. 6d), and Human Hepatocellular Carcinoma (HCC) samples (Fig. 4c, Supplementary Fig. 30). Each cell’s boundary was delineated by constructing a convex hull based on its peripheral RNA spot. Cells were visualized in 2D or 3D using uniform manifold approximation and projection, with colors assigned to each cell type.

For the 2D coronal section of the mouse brain, single-cell transcriptome datasets from mousebrain.org for four tissues (CTX: cortex, HP: hippocampus, TH: thalamus, HY: hypothalamus) were integrated with our data. After preprocessing, PCA, neighbor detection, Leiden clustering, and annotation, the cells were divided into 14 types, including excitatory neurons, inhibitory neurons, and glial cells (Supplementary Fig. 11).

For 3D mouse brain, the cell classification was performed by 4 tissue types respectively, and the strategy was similar. Different thresholds were used based on the differences between tissues (CTX, HP, TH, HY). The integration of single-cell datasets for these tissues, obtained from mousebrain.org, was conducted using the Harmony algorithm. Following integration, a graph-based structure was created, enabling the division of cells into multiple clusters using the Leiden algorithm at a high resolution and then manually annotated as various cell types and subtypes based on marker genes and tissue type (Fig. 6d). Detailed parameters are shown in the corresponding Jupyter Notebook.

For HCC, the data passed quality control was processed standard pipeline, including normalization, log1p, regress_out, and scaling, before performing Principal Component Analysis (PCA). This foundational work facilitated the integration of spatial transcriptomics data with three distinct single-cell transcriptome datasets (GSE151530, GSE140228, CNP0000650) using the Harmony algorithm within the Scanpy framework. Utilizing high-resolution Leiden shared nearest neighbor clustering. We identified a fine-grained set of subclusters, which were meticulously annotated based on the expression of numerous housekeeping and marker genes associated with HCC, revealing 35 subclusters representing a diverse array of immune and nonimmune cell types. Notably, the challenge posed by the absence of well-defined marker genes for liver cells was addressed by leveraging elevated Hepatitis B Virus (HBV) expression as a proxy, allowing for the precise annotation of liver cell types. This approach helped to reclassify cells initially tagged as ‘other’ types, ultimately enriching the final dataset with 60,329 cells categorized into 17 major types and 35 subtypes (Fig. 4d, Supplementary Fig. 25). Each cell type was visually distinguished using a unique color scheme in the convex hull representation, with the dataset displayed in two dimensions using Uniform Manifold Approximation and Projection (UMAP)^62^.

A similar methodology was applied to analyze the quasi-3D HCC dataset, yielding annotations that identified the same major cell types while revealing slight variations in cell subtypes.

### Cell classification evaluation using single-cell transcriptome data

To verify the accuracy of cell classification, we subjected the single-cell data from the mouse brain cortex, comprising 50,478 cells with 27,998 genes from mousebrain.org, to Leiden clustering. We manually annotated these cells based on the expression of the 30 marker genes we were interested in. Subsequently, we took the expression matrices processed by normalization, log1p, regress_out, and scale of corresponding cell types from both spatial transcriptomics and single-cell transcriptome data, and then averaged the data across the cells within each cell type. This process transformed the gene expression of each cell type into a 1×30 vector. We then conducted a Pearson correlation analysis on the representative vectors of these nine cell types, obtaining a correlation between PRISM data and single-cell transcriptome data (Fig. 4c, Supplementary Fig. 30b, Supplementary Fig. 35a).

### Spatial analysis in 2D mouse embryos and brains

Spatial-related quantitative analysis in mouse embryos involved coarse-grained spatial correlation and intracellular interaction analyses. For coarse-grained spatial correlation analysis, annotated cells were spatially sub-sampled to proper bin size (∼ 200 µm x 200 µm). Pearson correlation was calculated across each bin and cell type to analyze the co-occurrence between cell types (Fig. 2g). For intracellular interaction analysis, the nearest neighbor cell for each cell was first identified through the KD-Tree search, and such neighbor cell pair counts between any two cell types were calculated. Each pair was normalized by dividing the total cell numbers in each cell type, eliminating the bias caused by the total cell count difference (*e.g.*, muscle_neuron_ / muscle, muscle_neuron_ refers to the number of muscle cells whose nearest neighbour cell are neurons) (Fig. 2h). This approach can reveal the interaction between cell types from a direct-interaction perspective.

### Spatial analysis in 2D HCC

Spatial-related quantitative analysis in HCC tissue included three major parts: spatial domain segmentation using GASTON^44^, region-specific cell type diversity quantification, and neighborhood interaction analysis.

Tissue segmentation was performed using GASTON, an algorithm that integrates transcriptomic and spatial information. Dimensionality reduction of the gene expression matrix was conducted using Generalized Linear Model Principal Component Analysis (GLM-PCA) under a Poisson noise model, retaining the top 30 principal components. These GLM-PCs, combined with spatial coordinates, were used to train a dual-component neural network consisting of an isodepth network, which maps spatial coordinates to a continuous scalar isodepth value, and an expression reconstruction network, which predicts the GLM-PCs from the isodepth value. To ensure robustness, the model was initialized and trained 30 times with different random seeds.

After training, isodepth values were discretized into spatial domains using dynamic programming. The optimal number of domains was determined by identifying the knee point in the log-likelihood curve, resulting in six spatial domains in our analysis. These domains group tissue regions based on transcriptional similarity, not physical proximity, and are thus not necessarily spatially contiguous. This approach enables the identification of similar biological microenvironments that may occur in spatially disconnected regions of the tissue. Final visualization included both continuous isodepth contour plots and discrete spatial domain maps, providing complementary views of tissue structure. Cell subtype relationships within the tumor microenvironment were subsequently inferred from the spatial transcriptomics data.

From the GASTON-defined domains, five representative Regions of Interest (ROIs) were manually selected for in-depth analysis. These ROIs were chosen to represent archetypal, spatially contiguous regions within each domains and were guided by the isodepth map to ensure they reflected high-confidence transcriptional signatures. To assess cellular diversity, a Chi-squared test was performed across these ROIs. Cell type enrichment, particularly among immune populations, was quantified using observed-to-expected cell ratios (Ro/e)^41,63^. Spatial relationships between cell subtypes were then further examined within each ROI using the spatial transcriptomics dataset.

Spatial adjacency relationships were defined using Delaunay triangulation, as implemented in the Squidpy framework^64^. Cell-cell interactions were quantified through identified adjacency frequencies matrices, normalized by subtype abundance to correct for compositional bias. A significance threshold (θ = 0.06) was applied to identify biologically relevant interactions. The filtered interaction matrix was represented as a directed, weighted graph using igraph library^65,66^, where nodes represent cell subtypes (n=32) and edges denote significant spatial interactions, weighted by normalized interaction frequency. Visualization employed a force-directed Kamada-Kawai layout, with node color indicating lineage (e.g., T cells, B cells, myeloid cells), and edge width scaled logarithmically (log₁₀(W)) to emphasize strong interactions. For cell type-specific analyses (e.g., AFP-high tumor cells), interaction profiles were extracted, transformed using log₁₀(x+1) to enhance dynamic range, and visualized via ranked bar plots, distinguishing encirhed interactions from background.

### Spatial analysis in 3D Stacked HCC Data

For 3D HCC data, we extended spatial analysis to account for volumetric tissue structure, including the assessment of neighborhood enrichment, cell type distribution evaluation with respect to specific orientation, and spatial functional modules defined by STAGATE^67^.

Neighborhood enrichment was computed using Squidpy^64^. A spatial neihbor graph was constructed from 3D cell centroid coordinates using default parameters. Enrichment scores were calculated via a 1000-round permutation test using the ‘gr.nhood_enrichment’ function and through heatmaps generated by ‘pl.nhood_enrichment’ function. To assess directional distribution of specific cell types (e.g., to examine the ‘barrier effect’ of CAF), cells were projected onto a reference line based on their centroids, and density distributions were computed (Fig. 5f).

We employed STAGATE for domain segmentation in stacked slices. Spatial neighborhood graphs of continuous slices were generated with radial distance cutoffs (8-65 μm) to define cellular interactions. A graph autoencoders was trained to embed spatial and transcriptomic features into a low-dimensional latent space. Based on UMAP visualization of embeddings, a cutoff of 40 μm was selected as optimal for balancing spatial continuity with resolution. Subsequent multi-resolution clustering yielded 10 biologically meaningful spatial modules. Each domain was characterized by its cell type composition, enrichment ratio (Ro/e), and spatial distribution, revealing distinct microenvironmental niches in HCC tissues.

### Subcellular analysis

Given the coordinates of each RNA spot, the nucleus centroid, and the nucleus region, we profiled the nuclear RNA enrichment level, along with subcellular polarity, which exhibited certain specificities on the tissue or cell type. The chi-square test was used to analyze the relation of different variables. The variables in our experiment are RNA distribution and tissue or cell type. To further represent the direction of the relation, we calculate the Ratio of Observed to Expected (Ro/e) of annotated RNA molecules. This allowed us to observe under which classification conditions RNA “clusters”, thereby determining its distribution characteristics.

For the nuclear RNA enrichment analysis, our analytical approach relied on the classification of RNA molecules based on four parameters for each RNA molecule: tissue type (CTX, HP, TH, HY), cell type (including excitatory neurons, inhibitory neurons, glial cells, and their subtypes), gene type (among the thirty genes detected), and the RNA’s cellular location (either inside or outside the nucleus). RNA was divided into several categories across four dimensions, and the Ro/e for each subdivided category was obtained by dividing the observed count of RNA molecules by the expected count of RNA molecules calculated for each category.

To focus on the differences in RNA distribution across different tissues and cell types, we reduced the dimensionality for the gene type dimension by averaging Ro/e values. Due to the enrichment effect of marker genes in corresponding cell types, marker genes contribute significantly when averaging across the thirty genes (Supplementary Fig. 36b). We then reduced the dimensionality for the nuclear/extranuclear dimension by using the extranuclear/nuclear ratio, ultimately obtaining the average relative abundance of RNA from thirty genes inside and outside the nucleus across different tissue and cell types (Fig. 6g). Nuclear RNA enrichment analysis was validated using RNAscope, with *Plp1* selected as the representative gene (Supplementary Fig. 37b-d).

In addition to the nuclear RNA enrichment analysis, we observed that the distribution of some RNAs within the cell exhibited polarity. To reflect the overall polarity of RNA, we first calculated the ratio of the distance between each RNA’s centroid and the cell nucleus’s centroid to the average distance of each RNA to the cell nucleus’s centroid for each cell. This ratio was used to indicate the overall polarity of RNA, and after compiling statistics for all cells, we obtained a quasi-normal distribution with a mean of 0.35. We designated the top 20% of this distribution as high polarity, the bottom 20% as low polarity, and the middle portion as moderate polarity, thus setting the thresholds for polarity. Subsequently, to reflect the variability among different genes, we focused on genes with counts of more than 5 per cell for our analysis. We calculated their relative polarity using the aforementioned method and classified them into one of three categories (low, medium, high) based on the calculated thresholds.

Similar to the nuclear and extranuclear analysis, we classified each RNA molecule based on four parameters: tissue type (CTX, HP, TH, HY), cell type (excitatory neurons, inhibitory neurons, glial cells, and their subtypes), gene type (the thirty genes we detected), and the polarity of the gene within the cell (low, medium, high). RNA was categorized across four dimensions, and the Ro/e value for each category was obtained. Likewise, since our main interest was in the differences in RNA polarity distribution across different tissues and cell types, we reduced the dimensionality for the gene type dimension by averaging different gene types. Due to the enrichment effect of marker genes in corresponding cell types, resulting in higher Ro/e values, thus making marker genes contribute significantly when averaging across the thirty genes (Supplementary Fig. 36b). We further reduced the dimensionality for the polarity dimension by using the high-to-low polarity ratio, ultimately obtaining the average RNA polarity distribution of thirty genes across different tissue and cell types (Fig. 6f). It’s worth noting that since we assigned a high, medium, or low polarity to each RNA molecule, the higher the RNA counts of a particular gene in a cell, the greater its impact on the polarity calculation result. To validate subcellular RNA polarity, we used an in situ sequencing dataset from mouse brain sections^26^ that includes subcellular information. Due to different cell type annotations between datasets, analysis focused on shared marker genes with well-defined subcellular localization patterns. For each gene, polarity was quantified as the normalized distance between the transcript centroid and the nuclear centroid, scaled by the cell’s longest axis to control for size variation. RNA polarity measurements were consistent between datasets (spearman correlation r = 0.857, Supplementary Fig. 37a).

## Data availability

Single cell RNA-seq data were obtained from mousebrain.org, GEO accession number (GSE151530, GSE140228) and CNP0000650. Raw data of this study was deposited in zenodo including raw images (HCC: 10.5281/zenodo.12750711; MouseEmbryo: 10.5281/zenodo.12750725; MouseBrain: 10.5281/zenodo.12673246) and analysis related data (10.5281/zenodo.12755414)^68–71^. A website (http://www.spatialprism.org) is available to provide a clear understanding of PRISM’s capabilities.

## Code availability

Source code is provided in Github repository at: https://github.com/HuangLab-PKU/PRISM-Code^72^ for gene calling pipeline and https://github.com/HuangLab-PKU/PRISM-Analysis^73^ for post-gene-calling analysis for this manuscript.

