## Extended Data Figs 1-10 for "High-plex spatial RNA imaging in one round with conventional microscopes using color-intensity barcodes"

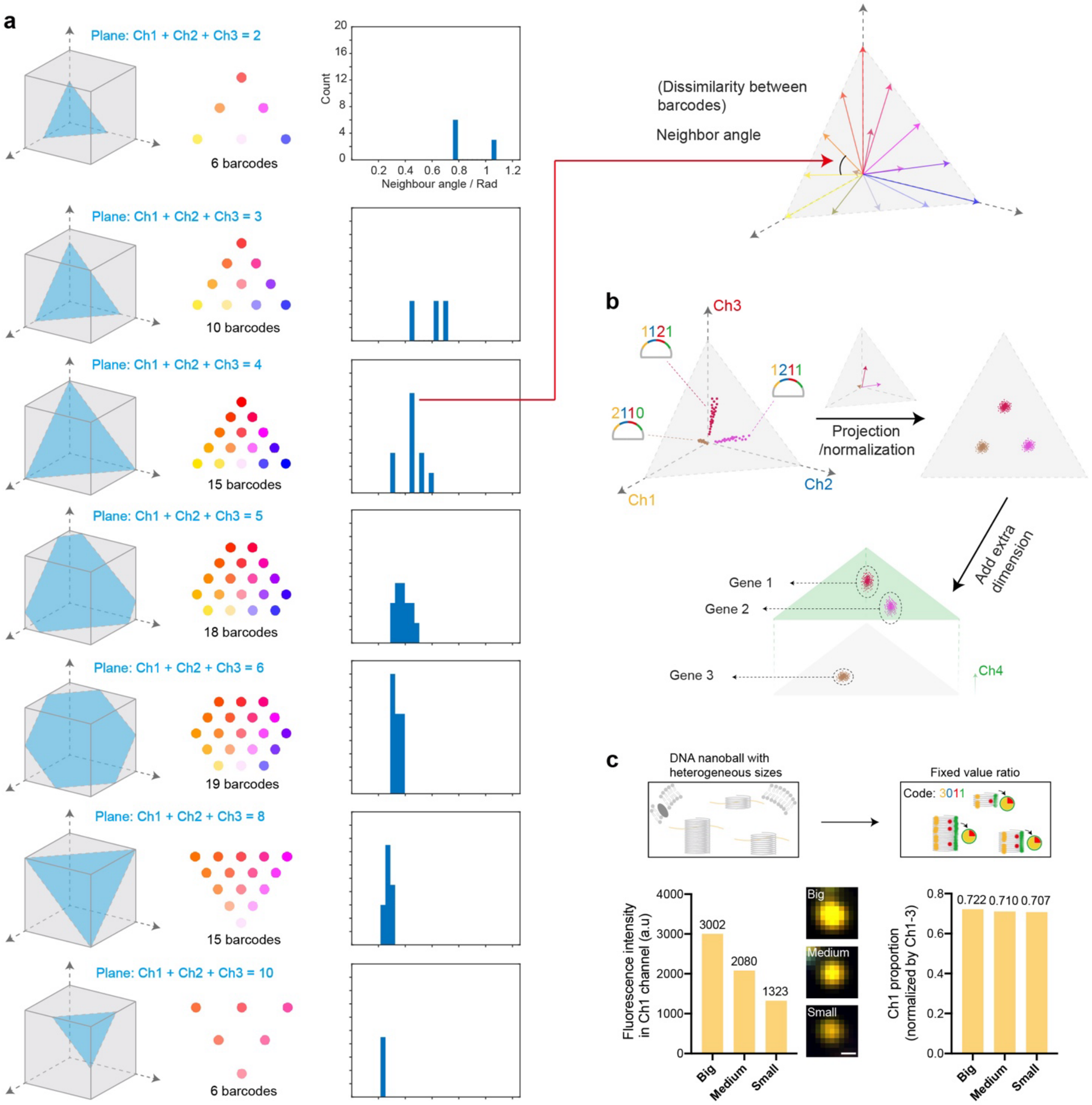


**Extended Data Fig. 1 | Radius vector filtering. a**, Selection of different intersecting planes in color space. Different planes correspond to different barcode vector sets with various numbers of vectors. A well-balanced barcode set should consider both the number of barcodes and the neighbor angle. The neighbor angle, which indicates the dissimilarity between the closest barcodes (vectors), must be large enough to ensure accurate decoding of the barcodes. **b**, Workflow of signal spot decoding. In color space (Ch1, Ch2, Ch3), all spots were L1-normalized to the sum value of (Ch1+Ch2+Ch3), making all spots sum into ‘1’. All spots were therefore projected onto a 2-D plane. Subsequently, the Ch4 value was added as the third axis, creating a new 3-D color space that visualizes all 30 clusters. **c**, PRISM barcode after radius vector filtering is robust to bias from heterogeneous amplification in RCA. Within the same barcode, absolute values of specific channel may vary between different rolling circle DNA nanoballs due to RCA heterogeneity, but the ratios of these values remain consistent. Scale bar: 500 nm.

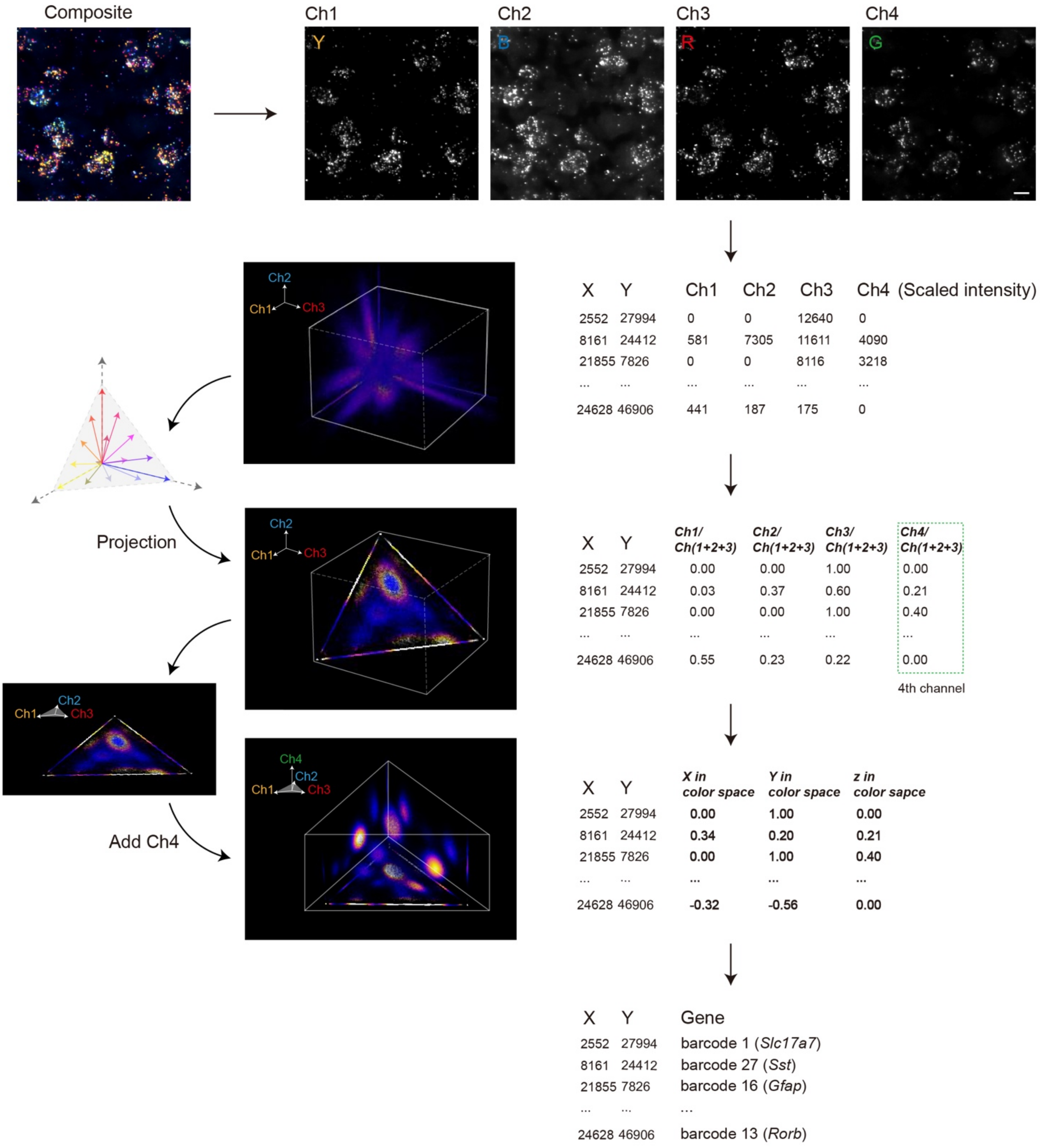


**Extended Data Fig. 2 | Creation of color space.** Raw intensity from four channels was extracted for each spot. After intra-spot L1-normalization by sum (Ch1+Ch2+Ch3), intensity information from four channels (Ch1’, Ch2’, Ch3’, Ch4’) was converted into three-dimensional coordinates in color space (X in color space: 2*Ch3’-1, Y in color space: Ch2’-Ch1’, Z in color space: Ch4’). Scale bar: 10 μm.

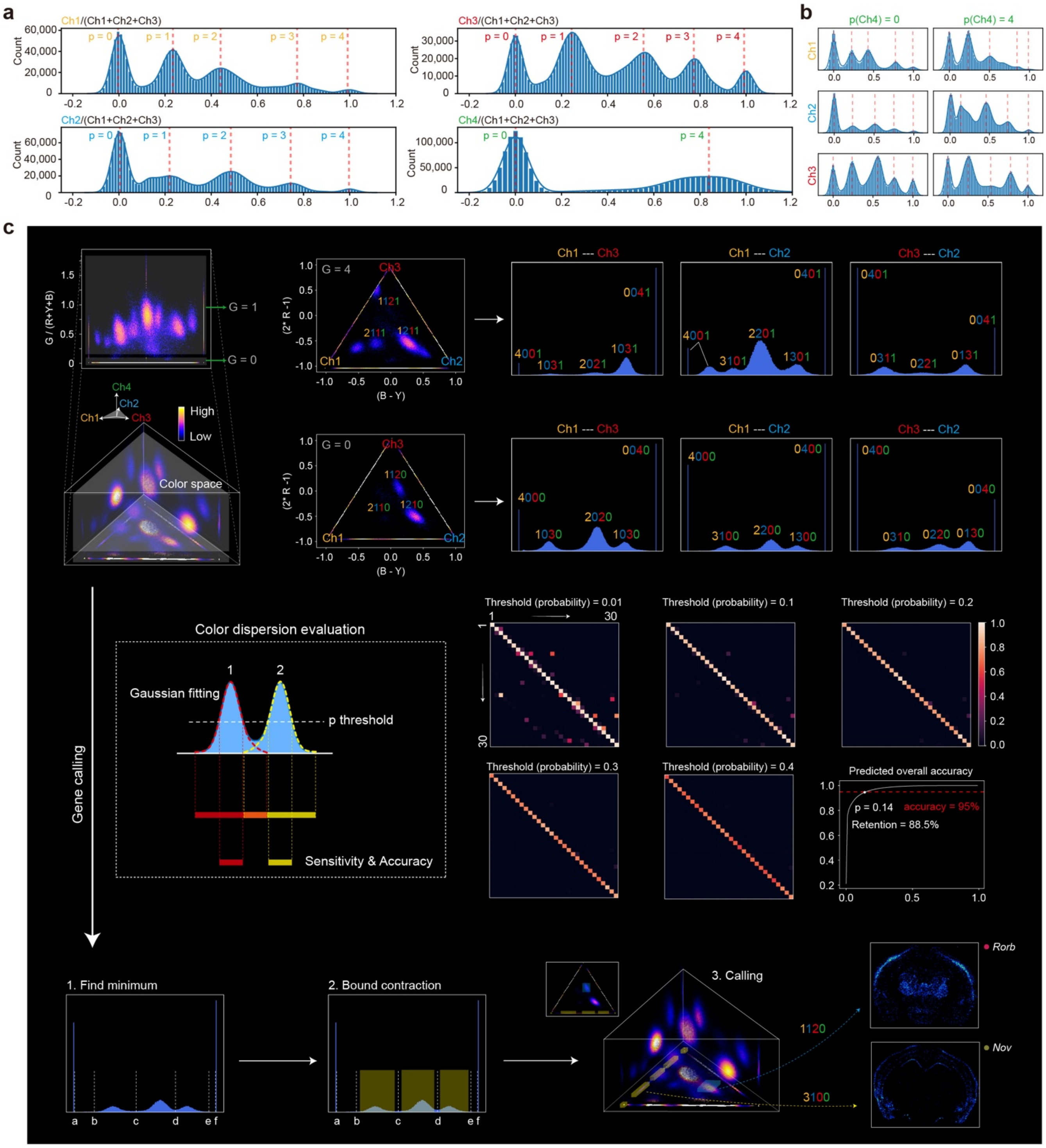


**Extended Data Fig. 3 | Gene calling workflow. a**, Intensity distribution of each channel after normalization. A small random number was introduced to coordinates from each spot to improve the assessment of cluster distribution. This modification results in the formation of a pseudo-Gaussian distribution at endpoint “0” and “1” (*e.g.* barcode 4000, 0040), allowing for an equal evaluation between these endpoint clusters and other clusters. **b**, Intensity distribution of Ch1, Ch2 and Ch3 under Ch4=0 and Ch4=1, respectively. **c**, Gene calling based on spots position in color space. The distribution of the 30 barcode clusters can be observed through different cross-sections and projections. Gaussian fitting was utilized to measure the separation between adjacent clusters. Manual delineation of three-dimensional boundaries for each cluster in the color space was performed for gene calling.

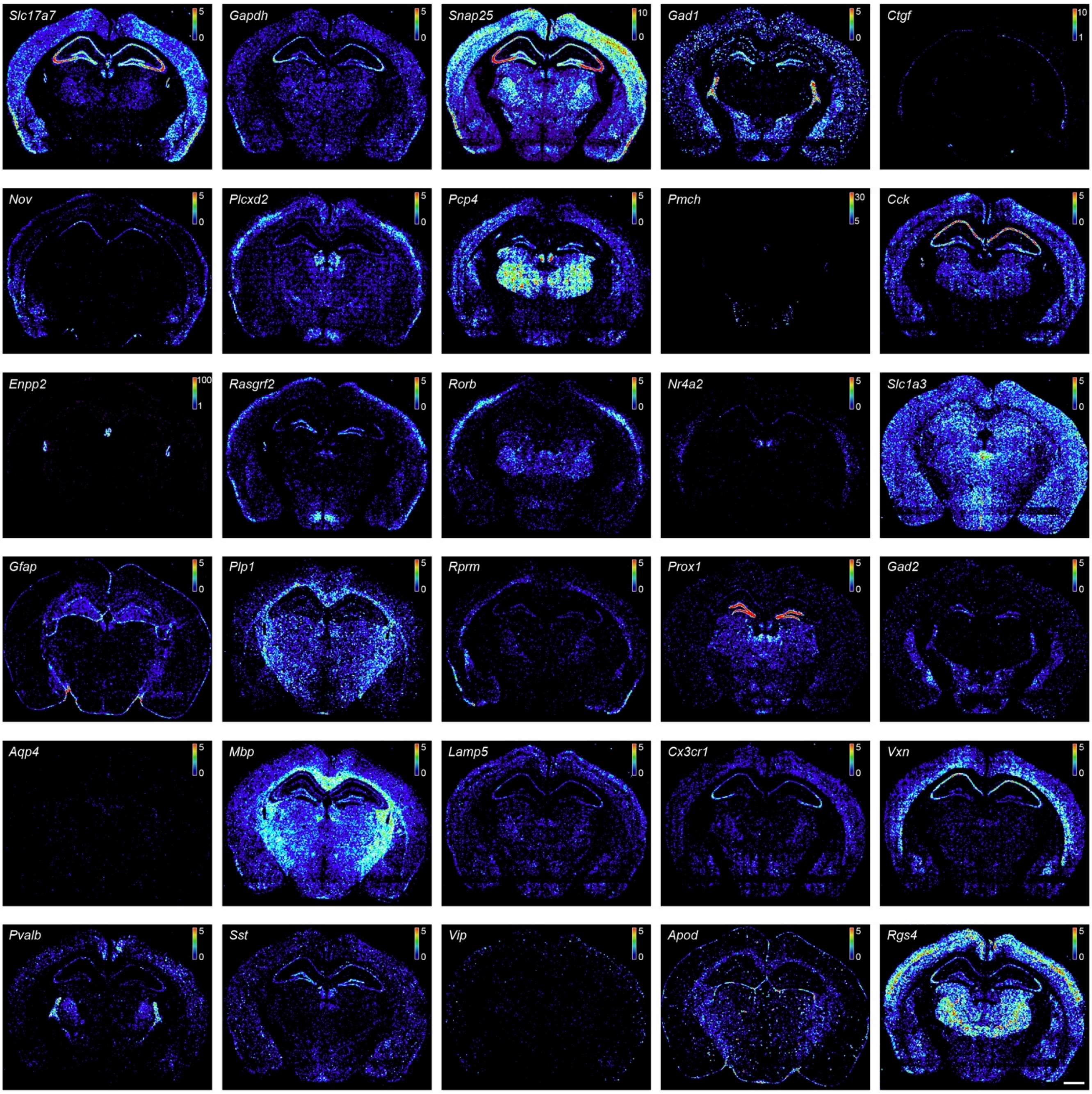


**Extended Data Fig. 4 | Spatial expression patterns of 30 called genes for mouse brain coronal section.** For better visualization, the transcripts are down-sampled (coarse-grained). The dynamic range of transcripts density (counts per 100 x 100 pixel$^2$, 1 pixel = 0.1625 μm) is represented as shown in the colormap in each image. Scale bar: 1 mm.

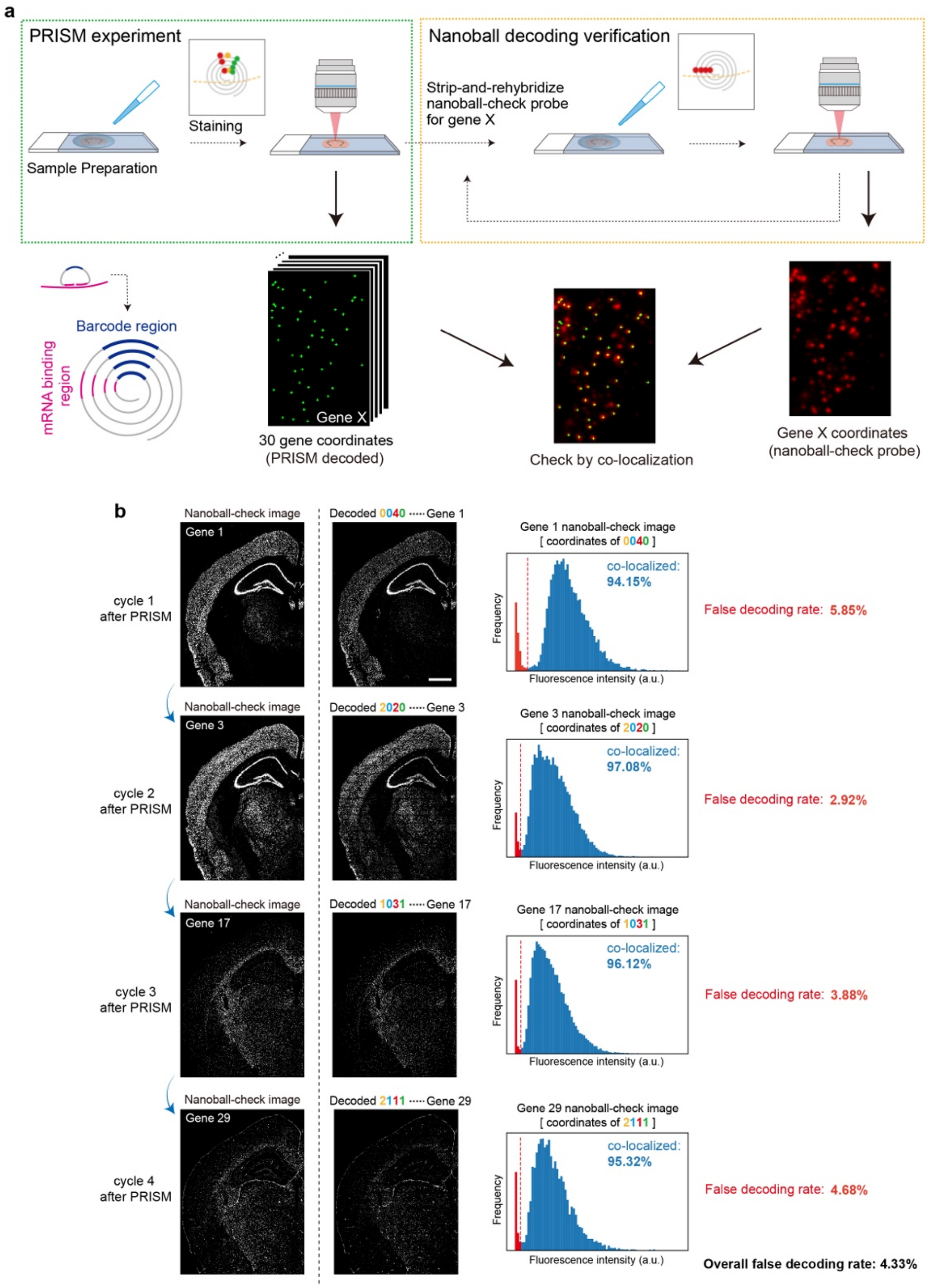


**Extended Data Fig. 5 | PRISM decoding accuracy characterization. a**, Experimental workflow of PRISM decoding accuracy check. After PRISM fluorescent staining and gene identification, the imaging probes were stripped away. Subsequently, fluorescently labeled nanoball-check probe that specifically targeted at rolling circle nanoballs (targeting mRNA-binding region, shown as magenta) of individual gene was applied to reveal the ground truth position of nanoball. This process was performed iteratively within a flow cell for checking different genes. We then overlapped the decoded specific gene coordinate (from PRISM experiment) with corresponding gene's nanoball-check image one-by-one to examine decoding accuracy. **b**, Decoding accuracy calculation. The decoding accuracy for each gene was calculated as the ratio of PRISM-decoded spots overlapping with nanoball-check-probe stained spots to the total PRISM-decoded spots. Fluorescent intensity value was obtained by reading the nanoball-check image using coordinates decoded by PRISM. The intensity frequency distribution was plotted to characterize the decoding accuracy. Correctly decoded spots (overlapping with nanoball-check-probe stained spots) formed a near-Gaussian distribution (right peak), while false-decoded spots ("non-overlapping") appeared as a sharp left peak . The proportion of non-overlapping coordinates defined the false decoding rate. Four different barcodes/genes were analyzed, yielding an average false decoding rate of 4.33%. Scale bar: 1 mm.

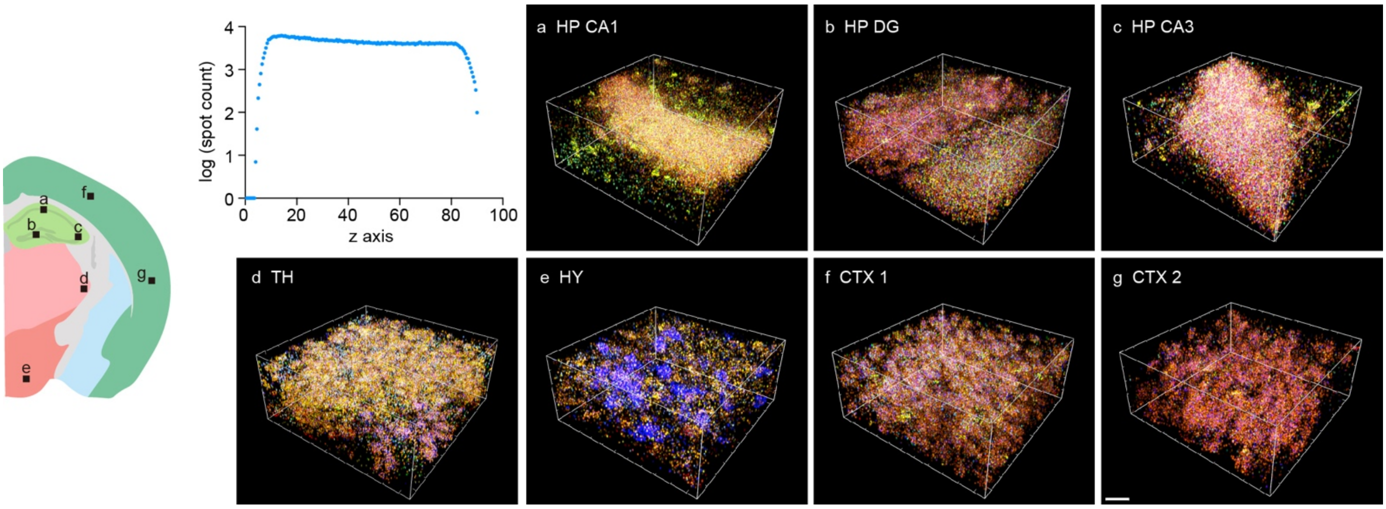


**Extended Data Fig. 6 | Reagent penetration in thick tissue.** Seven distinct regions (200 µm x 200 µm x 100 µm) within the mouse brain were selected for conducting a reagent penetration test. The penetration performance in thick tissue was reflected by overall extracted signal distribution along z-axis. Thickness: 100 µm, Scale bar: 30 µm.

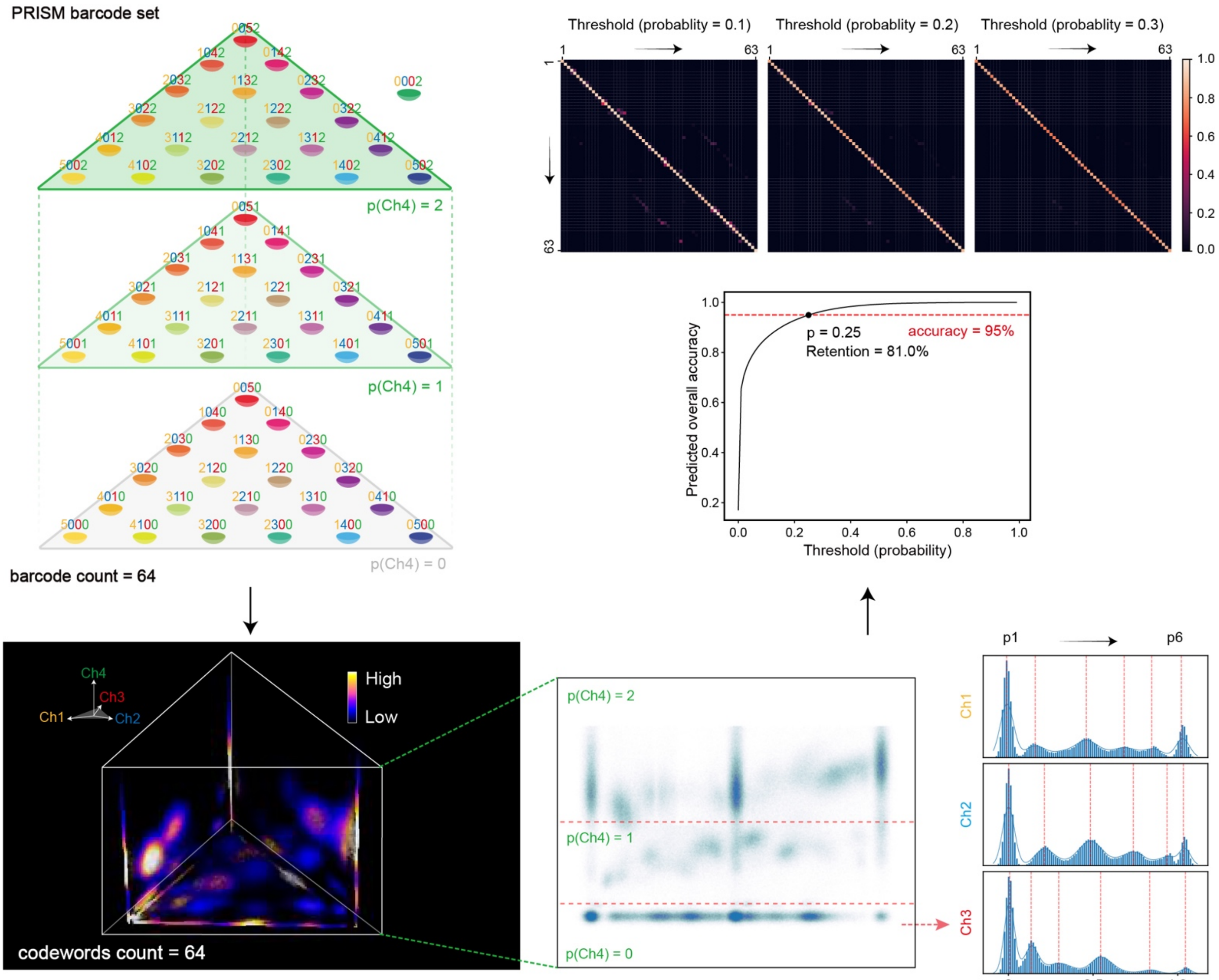


p(Ch1/2/3) = k, p(Ch4) = j:

$$\text{codewords count} = j \times \sum_{a=0}^{k} a + 1$$

**Extended Data Fig. 7 | PRISM Barcode expansion to 64 barcodes by increasing intensity levels.** Ch1, Ch2 and Ch3 were quantized into fifths (0, 1/5, 2/5, 3/5, 4/5 and 5/5) and Ch4 was divided into high (2), low (1) and zero (0). This expansion resulted in an increased coding capacity of 64 (21*3+1).

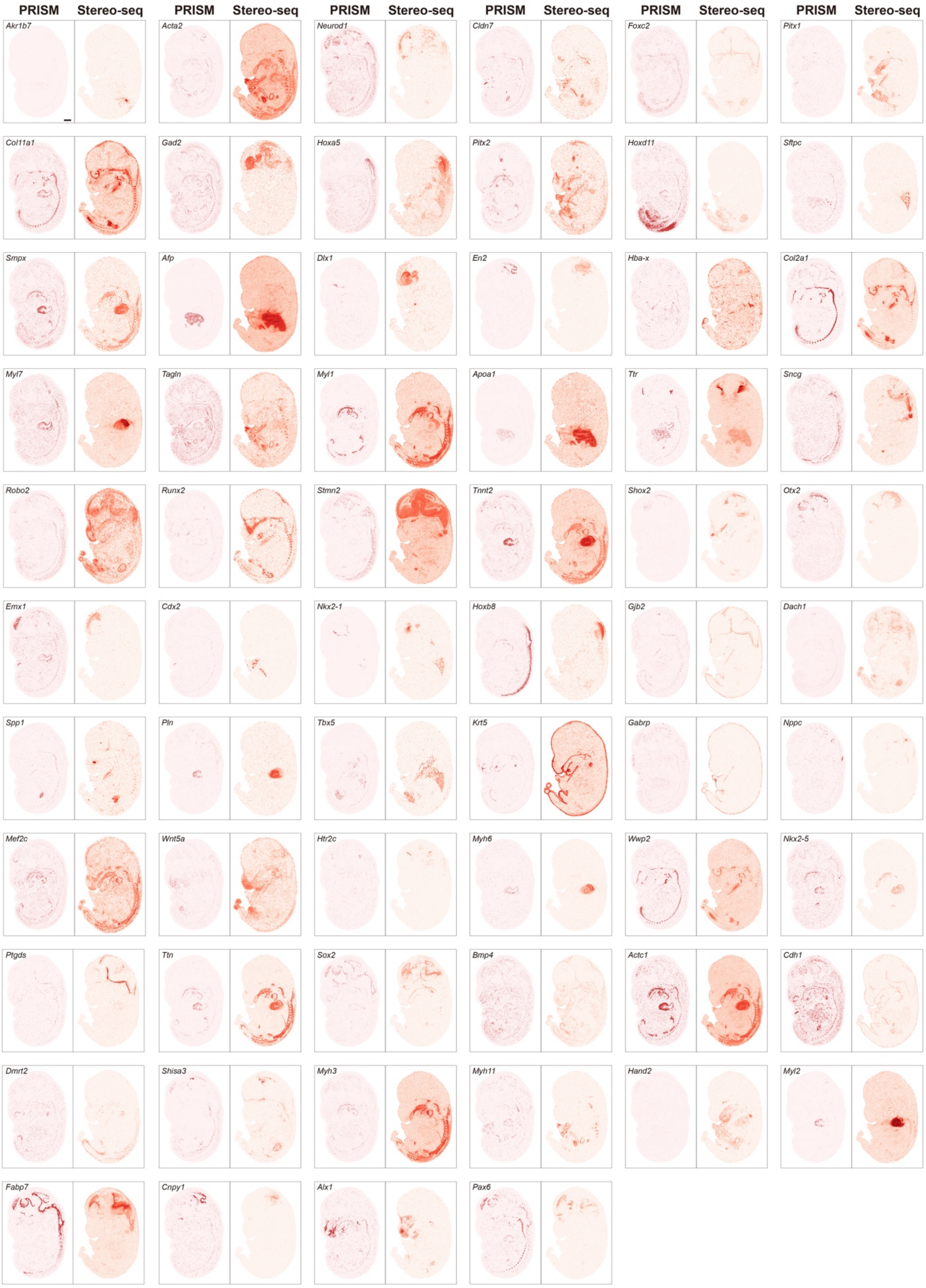


**Extended Data Fig. 8 | Spatial expression pattern comparison between 64-plex PRISM and MOSTA database.** This comparison was performed using sagittal sections of mouse embryos at E14.5 stage. Selected data has a similar sectioning position (matched anatomical regions) as our data. Scale bar: 1 mm.

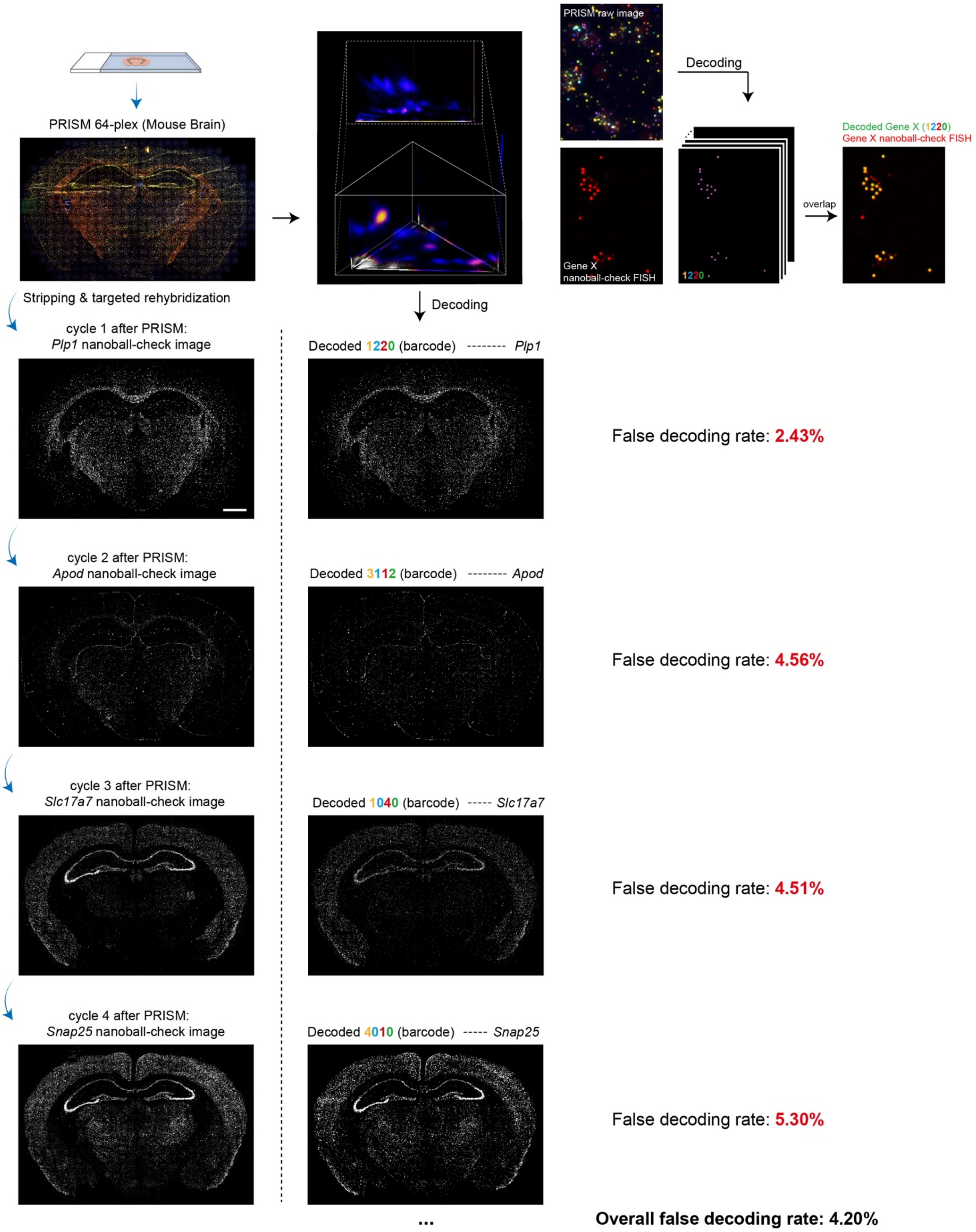


**Extended Data Fig. 9 | PRISM decoding accuracy characterization for 64-plex assay on mouse brain tissue.** Experiment and analysis were performed the same with Extended Data Fig. 5. Briefly, after PRISM fluorescent staining and gene identification, the imaging probes were stripped away. Subsequently, fluorescently-labeled nanoball-check probe that specifically targeted sat rolling circle nanoballs (targeting mRNA-binding region) of individual gene was applied to reveal the ground truth position of nanoball. The decoding accuracy for each gene was calculated as the ratio of PRISM-decoded spots overlapping with nanoball-check-probe stained spots to the total PRISM-decoded spots. The proportion of non-overlapping coordinates defined the false decoding rate. The average false decoding rate is calculated to be 4.20%. Scale bar: 1 mm.

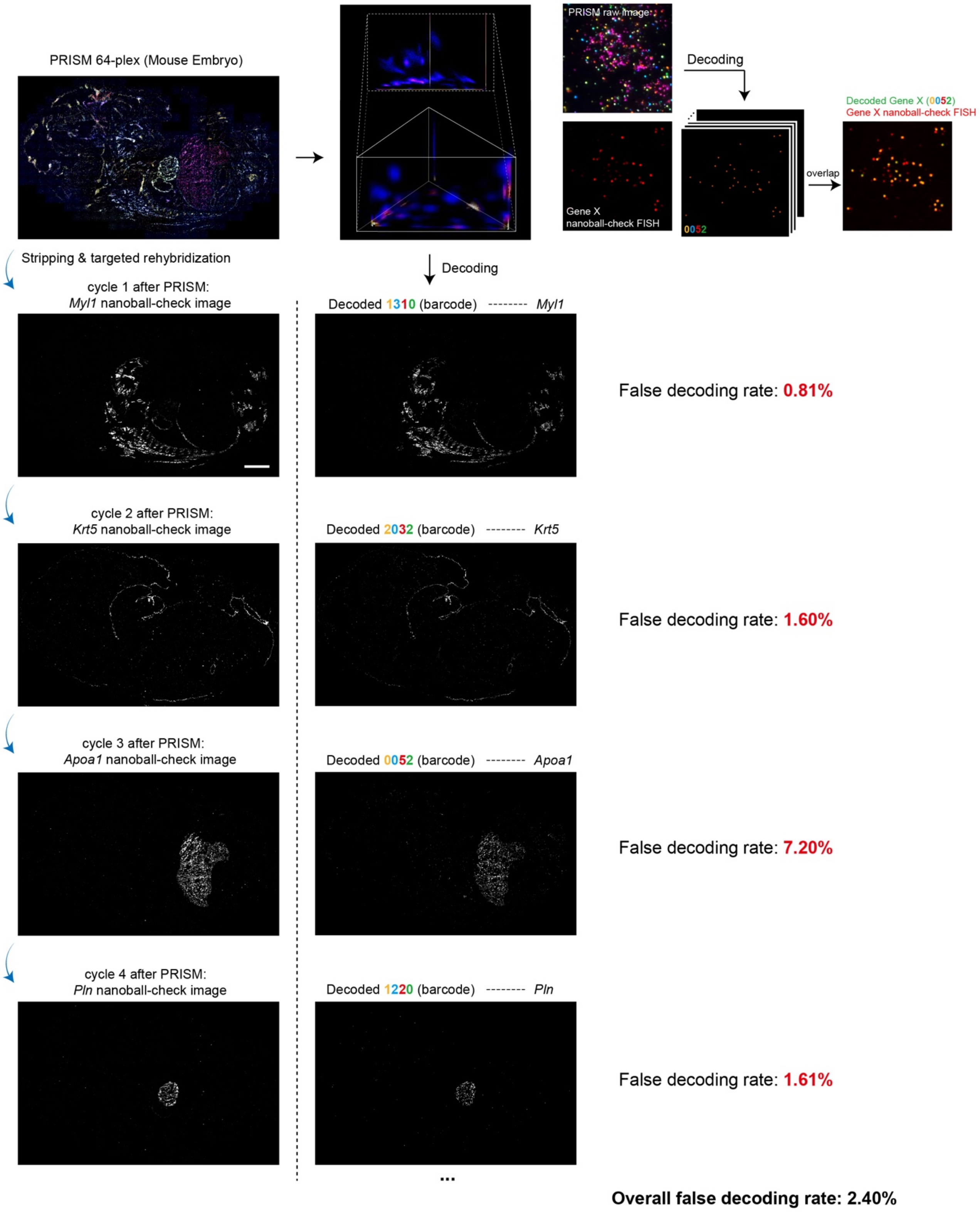


**Extended Data Fig. 10 | PRISM decoding accuracy characterization for 64-plex assay on mouse embryo tissue.** Experiment and analysis were performed the same with Extended Data Fig. 5. Briefly, after PRISM fluorescent staining and gene identification, the imaging probes were stripped away. Subsequently, fluorescently-labeled nanoball-check probe that specifically targeted at rolling circle nanoballs (targeting mRNA-binding region) of individual gene was applied to reveal the ground truth position of nanoball. The decoding accuracy for each gene was calculated as the ratio of PRISM-decoded spots overlapping with nanoball-check-probe stained spots to the total PRISM-decoded spots. The proportion of non-overlapping coordinates defined the false decoding rate. The average false decoding rate is calculated to be 2.40%. Scale bar: 1 mm.
