## Supplementary Figs 1-41 for "High-plex spatial RNA imaging in one round with conventional microscopes using color-intensity barcodes"

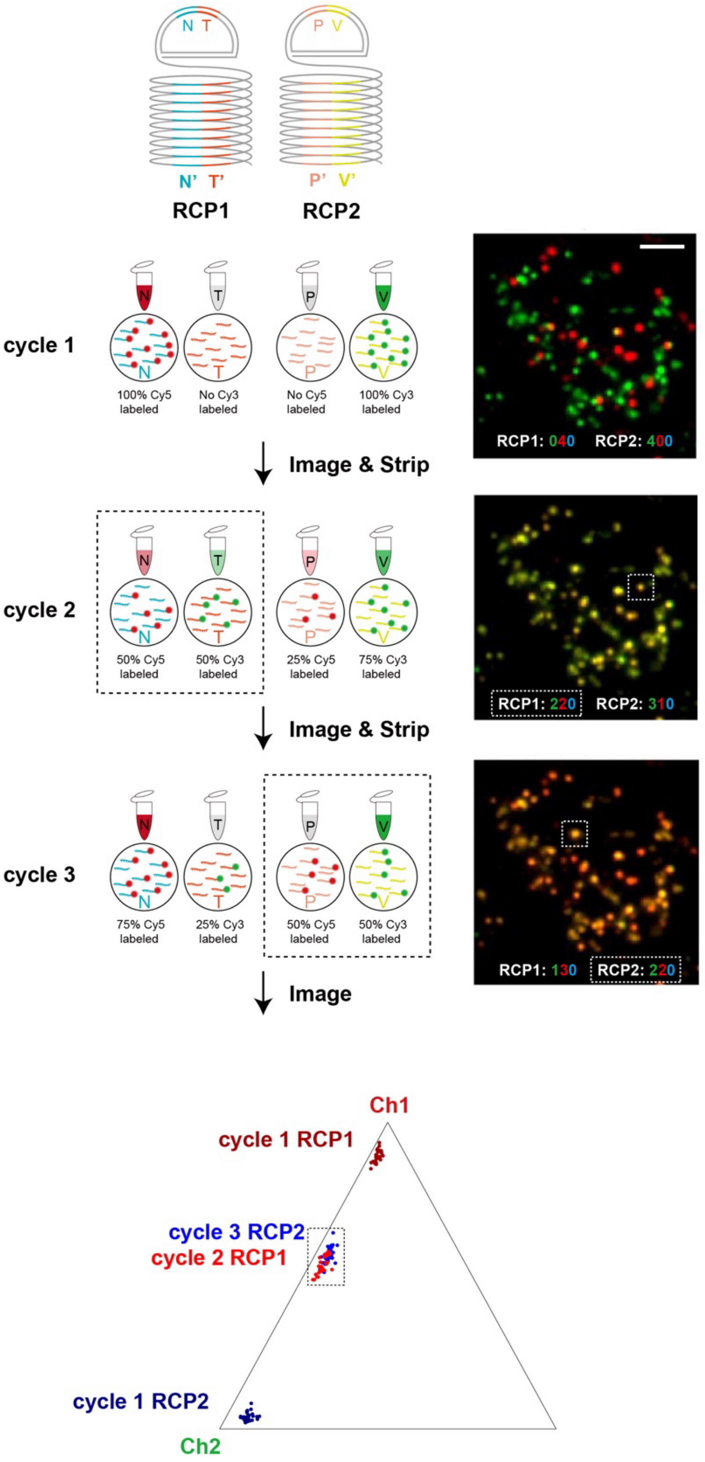


**Supplementary Fig. 1 | Robustness of barcode identification with different imaging probe sequences.** To evaluate whether differences in imaging probe sequences could lead to variations in hybridization efficiency and thus affect barcode identification, two amplification products (RCPs) with different barcode sequences were generated on the same sample. These RCPs underwent multiple imaging cycles within a flow cell (imaging probe hybridization, imaging, stripping, imaging probe hybridization,…), with adjusted ratios of the corresponding imaging probes in each cycle to ensure that both RCPs presented intended barcode signal. In cycle 1, RCP1 and RCP2 were labeled with 100% Cy5 and 100% Cy3, respectively, for spatial localization. In cycles 2 and 3, RCP1 and RCP2 were configured to display the '022' barcode, and their corresponding positions in color space were examined for consistency. The resulting '022' signal from two RCPs occupied the same positions in color space, demonstrating that differences in imaging probe sequences do not impact barcode identification. Scale bar: 5 μm.

**Pseudocode part 1**

```
// Generate all 3-Chn barcodes (Level 0-4)
all_barcodes = []

for i from 0 to 4:
    for j from 0 to 4:
        for k from 0 to 4:
            all_barcodes.append(str(i) + str(j) + str(k))

return all_barcodes
```

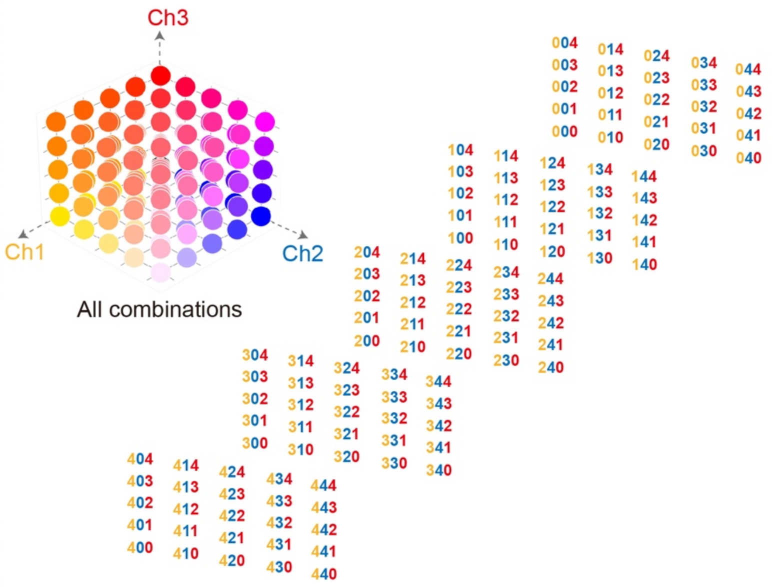


**Pseudocode part 2**

```
// Filter barcodes where i + j + k = 4
filtered_barcodes = []

for barcode in all_barcodes:
    i = int(barcode[0])
    j = int(barcode[1])
    k = int(barcode[2])
    if i + j + k == 4:
        filtered_barcodes.append(barcode)

// filtered_barcodes now contains
// 15 matching combinations
```

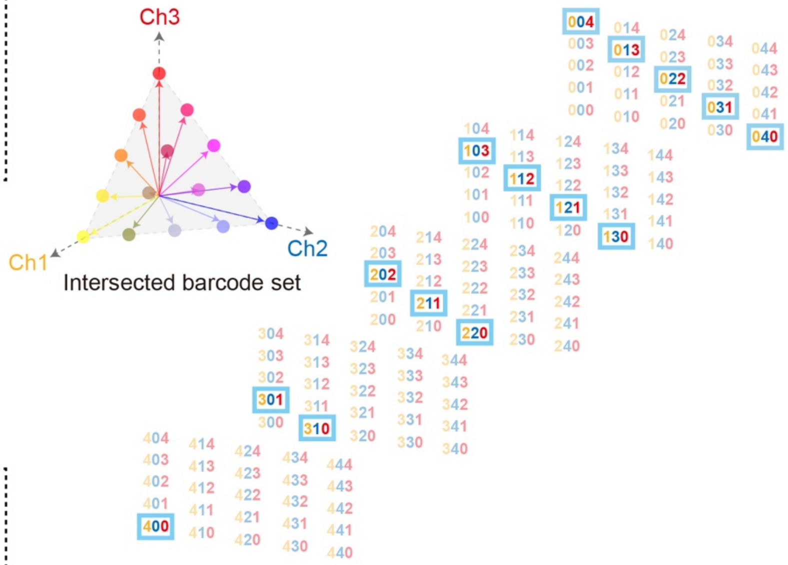


**Pseudocode part 3**

```
// Expand barcode with 4th Chn (0 or 1)
expanded_barcodes = []

for barcode in  filtered_barcodes:
    // Append 0
    expanded_barcodes.append(barcode + "0")
    // Append 1
    expanded_barcodes.append(barcode + "1")

// expanded_barcodes now contains
//30 final barcodes (15×2)
```

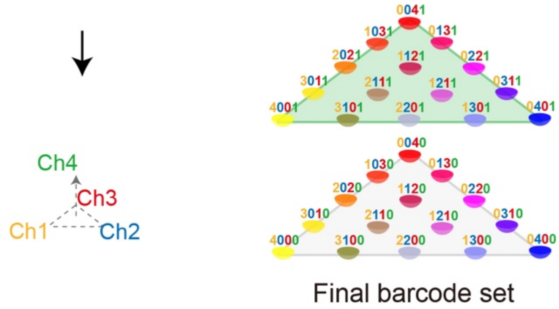


**Supplementary Fig. 2 | Pseudocode diagram for 30-barcode generation using 4-channel PRISM color-coding.** The process consists of three steps: (1) three spectral channels with each quantized into quarters (0, 1/4, 2/4, 3/4, 4/4), generating $5^3 - 1 = 124$ possible barcodes through combination; (2) 'radius vector filtering' to select 15 highly distinguishable barcodes; (3) barcode expansion by adding a fourth channel with 2 intensity levels, resulting in 30 final barcodes.

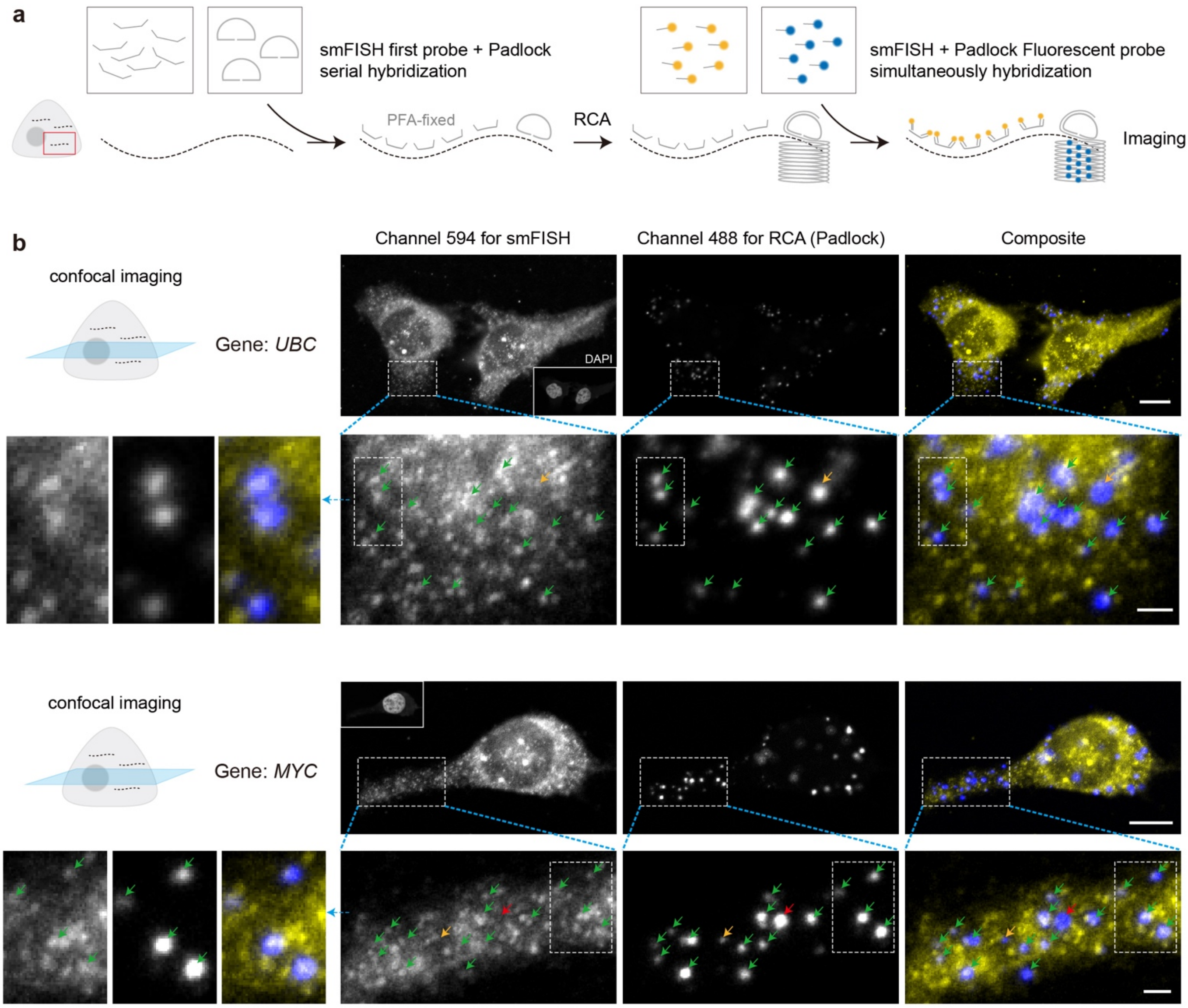


**Supplementary Fig. 3 | Stellaris smFISH and RCA simultaneous staining for nanoball accuracy evaluation. a,** Experimental workflow. Since smFISH signal is susceptible to RNA degradation, smFISH probe hybridization step was prioritized before RCA. In order to better preserve smFISH signal during RCA, we adopted a two-stage smFISH probe strategy, where primary smFISH probe has a single strand end and is less vulnerable to Phi29 polymerase's 3'-5' exonuclease activity. Following RCA, simultaneous staining was performed using both secondary smFISH probes and RCP-targeted imaging probes, followed by confocal microscopy imaging. **b,** Co-localization result. TxRed (594 nm) and AF488 (488 nm) were used to label smFISH signal and Padlock (RCP), respectively. In zoom-in regions, green arrows indicate co-localized signals, yellow arrows indicate signals with a certain degree of deviation (uncertain whether smFISH and RCA target the same mRNA), and red arrows mark non-overlapping signals. This co-localization assay was performed for two genes: *UBC* and *MYC*. The simultaneous co-staining assessment was repeated three times with similar results. Scale bar: 10 μm, overview; 2 μm, below (zoom-in).

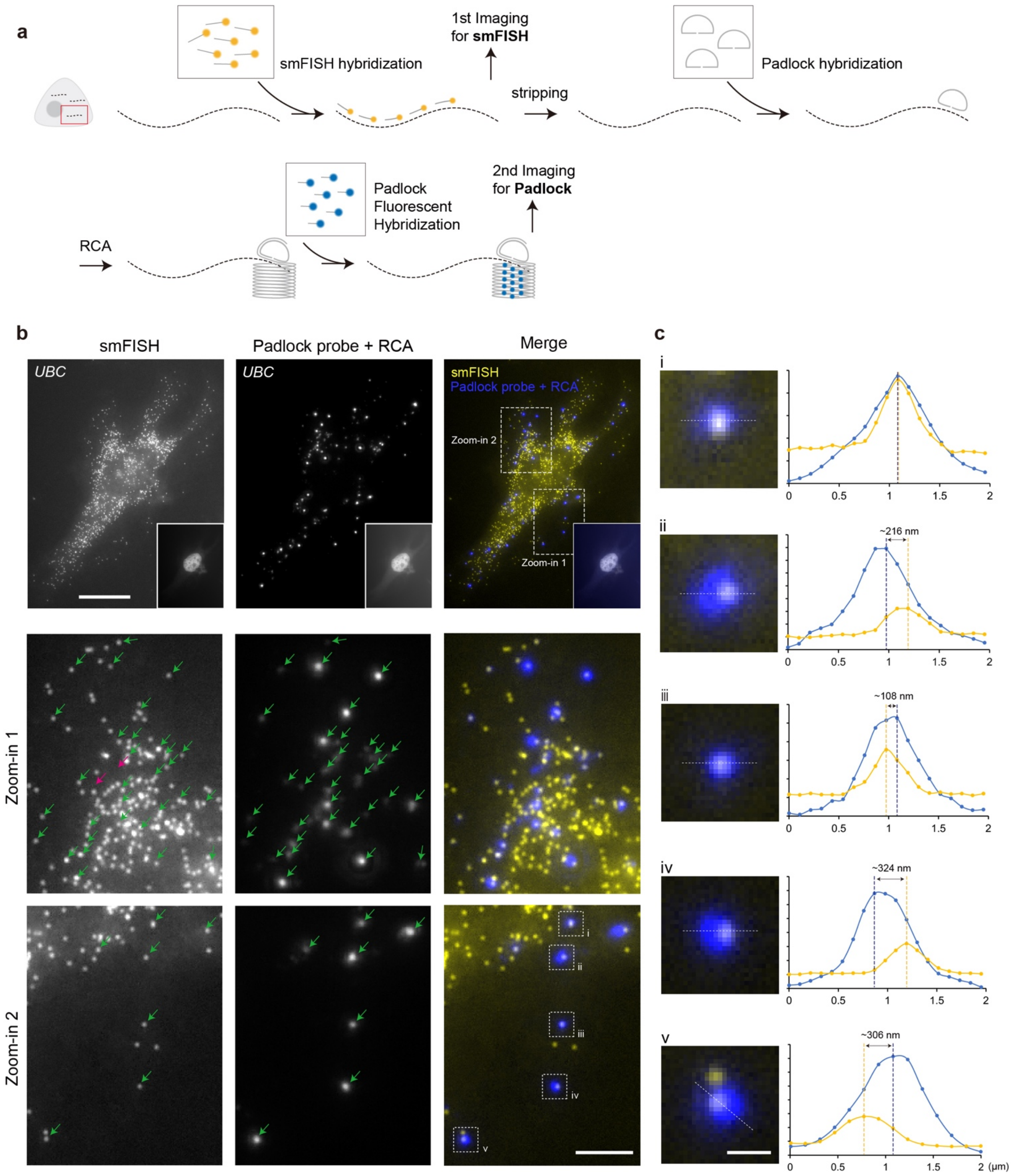


**Supplementary Fig. 4 | Serial detection using Stellaris smFISH and RCA for nanoball accuracy evaluation. a,** Experimental workflow. The smFISH experiment was also prioritized before RCA since it is susceptible to RNA degradation. In this experiment, fluorophore-labeled smFISH probe was used to directly hybridize sample and performed imaging, after which the smFISH probe was stripped and 'Padlock & RCA' (PRISM amplification) was performed, followed by imaging probes staining and widefield imaging. Fields of view were kept the same in two imaging rounds and images were registered. **b**, Co-localization result. Green arrows indicate co-localized signals and red arrows mark non-overlapping signals. The serial detection assessment was repeated three times with similar results. Scale bar: 20 μm, overview; 5 μm, below (zoom-in). **c**, The panel on the right shows the fluorescence intensity profile along the dashed line. Yellow represents the smFISH signal, while blue represents the RCA signal. Scale bar: 1 μm.

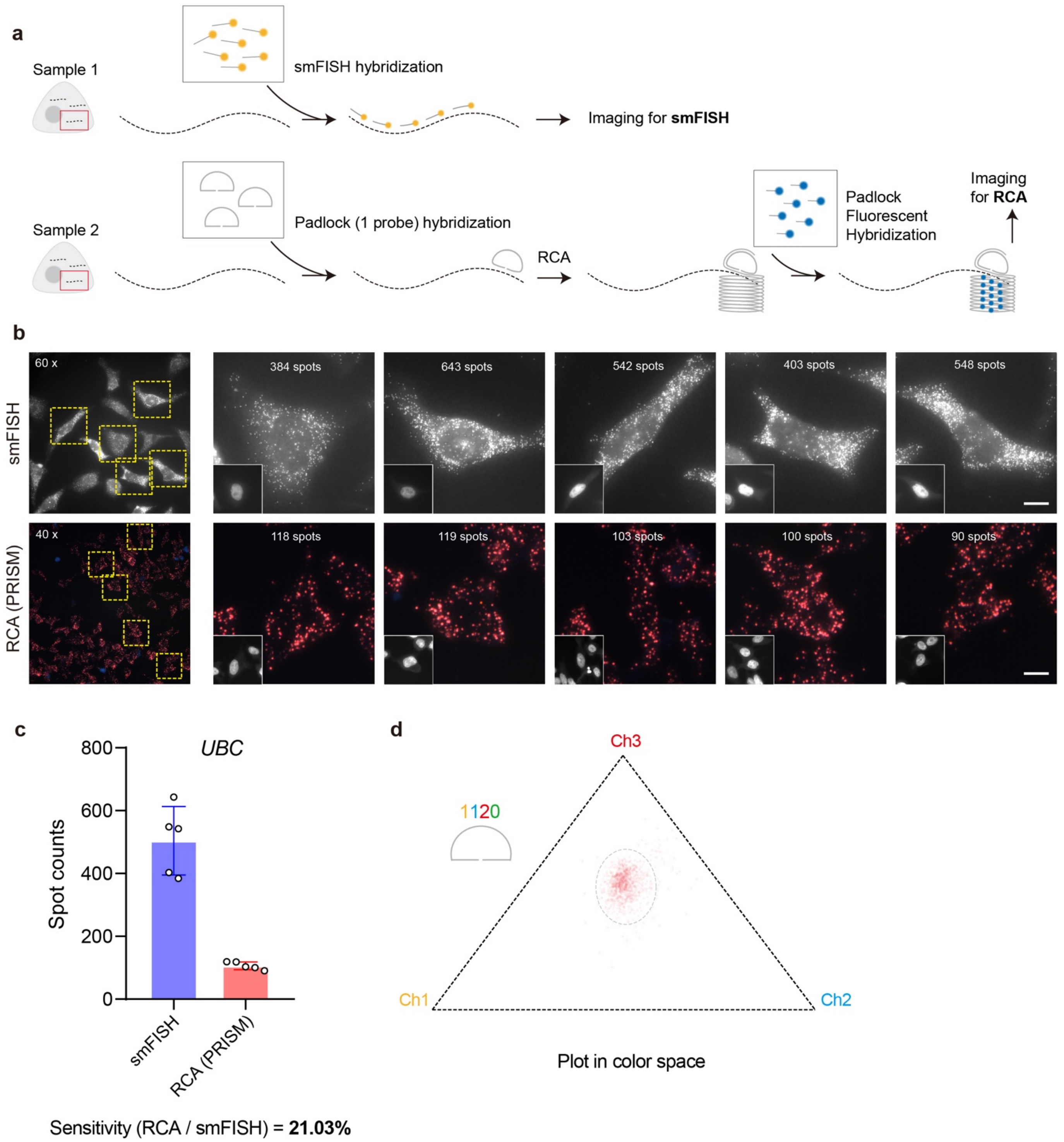


**Supplementary Fig. 5 | Stellaris smFISH and PRISM on parallel samples for RCA sensitivity evaluation. a**, Experimental workflow. smFISH and PRISM (one padlock probe) experiment were performed on different samples and mRNA counts were directly compared. The cell culture condition and pre-treatment procedures were kept consistent in two groups. Barcode for PRISM was set to be [1120]. **b**, Sensitivity result of smFISH and PRISM (RCA) for same gene. The sensitivity assessment was repeated three times with similar results. Scale bar: 10 μm. **c**, Sensitivity comparison between two methods. Bar plots show mean count ± s.d. (n = 5 randomly selected FoV for each group, shown in b). **d**, PRISM's signal plotting in color space.

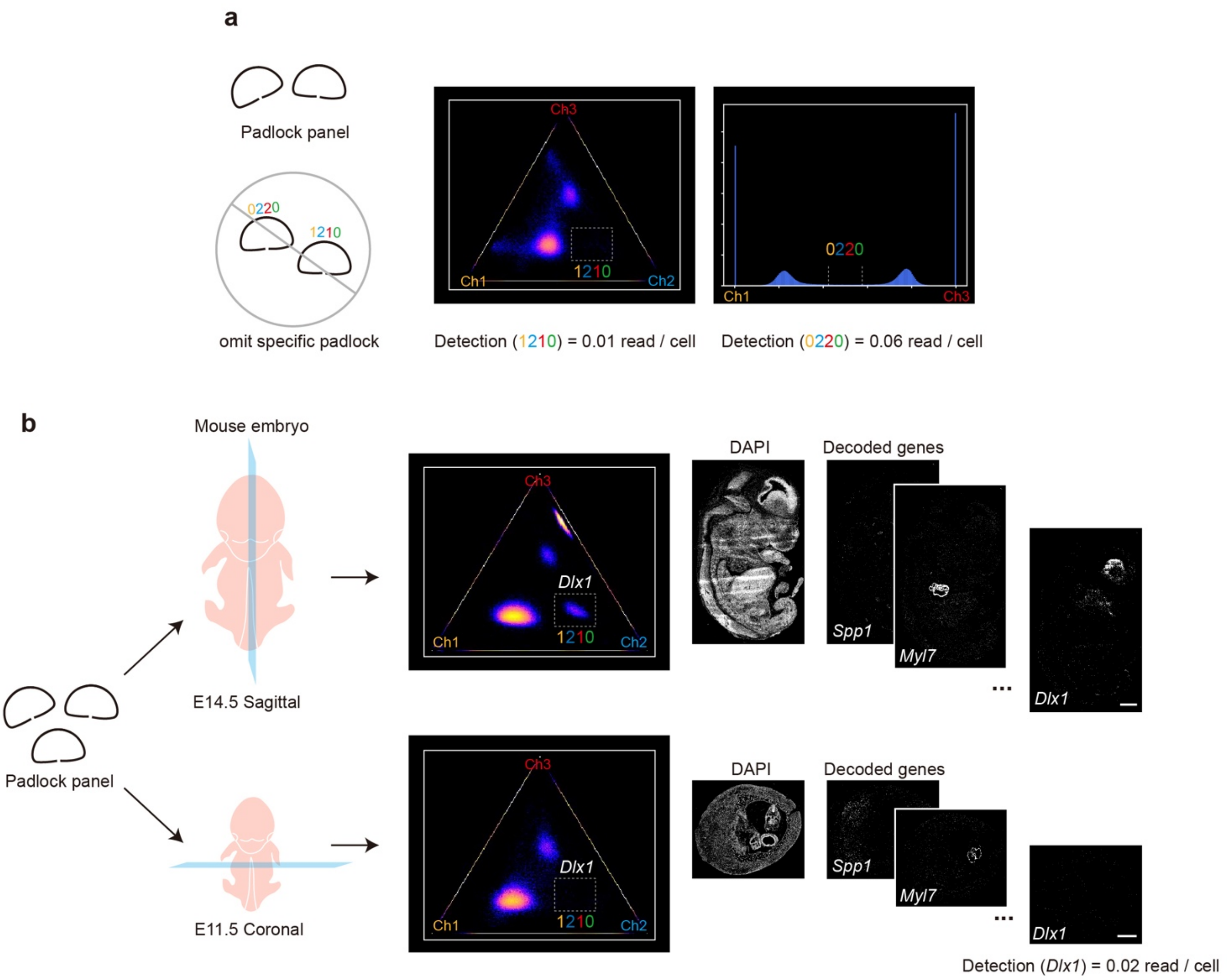


**Supplementary Fig. 6 | Negative controls for padlock targeting specificity and PRISM decoding. a,** False-positive control with omitted padlock probes. When padlock probes for barcodes [0220] and [1210] were excluded from the panel. Corresponding background detection rates (which comes from false detection) were 0.01 and 0.06 reads/cell, respectively. **b**, False-positive control experiment by targeting non-existing mRNA. *Dlx1* is mainly expressed in embryo brain, as shown in above part. In order to test false positive levels with no targets, the same gene panel was used to detect a coronal-section of E11.5 embryo, where *Dlx1* should not have signal. The detection rates of this gene showed 0.02 reads/cell. Scale bar: 1 mm.

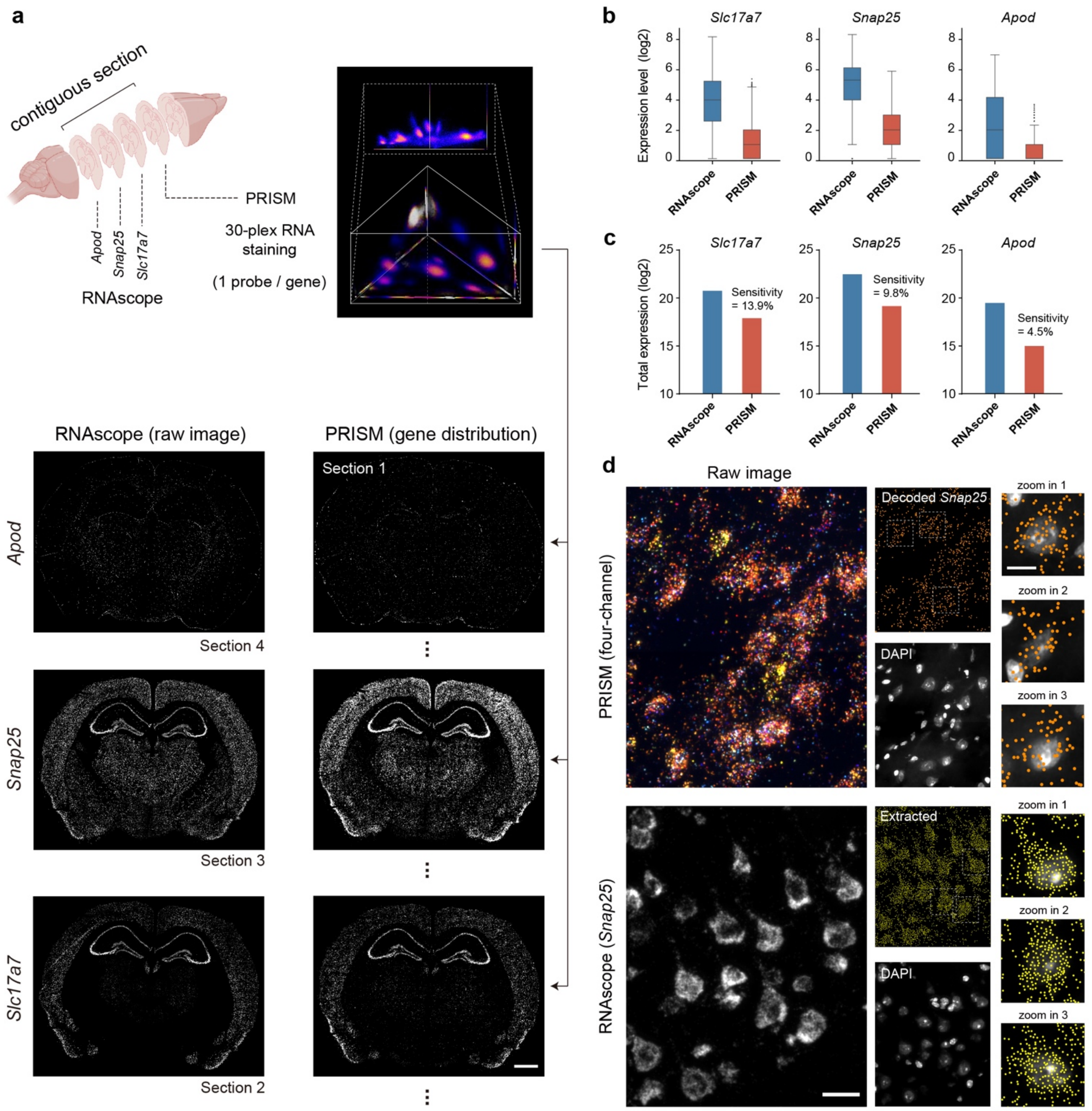


**Supplementary Fig. 7 | Benchmark with RNAscope using adjacent sections. a,** To systematically evaluate the accuracy (spatial expression pattern) and overall sensitivity (transcript counts) of PRISM experiment on tissue, we benchmarked a 30-plex PRISM (one padlock probe per gene) result with RNAscope. One brain section was used to perform PRISM and its adjacent sections were used to perform RNAscope for three different genes (*Apod*, *Snap25* and *Slc17a7*). Scale bar: 1 mm. **b,** Boxplots of three genes in cells with non-zero expression (log2 transformed). Boxplots display median (central line) and 25th-75th percentiles (box boundaries; whiskers at 1.5 IQR). **c,** Total detected RNA counts in brain section (log2 transformed). Sensitivity was calculated as the ratio of PRISM-decoded RNA counts to RNAscope-detected spots for each gene. **d,** Representative spots extraction of PRISM and RNAscope. The RNAscope assessment was repeated three times with similar results. Scale bar: 20 μm, overview (left); 10 μm, zoom-in image (right).

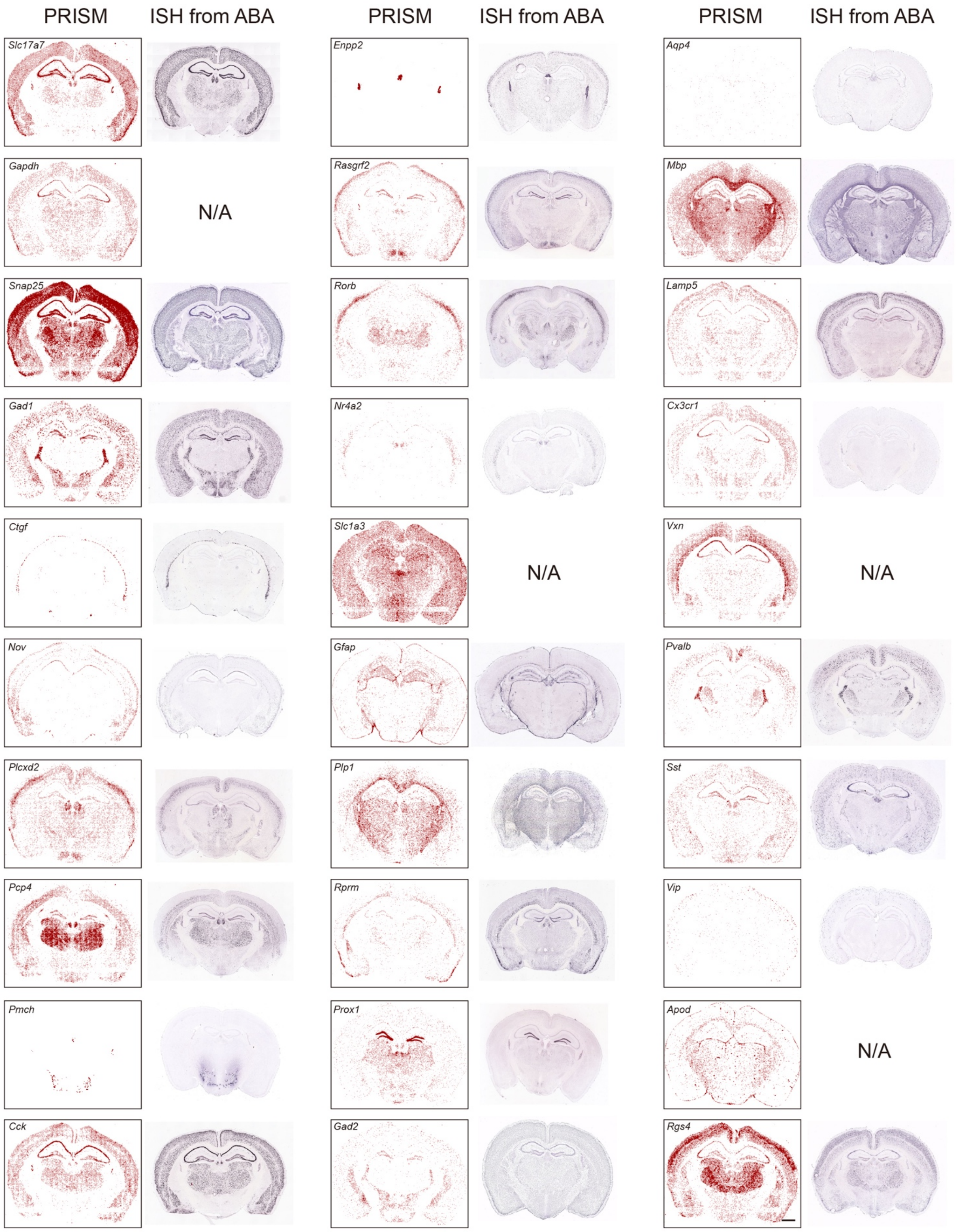


**Supplementary Fig. 8 | Spatial expression pattern comparison of 30 PRISM-called genes with Allen Brain Atlas.** Scale bar: 1 mm.

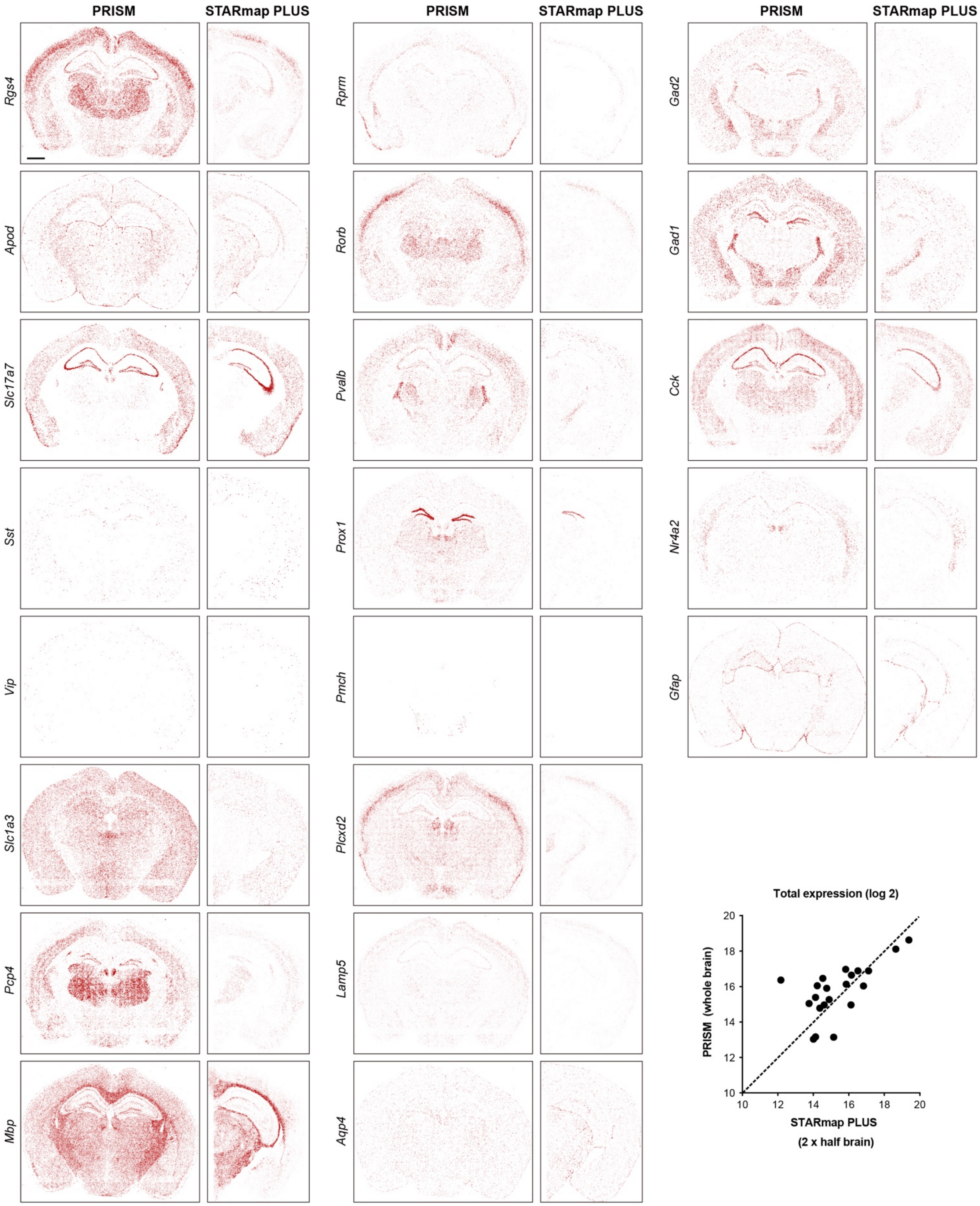


**Supplementary Fig. 9 | Spatial expression pattern comparison between PRISM and STARmap PLUS using shared genes.** The dynamic range of transcripts density was consistent between two methods. The STARmap PLUS dataset was from [Shi et al., Nature, 2023] well 06, which sample has a similar sectioning position (matched anatomical regions) as our data. For the comparative analysis (shown on the right below), STARmap PLUS total expression counts were doubled to account half-brain sampling. Scale bar: 1 mm.

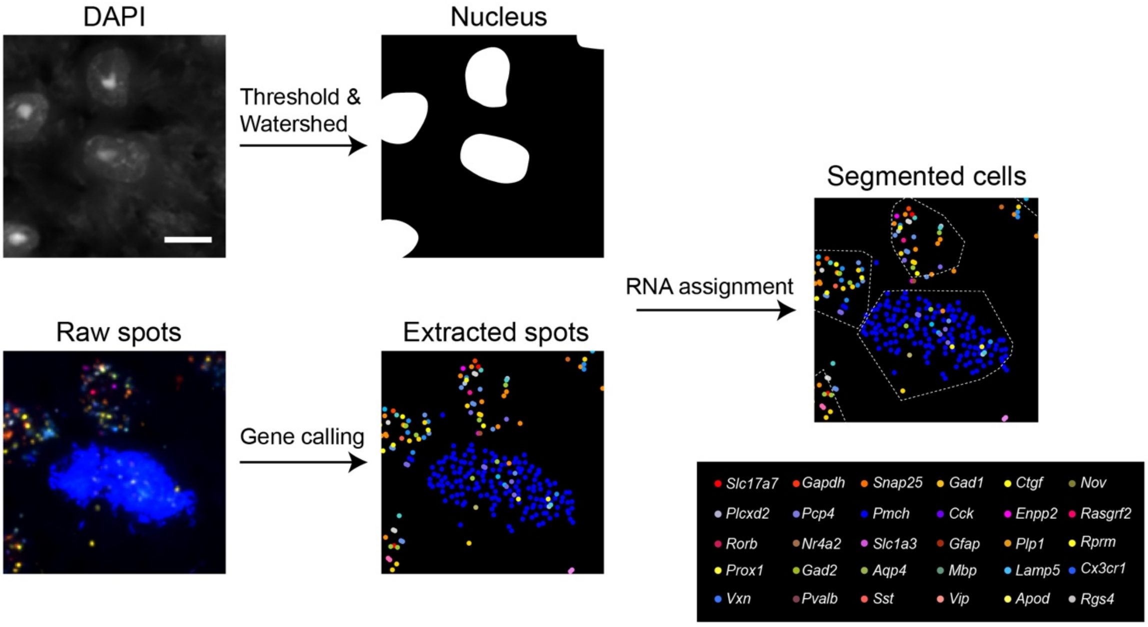


**Supplementary Fig. 10 | Cell segmentation and RNA assignment.** Cell nucleus was segmented through thresholding and watershed algorithm based on DAPI channel. The extracted RNA spot was assigned to the centroid of the nearest nucleus based on Euclidean distance within a designated range. Scale bar: 10 μm.

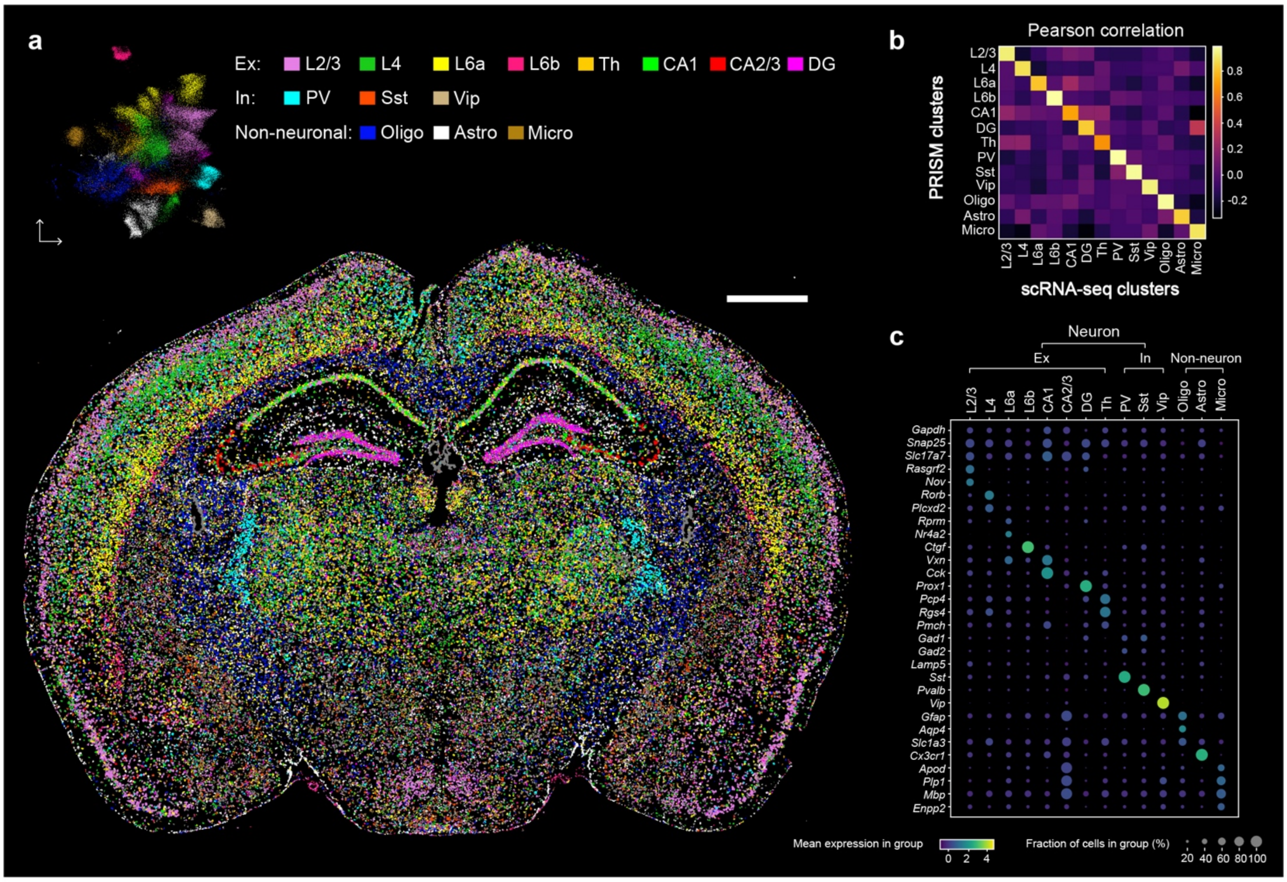


**Supplementary Fig. 11 | Cell classification in mouse brain based on PRISM staining. a**, Classified cells projection in mouse brain. Cell annotation result is achieved through Harmony embedding and Leiden clustering, along with the UMAP plot. Scale bar: 1 mm. **b**, Pearson correlation of PRISM cell classification results with single-cell transcriptome data. **c**, Dot plot of each subtype.

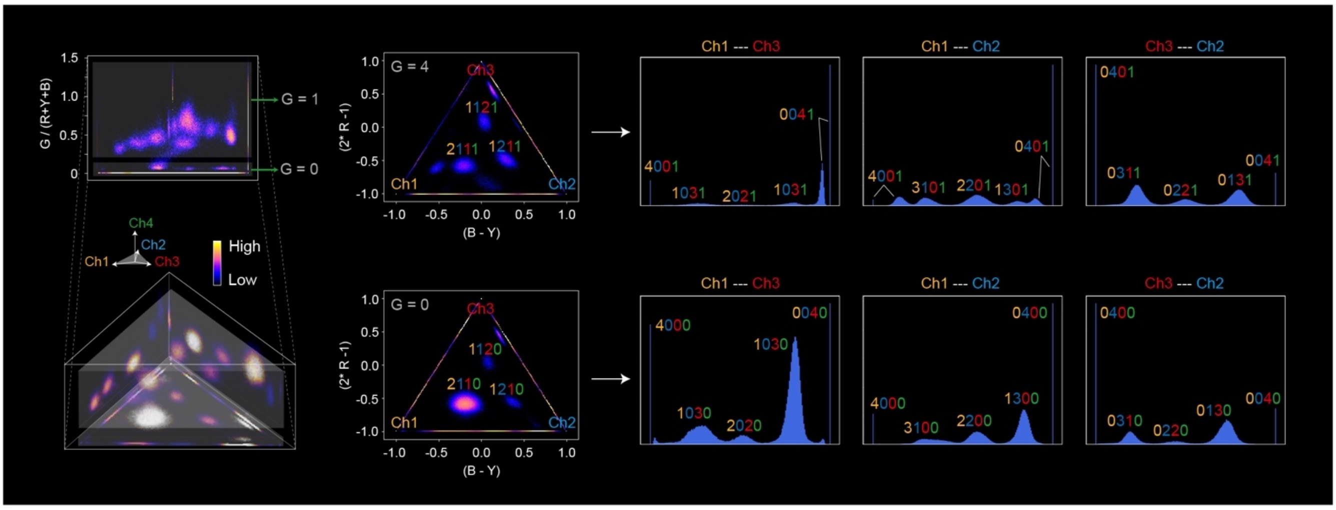


**Supplementary Fig. 12 | PRISM gene calling in color space for mouse embryo.** The distribution of the 30 barcode clusters can be observed through different cross-sections and projections.

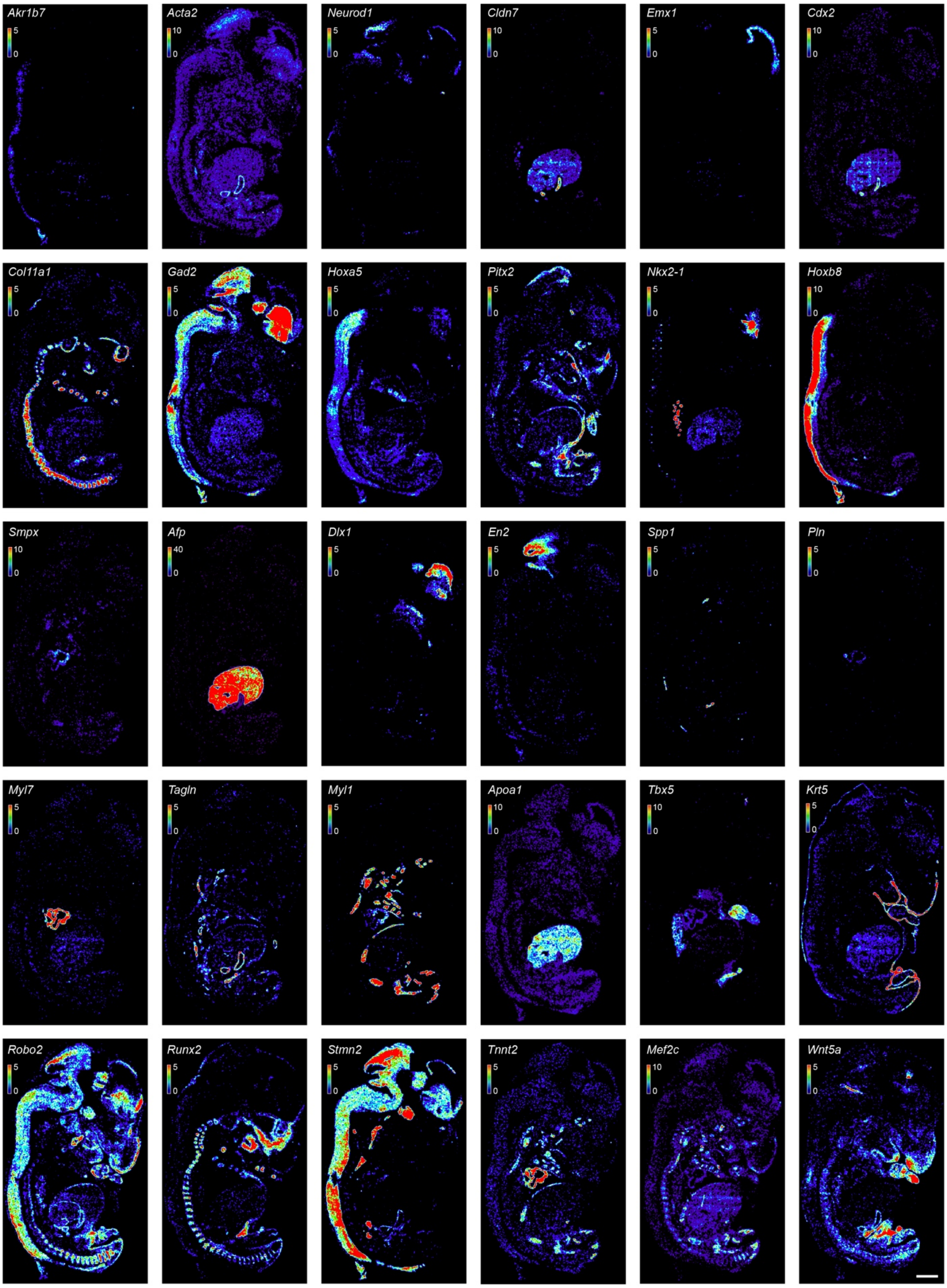


**Supplementary Fig. 13 | Spatial expression patterns of 30 called genes in mouse embryo E13.5.** For better visualization, the transcripts are down-sampled (coarse-grained). The dynamic range of transcripts density (counts per 100 x 100 $pixel^2$, 1 pixel = 0.1625 μm) is represented as shown in the colormap in each image. Scale bar: 1 mm.

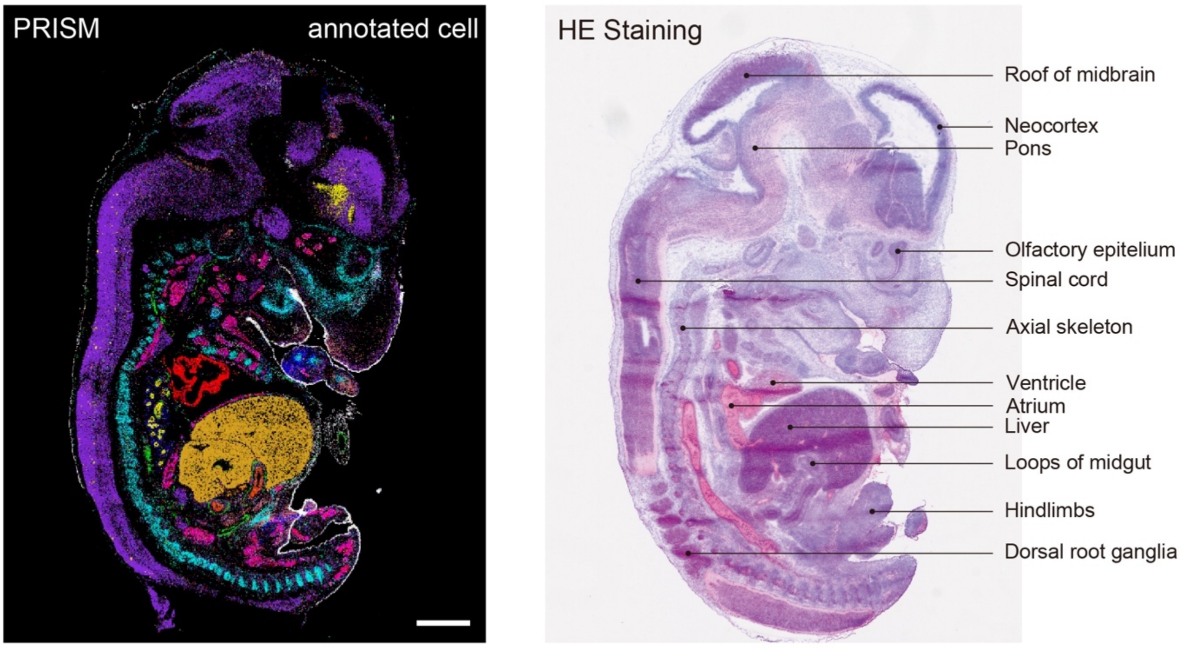


**Supplementary Fig. 14 | Benchmark comparison between PRISM result and HE staining result in adjacent section.** The PRISM experiment were repeated three times in adjacent sections with similar annotation results. Scale bar: 1 mm.

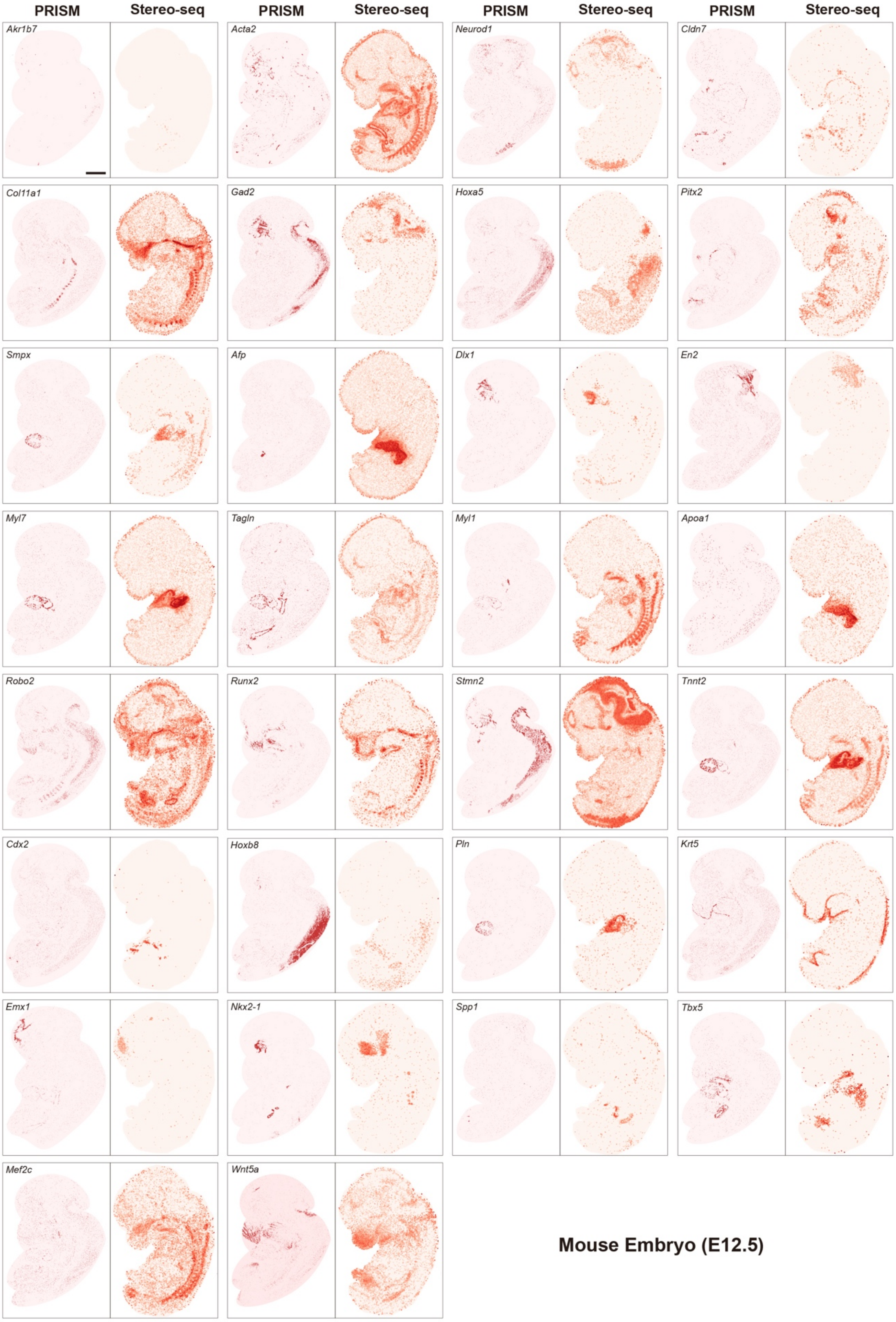
PRISM
Stereo-seq
PRISM
Stereo-seq
PRISM
Stereo-seq
PRISM
Stereo-seq
Akr1b7
Acta2
Neurod1
Cldn7
Col11a1
Gad2
Hoxa5
Pitx2
Smpx
Afp
Dlx1
En2
Myl7
Tagln
Myl1
Apoa1
Robo2
Runx2
Stmn2
Tnnt2
Cdx2
Hoxb8
Pln
Krt5
Emx1
Nkx2-1
Spp1
Tbx5
Mef2c
Wnt5a
Mouse Embryo (E12.5)

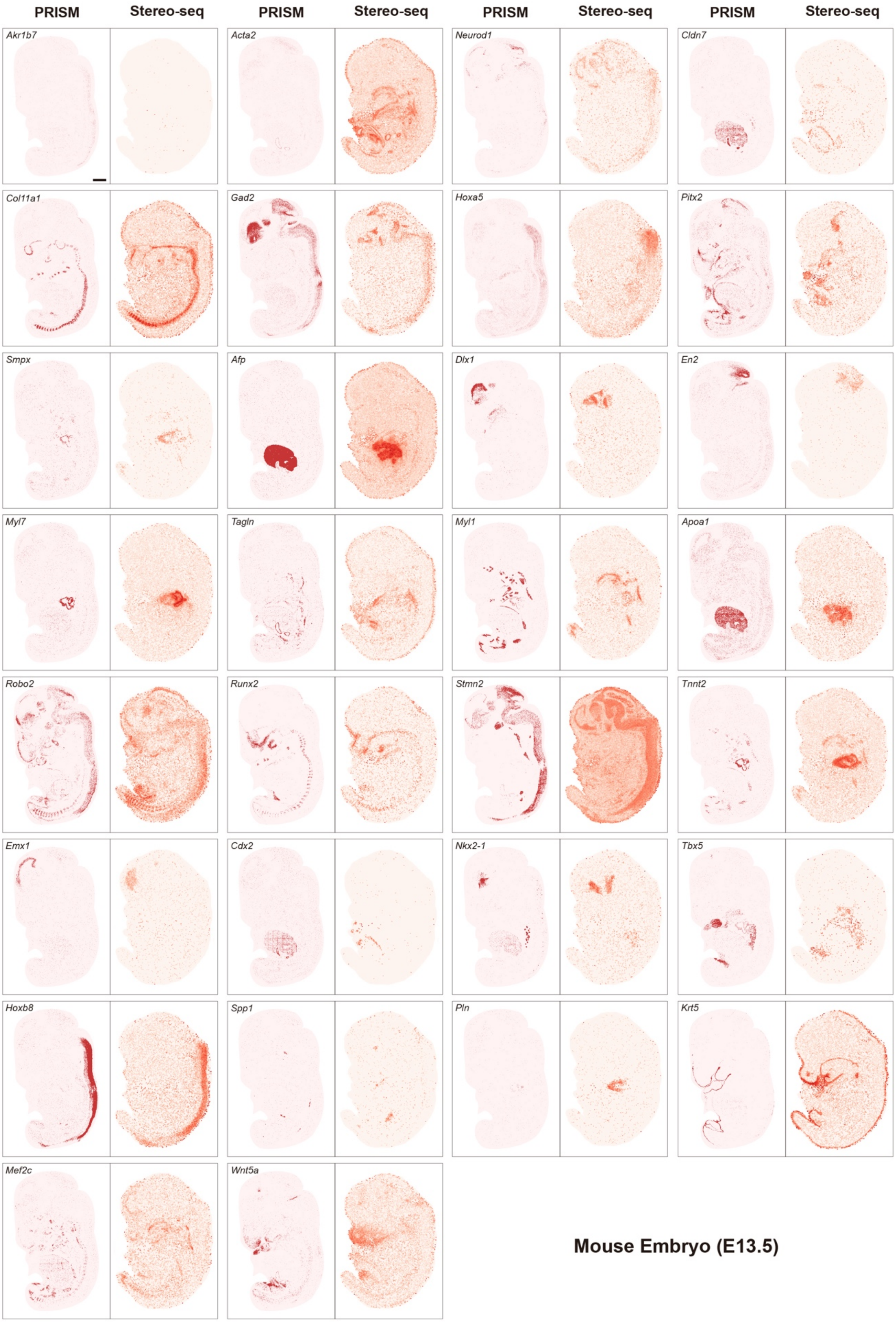
PRISM
Stereo-seq
PRISM
Stereo-seq
PRISM
Stereo-seq
PRISM
Stereo-seq
Akr1b7
Acta2
Neurod1
Cldn7
Col11a1
Gad2
Hoxa5
Pitx2
Smpx
Afp
Dlx1
En2
Myl7
Tagln
Myl1
Apoa1
Robo2
Runx2
Stmn2
Tnnt2
Emx1
Cdx2
Nkx2-1
Tbx5
Hoxb8
Spp1
Pln
Krt5
Mef2c
Wnt5a
Mouse Embryo (E13.5)

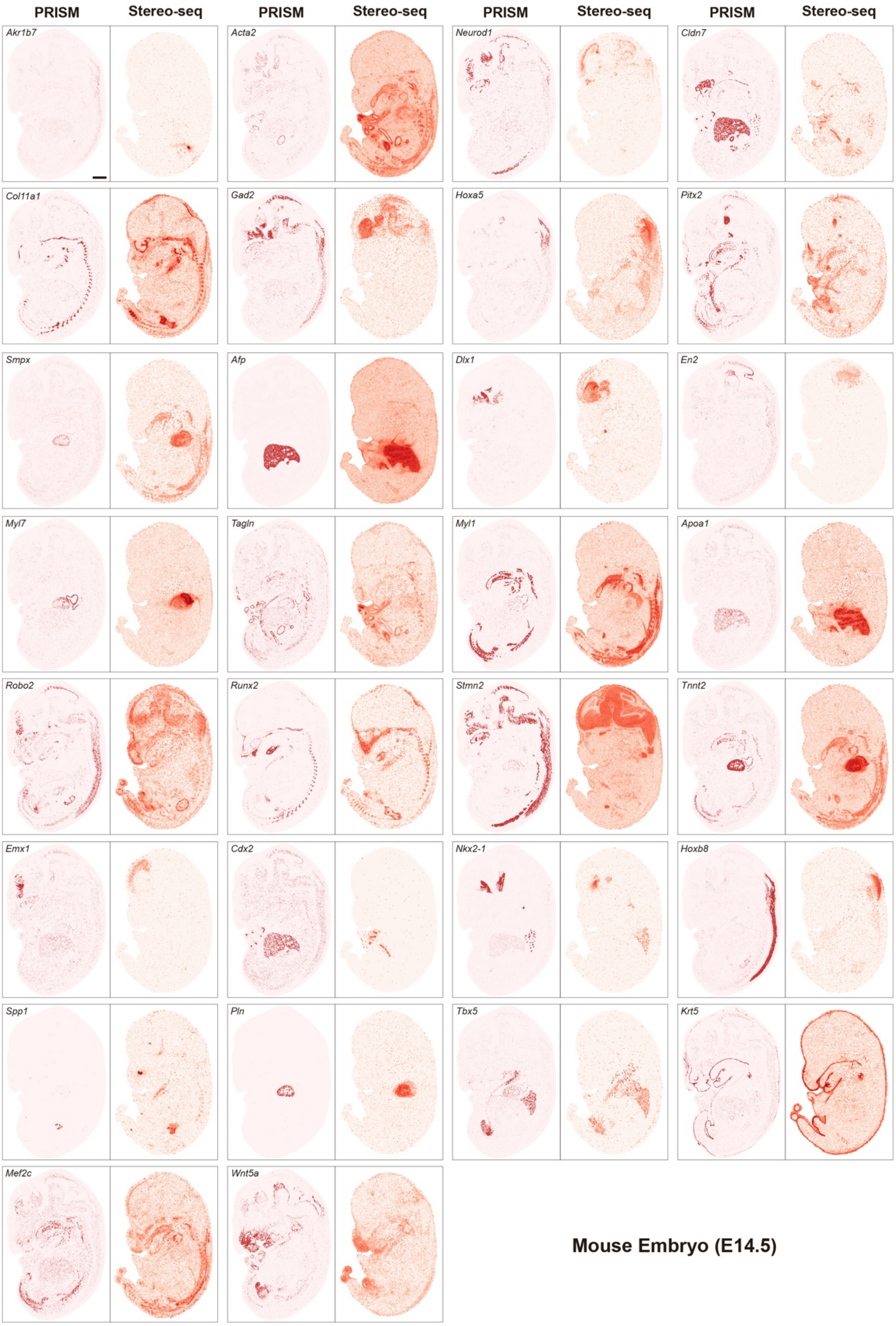


**Supplementary Fig. 15 | Spatial expression pattern comparison with MOSTA database (generated from stereo-seq).** This comparison was performed using sagittal sections of mouse embryos at E12.5-E14.5 stage. Selected data has a similar sectioning position (matched anatomical regions) as our data. Scale bar: 1 mm.

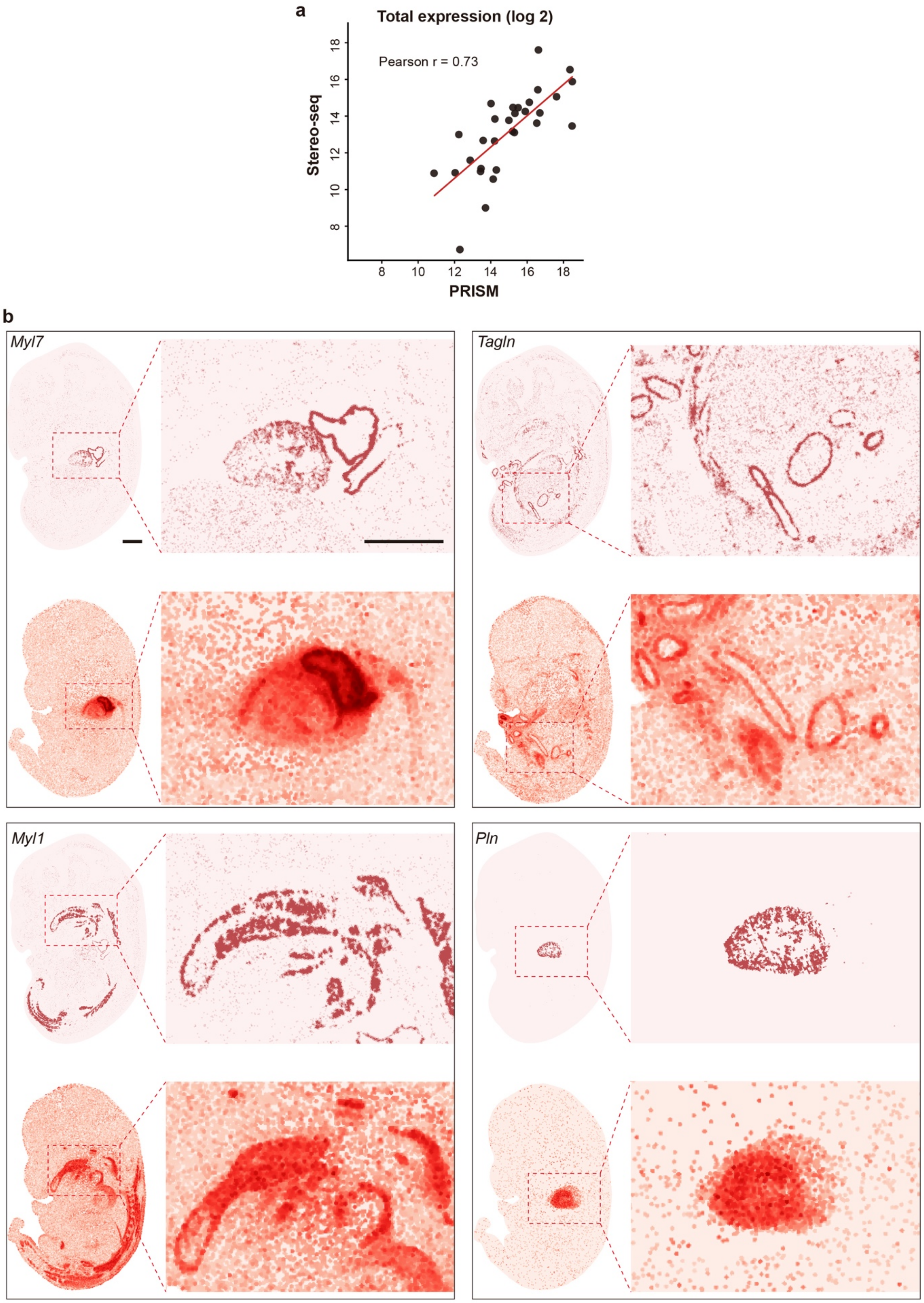


**Supplementary Fig. 16 | Total expression amount correlation and zoom-in comparison with MOSTA database. a**, total expression correlation. Pearson r is 0.73. **b**, Zoom-in spatial expression pattern comparison. Scale bar: 1 mm for both images. The PRISM total expression value in (a) and gene spatial expression pattern in (b) were repeated more than three times in adjacent sections with similar results.

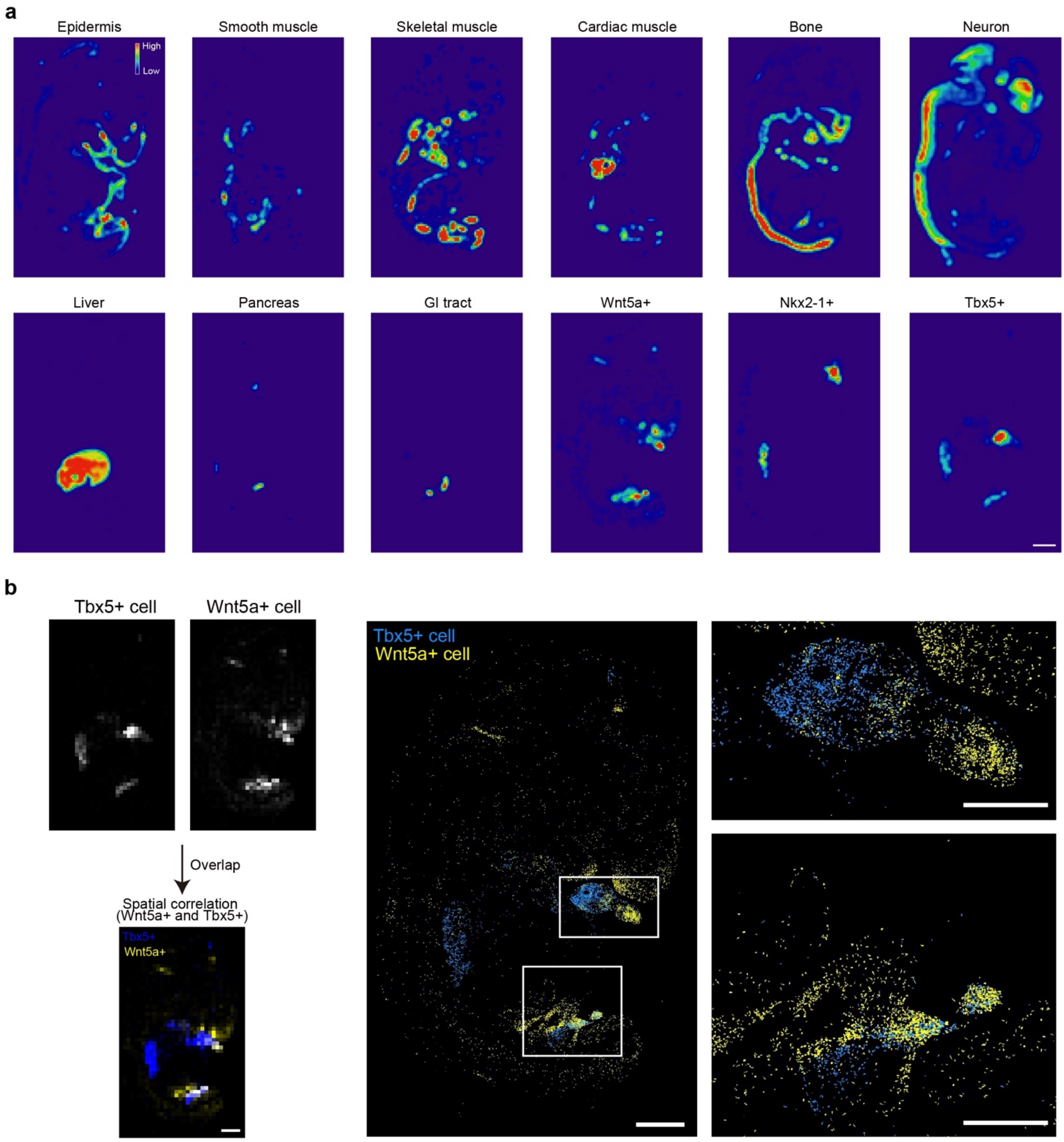


**Supplementary Fig. 17 | Cell type distribution and co-localization in mouse embryo. a**, Cell type distribution in mouse embryo E13.5. The cell density is represented as shown in the colormap. Scale bar: 1 mm. **b**, Spatial analysis of cell type co-localization. The left figure showed the spatial co-localization of 200 μm x 200 μm-binned Tbx5$^{+}$ cell (blue) and Wnt5a$^{+}$ cell (yellow), and zoom-in single cell resolution image on the right. The PRISM experiment was repeated for more than three times with similar cell distribution patterns. Scale bar: 1 mm, left; 500 μm, right.

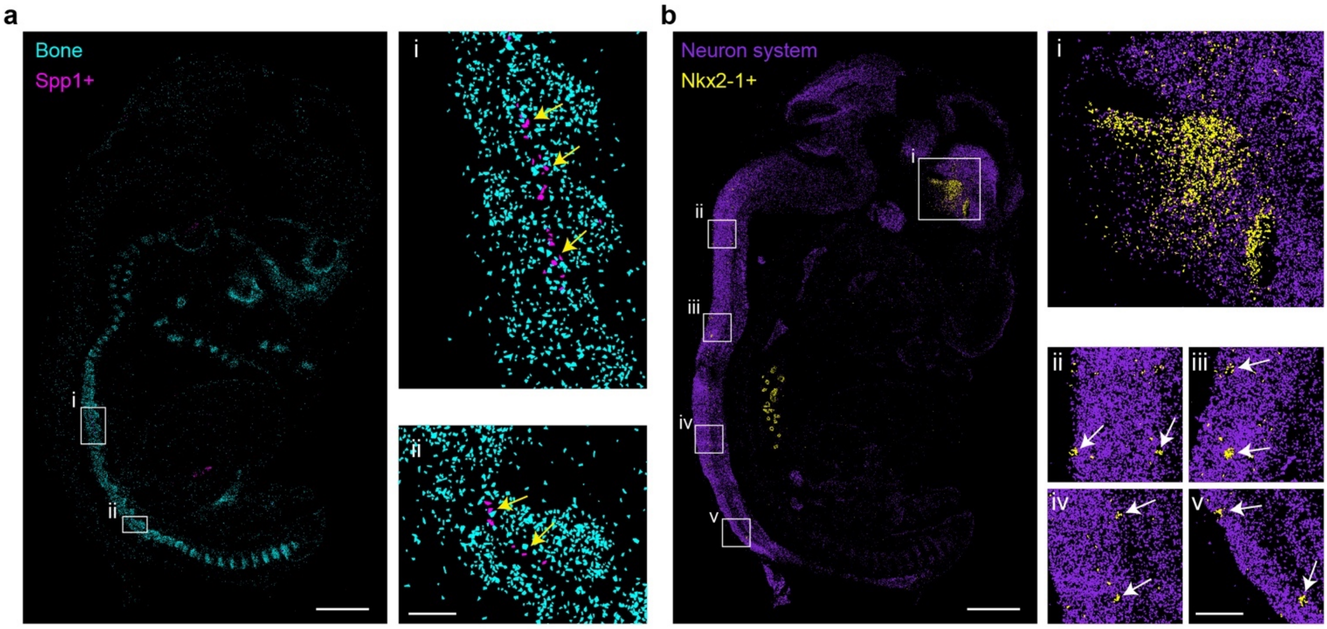


**Supplementary Fig. 18 | Fine intercellular-interaction in mouse embryo. a**, Interaction between Spp1$^{+}$ cells (shown in magenta) and bone cell (shown in cyan). Scale bar: 1mm, left; 100 μm, right. **b**, Interaction between Nkx2-1$^{+}$ cells (shown in yellow) and nervous system (shown in violet). Scale bar: 1 mm, left; 200 μm, right. The PRISM experiment was repeated for more than three times with similar cell distribution patterns.

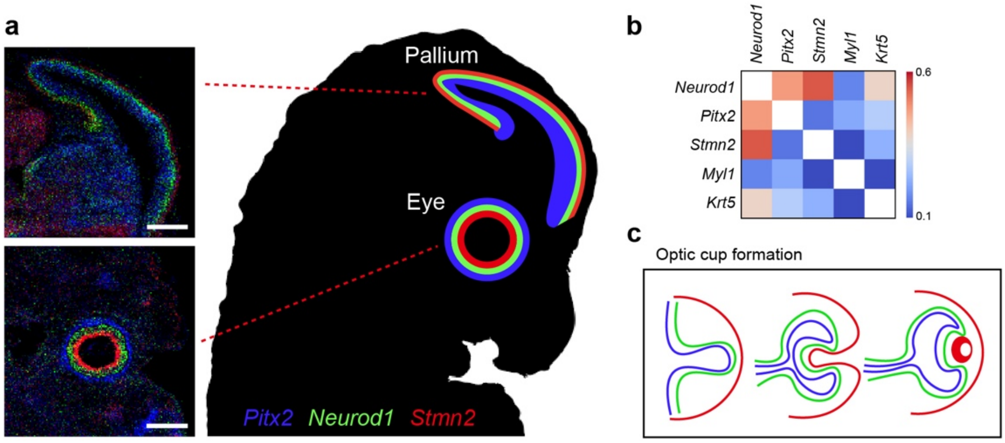


**Supplementary Fig. 19 | Layer inversion in optic cup and pallium reveals neural layer turnover during development.** **a**, Three neural layers marked by *Pitx2* (blue), *Neurod1* (green) and *Stmn2* (red) have an “inversed” pattern in optic cup and pallium. Scale bar: 500 μm. **b**, Spatial correlation between genes in the nervous system. The ‘*Neurod1*’ layer is situated between the ‘*Pitx2*’ layer and ‘*Stmn2*’ layer. **c**, Hypothesis of optic cup formation to explain the layer “turnover” during development.

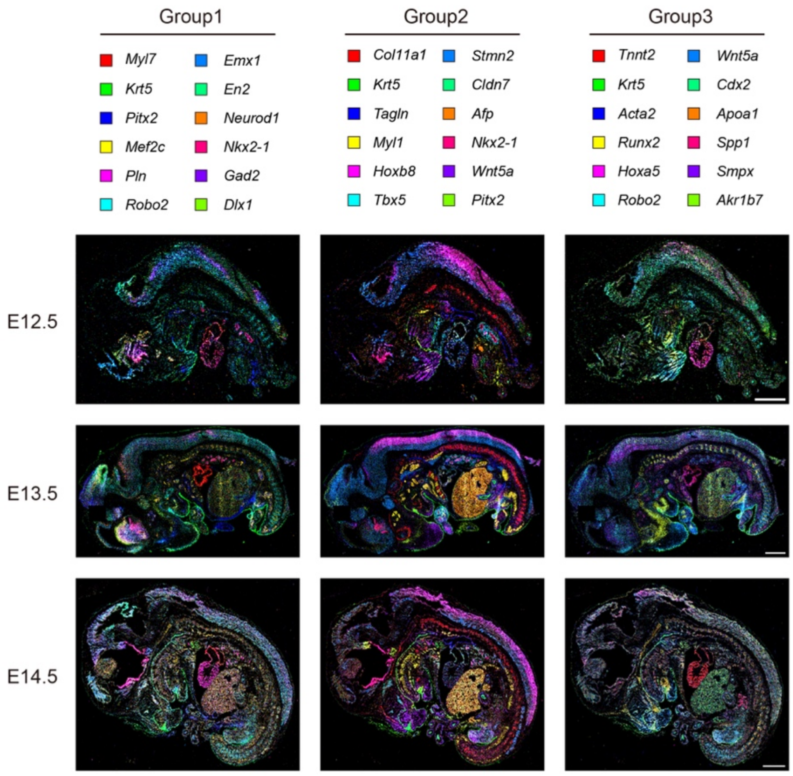


**Supplementary Fig. 20 | Demonstration of various organ structures in mouse embryo at E12.5, E13.5 and E14.5 through gene expression.** Genes were put in different group to emphasize different organs. Scale bar: 1 mm.

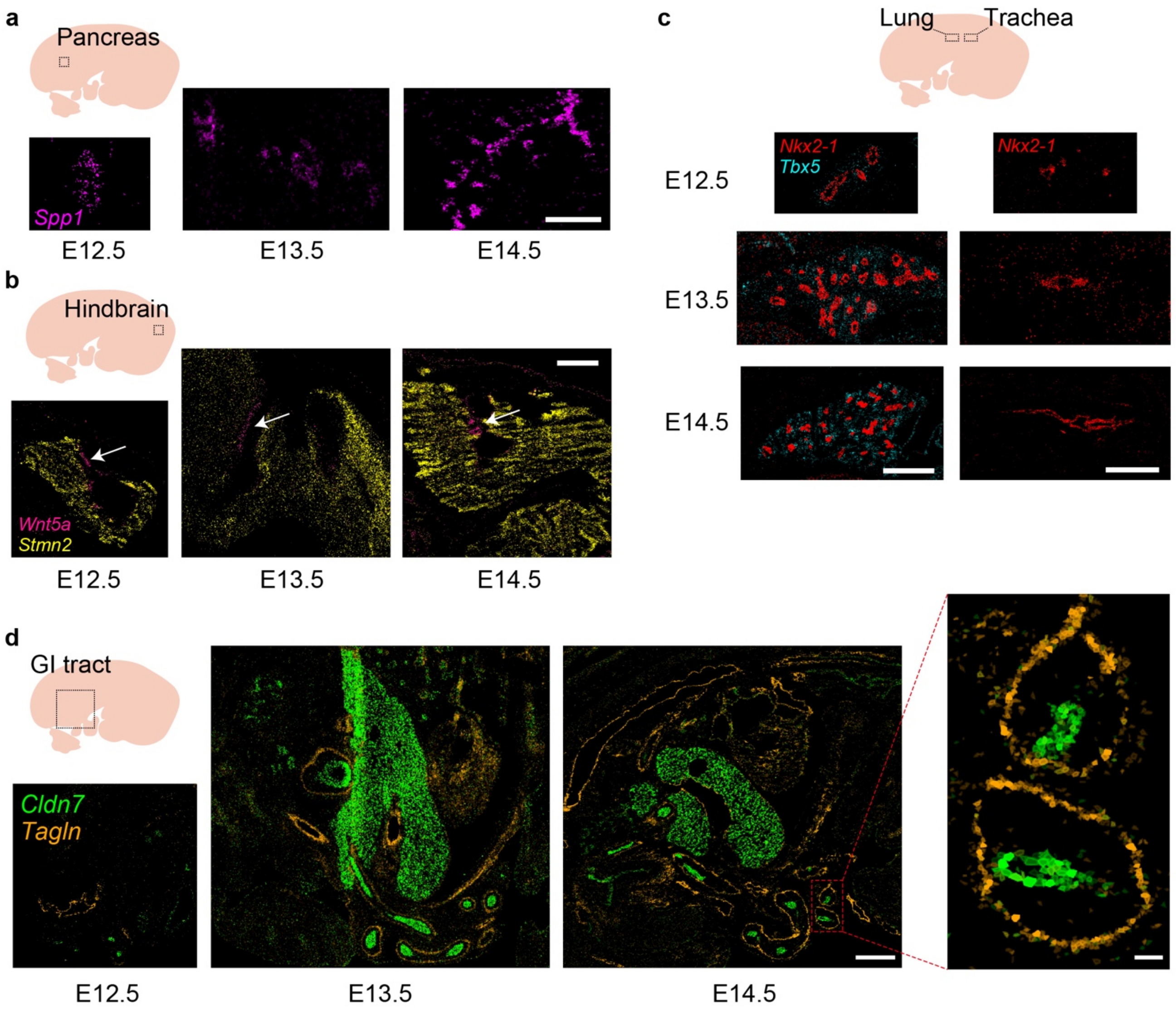


**Supplementary Fig. 21 | Temporal developmental changes at cellular level in mouse embryo. a**, Scale bar: 250 μm. **b-c**, Scale bar: 500 μm. **d**, Scale bar: 500 μm for left; 50 μm for zoom-in image. The cells were colored by gene expression and abundance. The PRISM experiment was repeated for more than three times with similar cell distribution patterns.

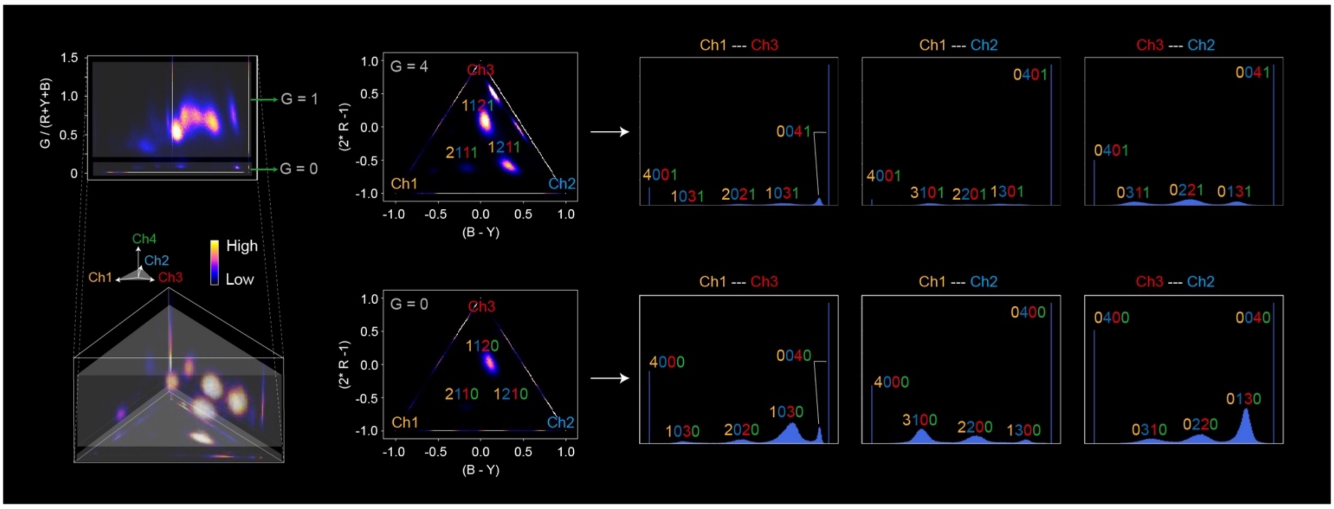


**Supplementary Fig. 22 | PRISM gene calling in color space for human hepatocellular carcinoma tumor (HCC) tissue.** The distribution of the 31 barcode clusters can be observed through different cross-sections and projections.

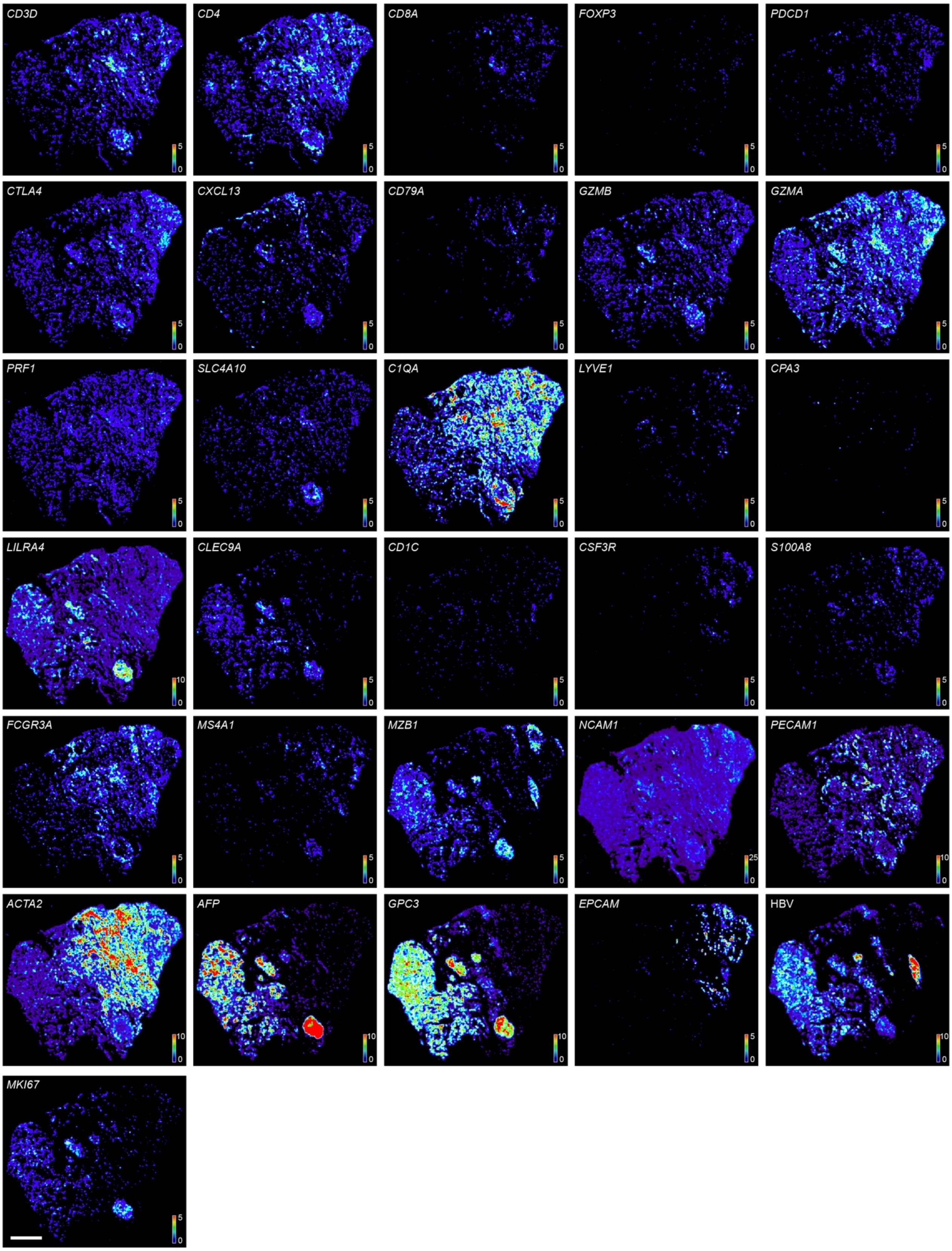


**Supplementary Fig. 23 | Spatial expression patterns of 31 called genes in HCC.** For better visualization, the transcripts are down-sampled (coarse-grained). The dynamic range of transcripts density (counts per 100 x 100 pixel$^2$, 1 pixel = 0.1625 μm) is represented as shown in the colormap in each image. Scale bar: 1 mm.

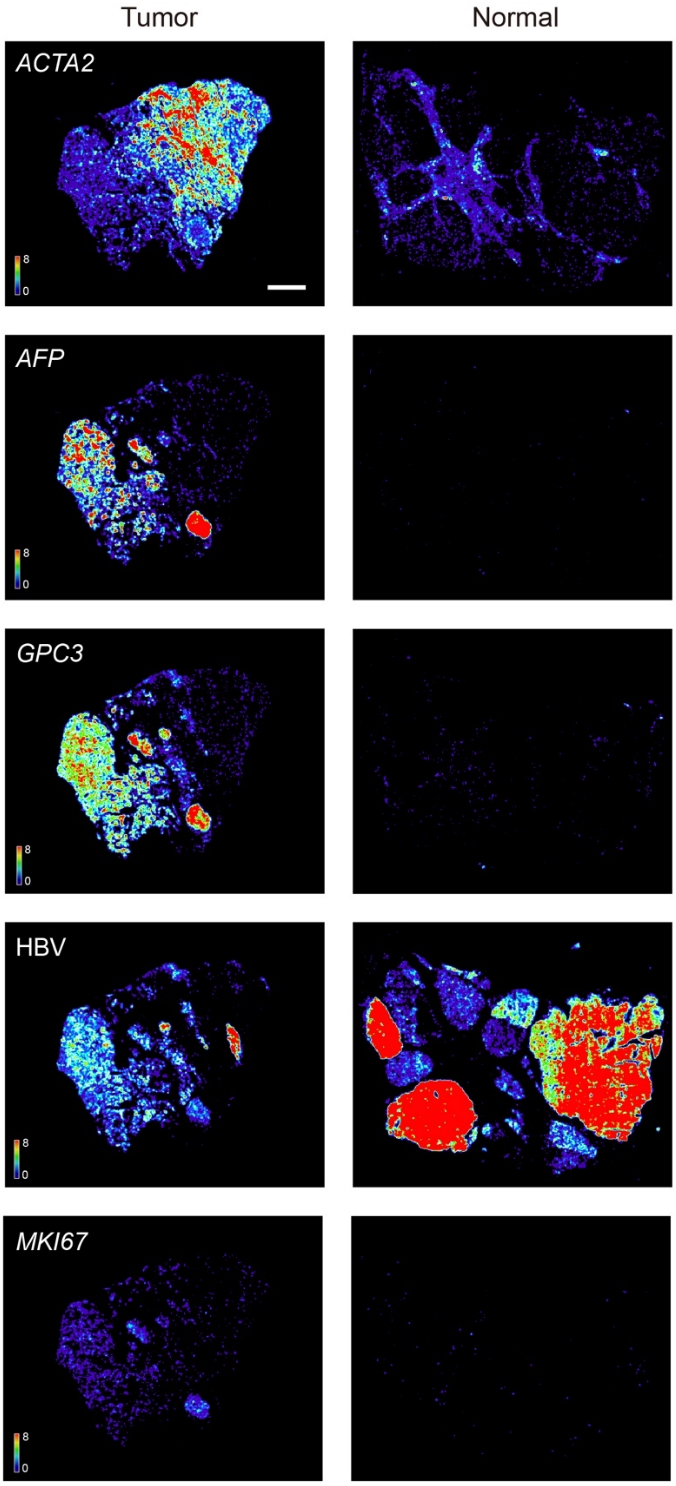


**Supplementary Fig. 24 | Comparison of selected gene spatial expression (by PRISM) between tumor and normal tissue.** Scale bar: 1 mm.

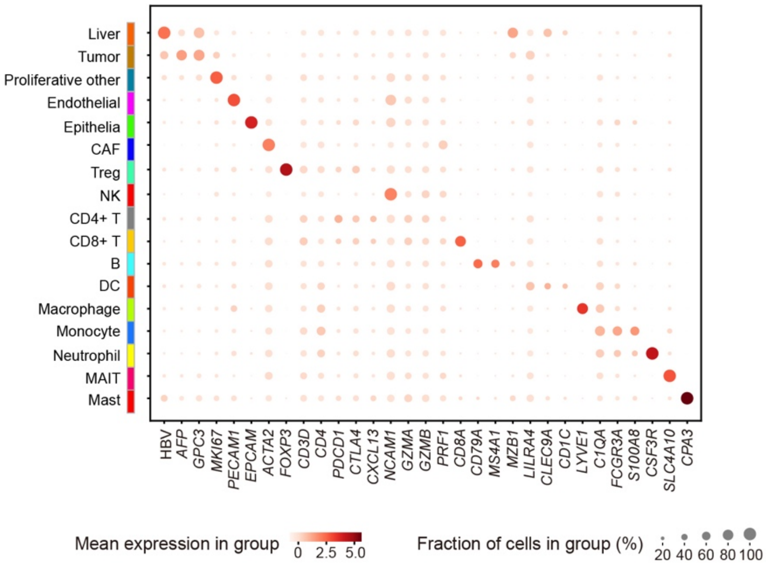


**Supplementary Fig. 25 | Dot plot of general cell types classified in HCC.**

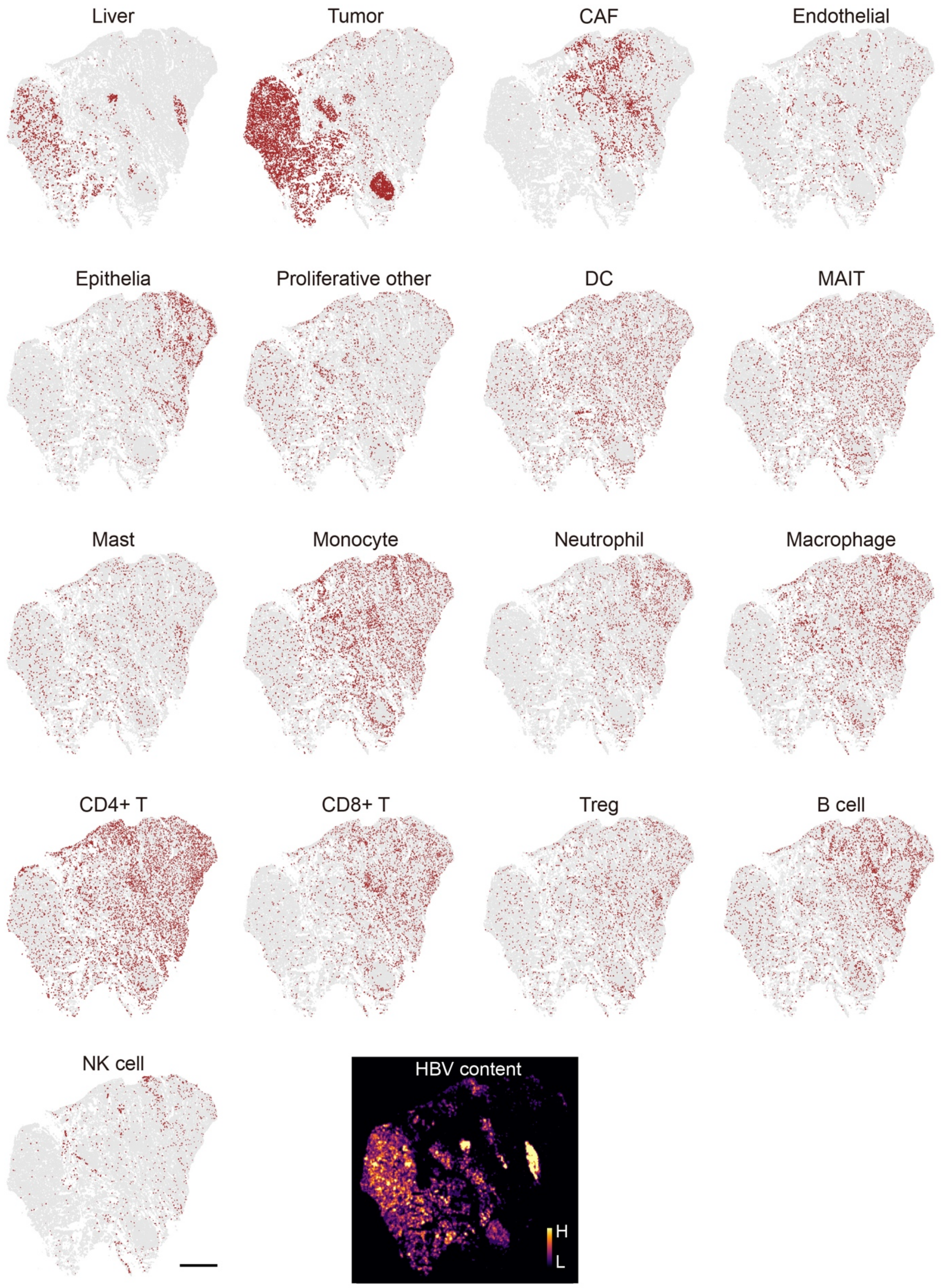


**Supplementary Fig. 26** | **General cell type and HBV distribution in HCC.** Scale bar: 1 mm.

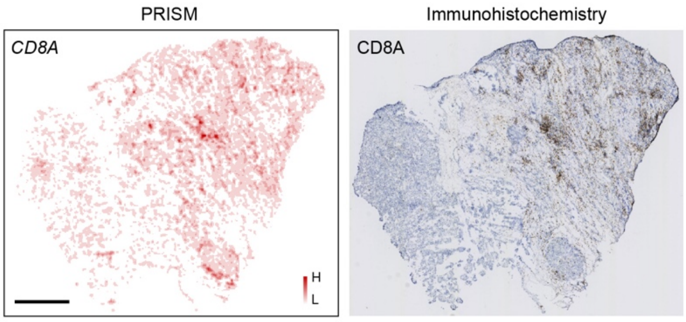


**Supplementary Fig. 27 | Comparison of PRISM RNA result against immunohistochemistry result in an adjacent section (CD8A).** The PRISM experiment was repeated for three times in adjacent sections with similar gene spatial expression patterns. Scale bar: 1 mm.

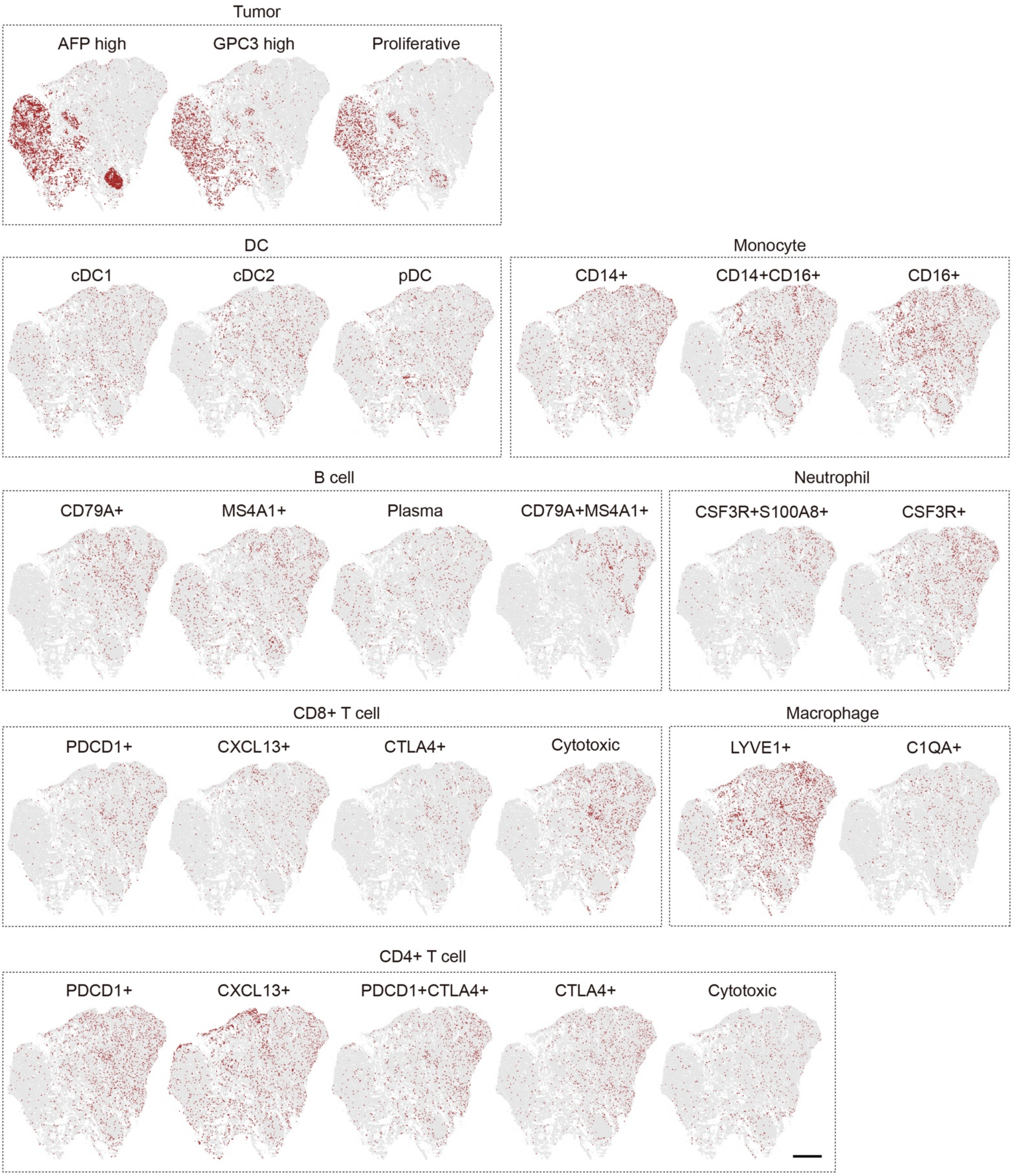


**Supplementary Fig. 28 | All cell type distribution in HCC.** Scale bar: 1 mm.

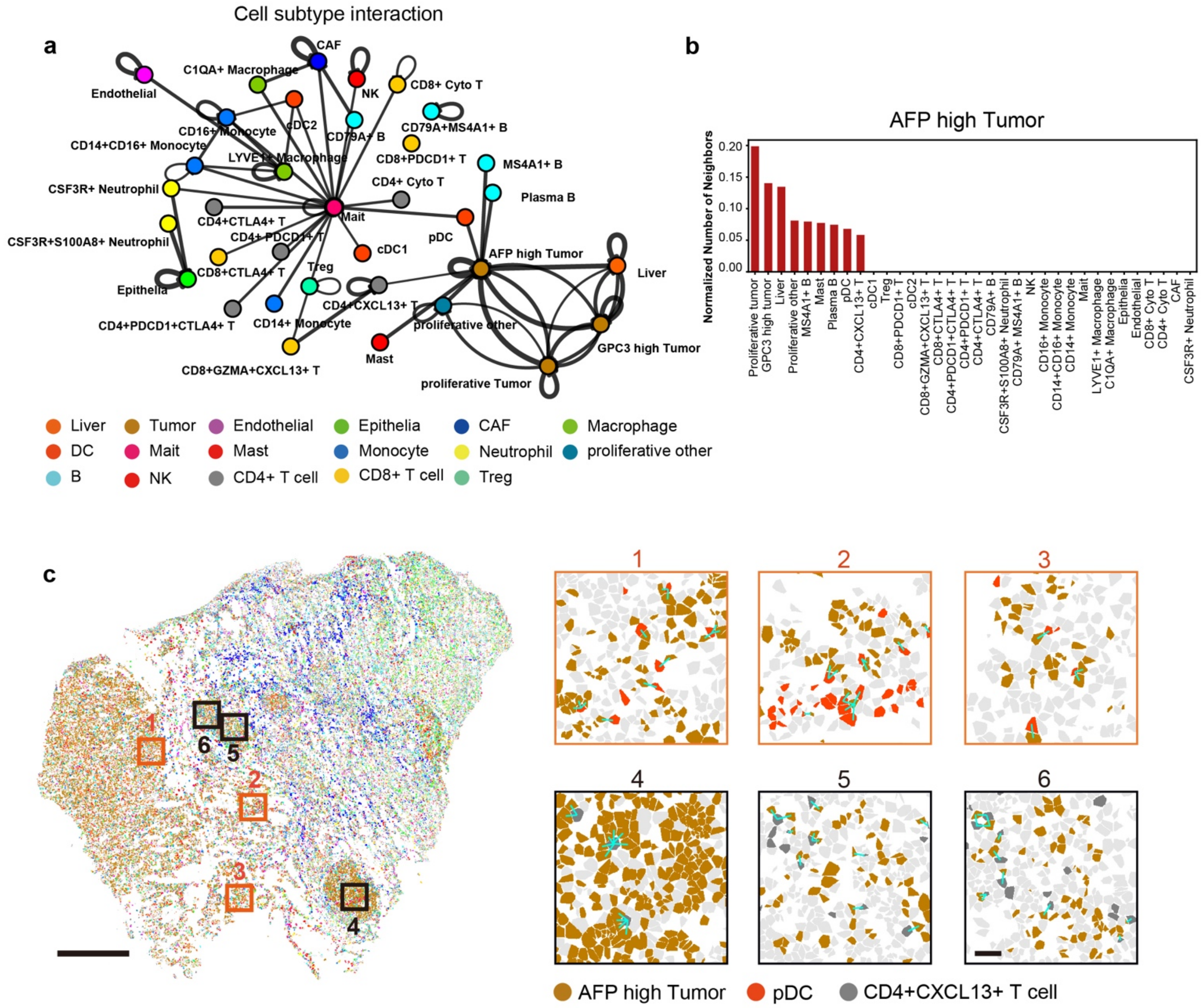


**Supplementary Fig. 29 | Cell subtype interactions in HCC. a**, Graph visualization of Squidpy neighbor interaction analysis reveals interactions among cell subtypes in HCC. **b**, Normalized counts of neighboring cells in proximity to AFP high tumor cells. **c**, Representative regions showing colocalization of AFP high tumor cells with pDCs or CD4+CXCL13+ T cells. Scale bar: 1 mm, left (overview); 10 μm, right (zoom-in images).

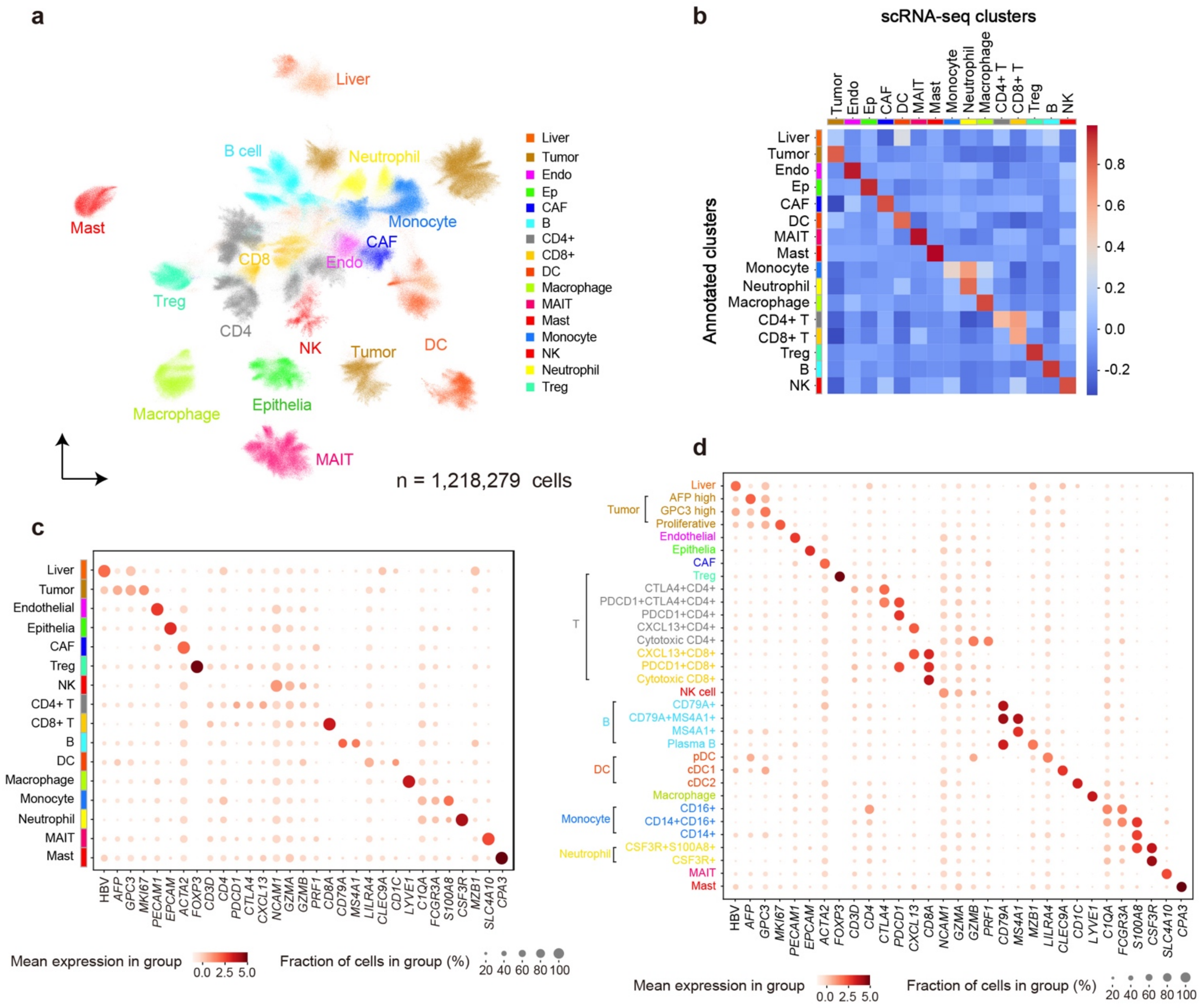


**Supplementary Fig. 30 | Cell classification in cells from 20-consecutive HCC sections. a**, UMAP plot. **b**, Pearson correlation between PRISM classification result and single cell transcriptome data. **c**, Dot plot of general cell types. **d**, Dot plot of all cell types.

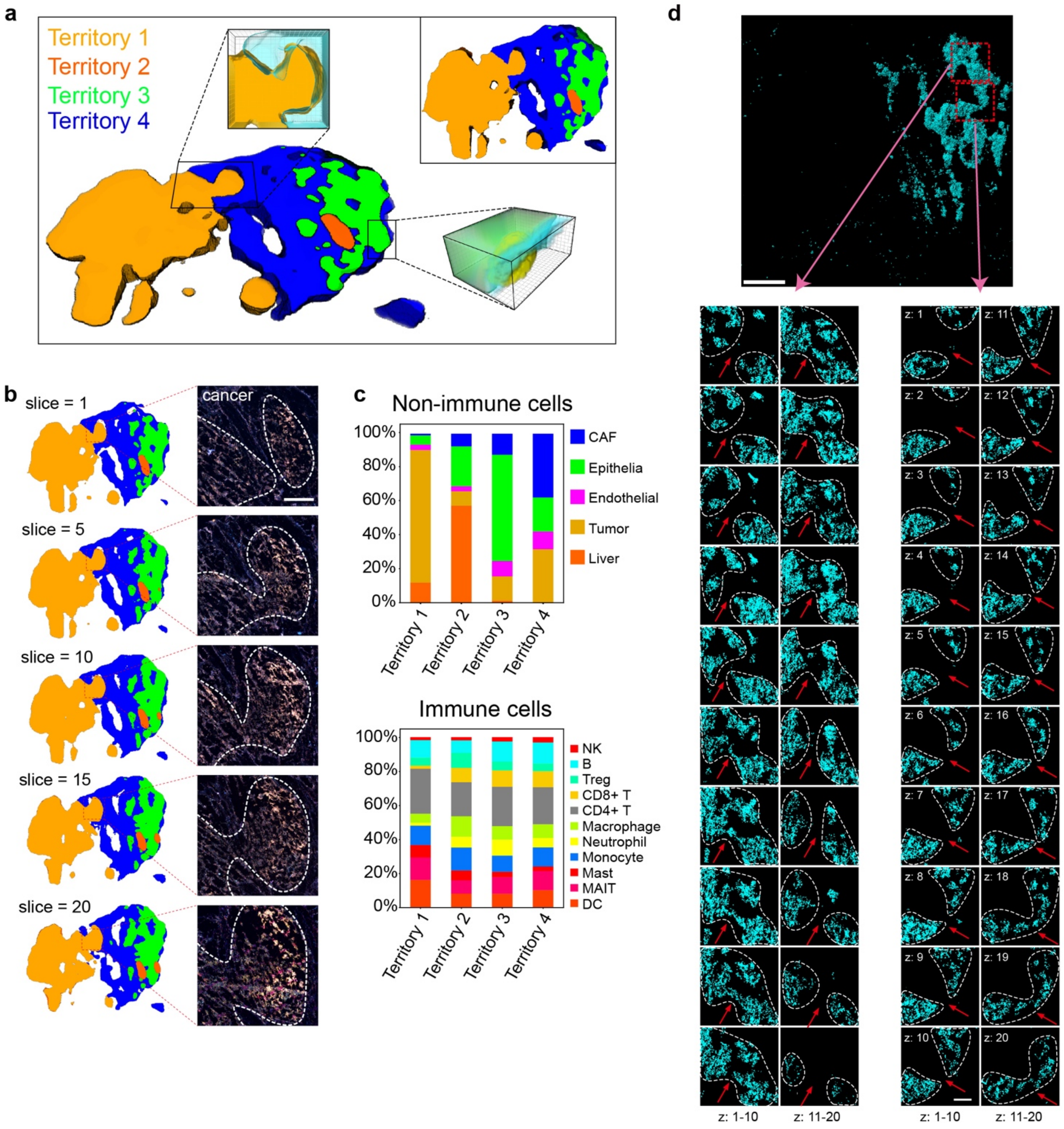


**Supplementary Fig. 31 | Projection of 20 slices based on regions. a**, 3D territory segmented based on gene distribution: tumor (Territory 1), normal liver (Territory 2), epithelia (Territory 3), and CAFs (Territory 4). **b**, Sections in 3D territory. The right part shows the raw image in a tumor-protrusion region. Scale bar: 300 μm. **c**, Percentages of cells in four territories. **d**, Sections of the 3D interconnected B-cell network. The 20 consecutive slices below (zoomed from the red dashed box in the overview) illustrate examples, with red arrows marking connected or disconnected regions within the network. Scale bars: 1 mm, overview; 200 μm, below.

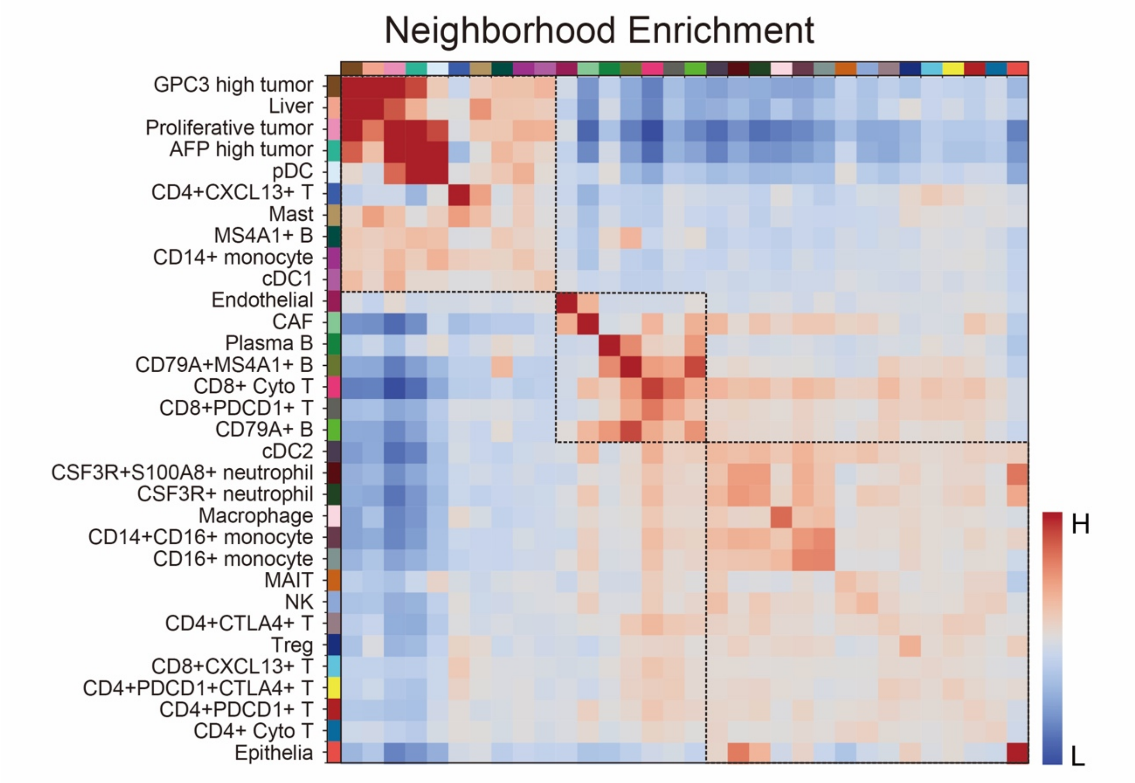


**Supplementary Fig. 32 | Neighborhood enrichment of HCC pseudo analysis.**

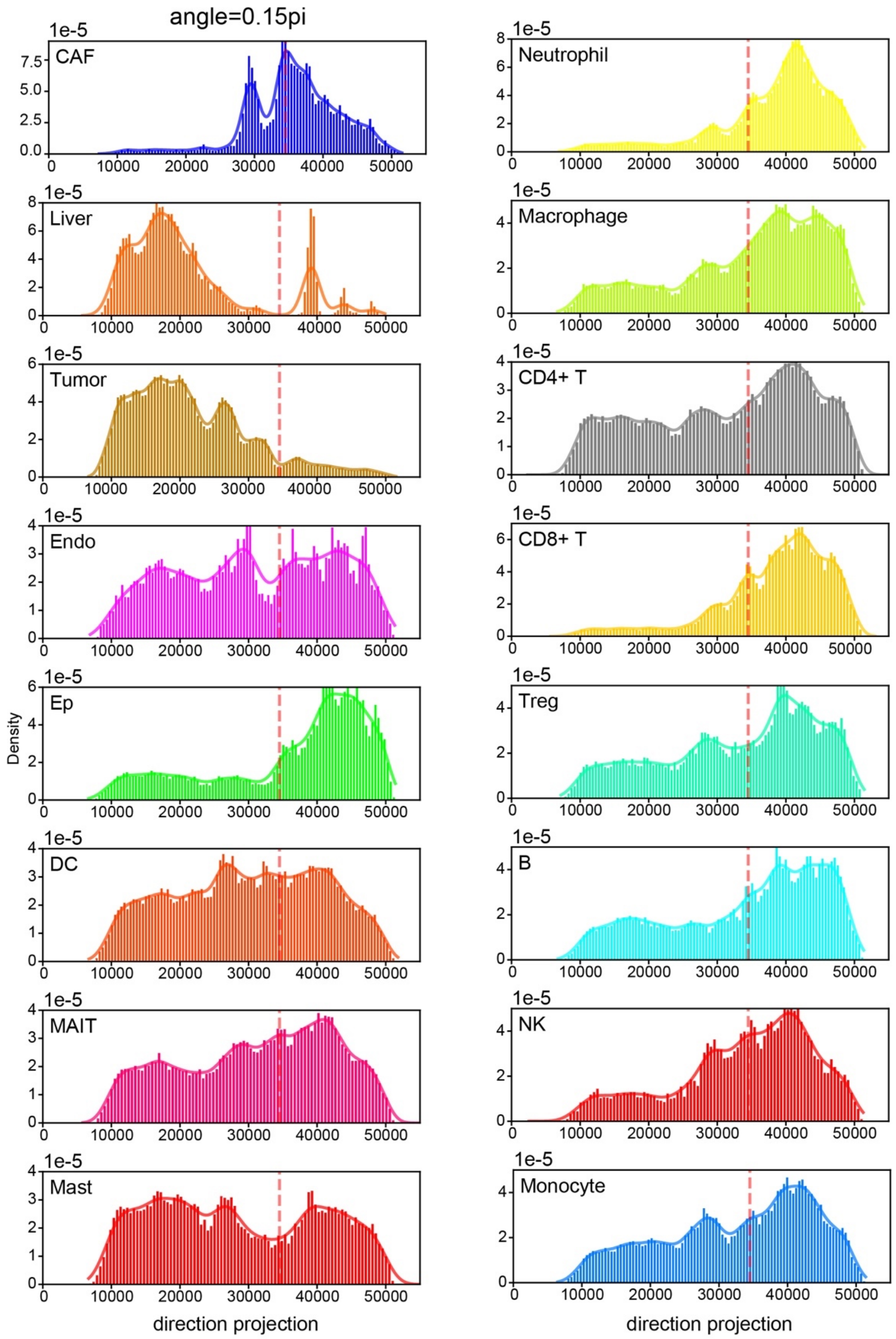


**Supplementary Fig. 33 | One-dimension distribution of cells after projection along the long axis of tumor (vertical to CAF wall).** Red dash line shows the position of CAF wall.

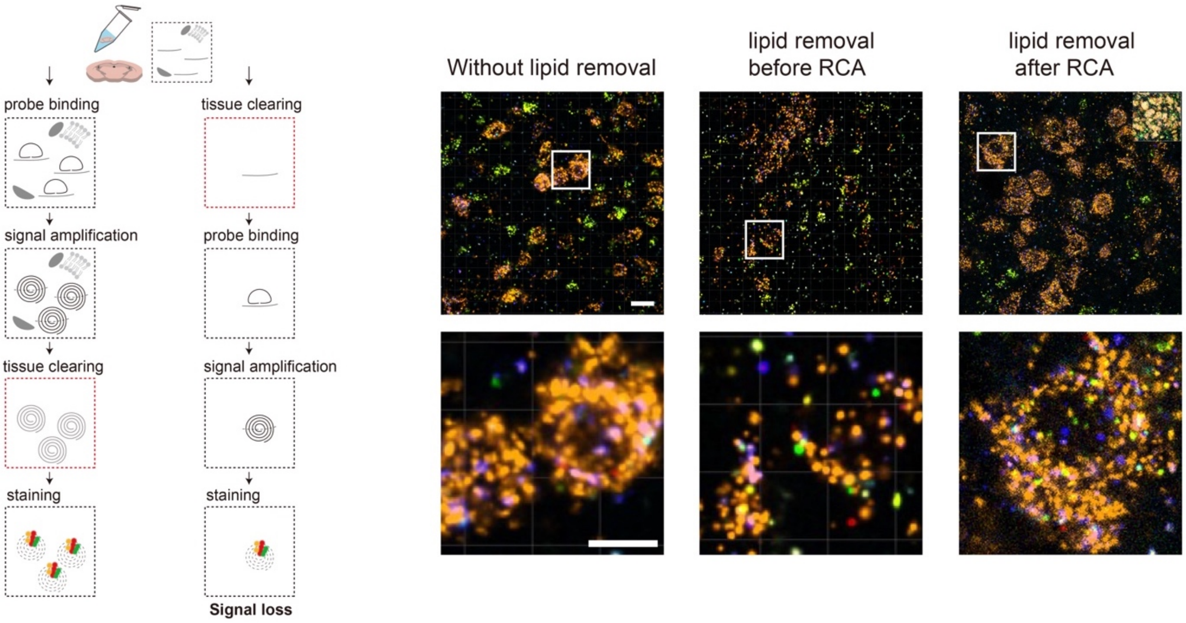


**Supplementary Fig. 34 | Lipid removal in thick tissue.** Removing lipids is necessary to improve optical transparency, performing this step before rolling circle amplification may result in a decrease in the signal count. This assessment was repeated three times with similar results. This decline could be caused by the loss of RNAs that were initially fixed within membrane proteins, which were removed during lipid removal. Scale bar: 20 μm, overview; 10 μm, zoom-in image.

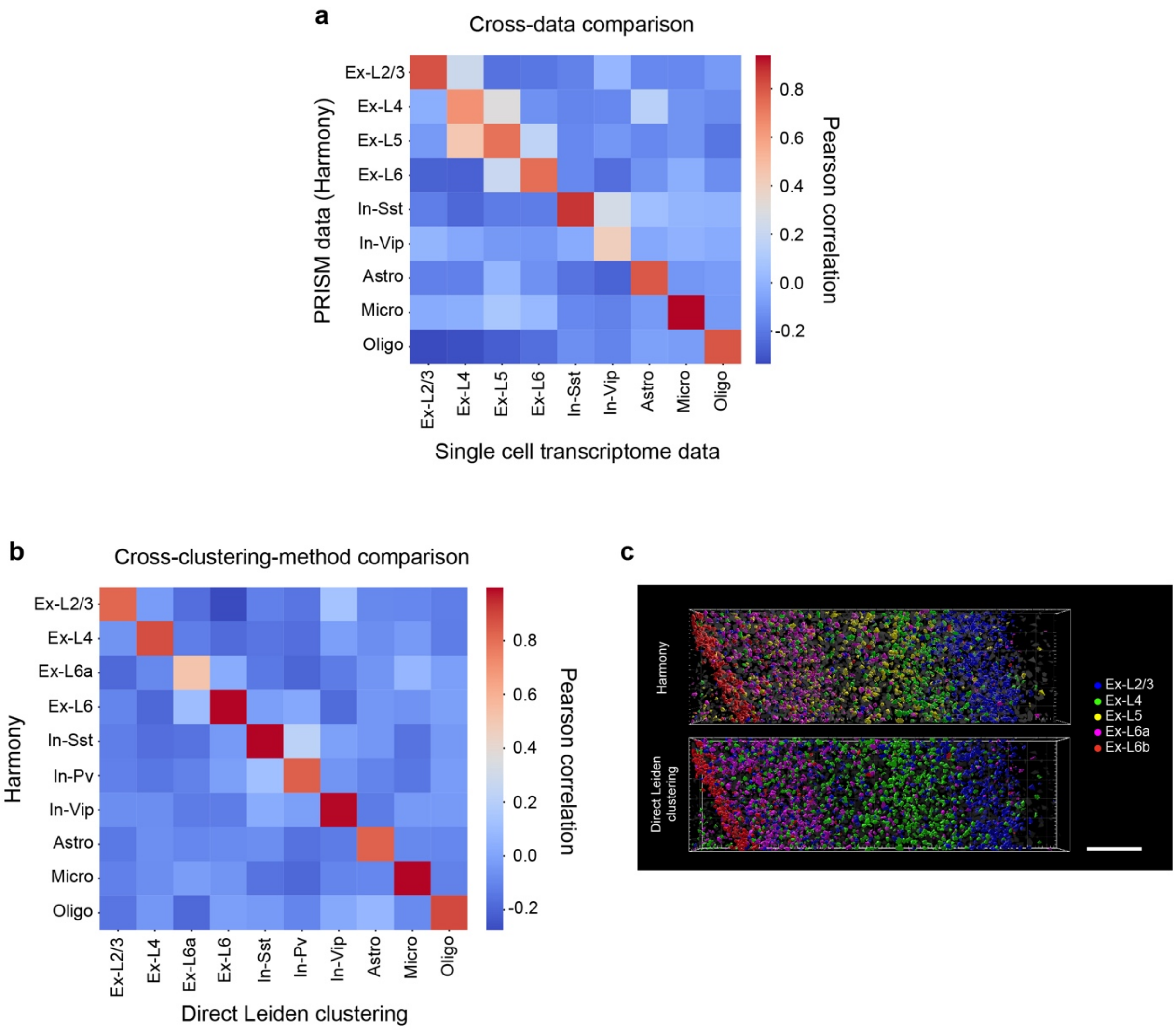


**Supplementary Fig. 35 | Cross method comparison in 3D mouse brain expression data. a**, Pearson correlation between PRISM 3D expression data and single cell transcriptome data. **b**, Pearson correlation between Harmony classification and direct Leiden classification (for PRISM data). **c**, The cell type distribution result from Harmony classification and direct Leiden classification. Scale bar: 200 μm.

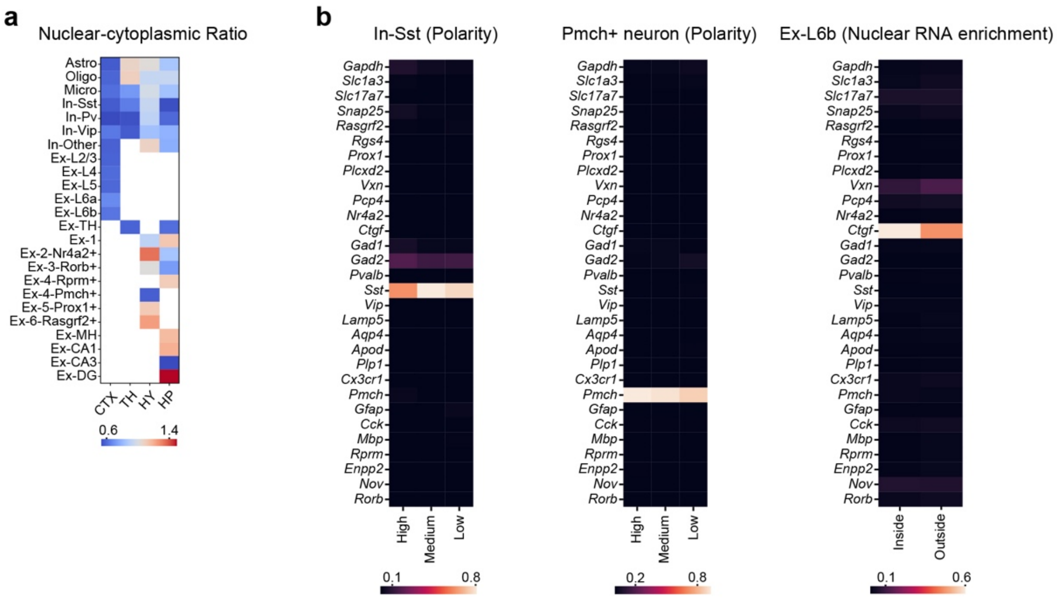


**Supplementary Fig. 36 | Subcellular RNA distribution across different cell type and brain region. a**, Nuclear-cytoplasmic ratio. **b**, Gene contribution in subcellular analysis. The subcellular analysis results of each cell type were predominantly influenced by its marker gene, as these genes had a high weighting in the expression abundance in corresponding cell types.

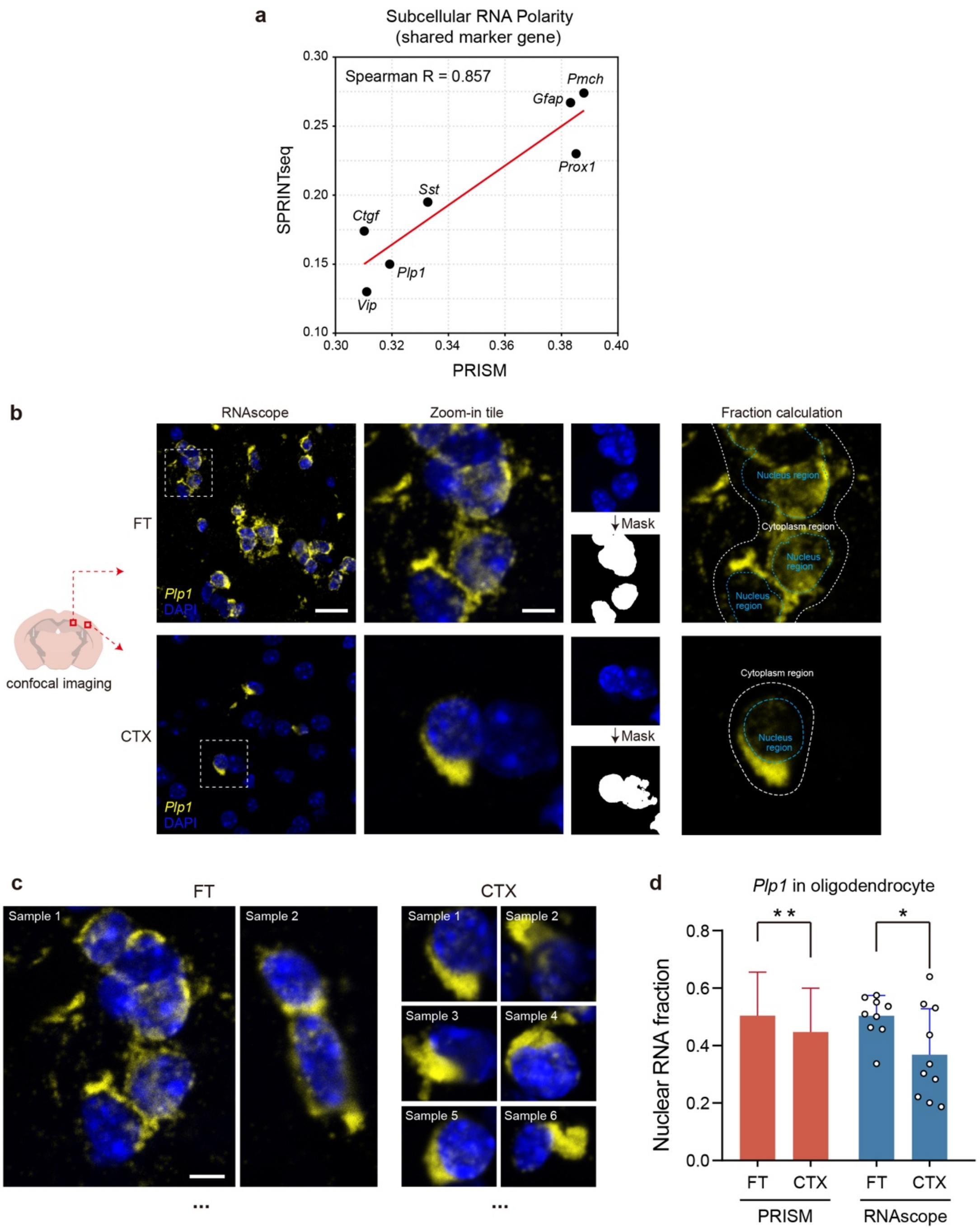


**Supplementary Fig. 37 | Validation of subcellular RNA distribution using existing datasets and RNAscope. a**, Validation of RNA polarity by benchmarking PRISM data against published in situ sequencing (ISS) data. Polarity values were calculated as the normalized distance between transcript centroids and nuclear centroids. Shared marker genes present in both datasets were used for comparison. **b**, Validation of subcellular RNA distribution (RNA enrichment within nucleus) difference across brain regions. *Plp1*, a highly and predominantly expressed gene in oligodendrocytes included in our PRISM detection panel, was selected for RNAscope validation. Confocal imaging was performed to simultaneously capture *Plp1* RNA signals and DAPI staining. Nuclear regions were delineated using DAPI masks (blue dashed lines). Enrichment was quantified as the ratio of total intensity (RNA channel) inside the nuclear region (area within blue dashed lines) to that in the whole cell region (area within white dashed lines). Scale bar: 20 μm for left; 10 μm for right. **c**, Representative RNAscope images of *Plp1* gene expression across brain regions. RNA signal was shown in yellow and DAPI was shown in blue. Scale bar: 10 μm. **d**, Comparative quantification of nuclear RNA enrichment between brain regions by PRISM and RNAscope. Nuclear RNA fractions were measured in annotated oligodendrocytes (PRISM, n = 460 for FT and 77 for CTX) and in 10 randomly selected fields per region (RNAscope). Both methods indicate higher nuclear RNA enrichment in FT (HP) region compared to CTX region. Data shown as mean ± s.d. Two-tailed t-test. Statistical significance: PRISM, p = 0.002; RNAscope, p = 0.025.

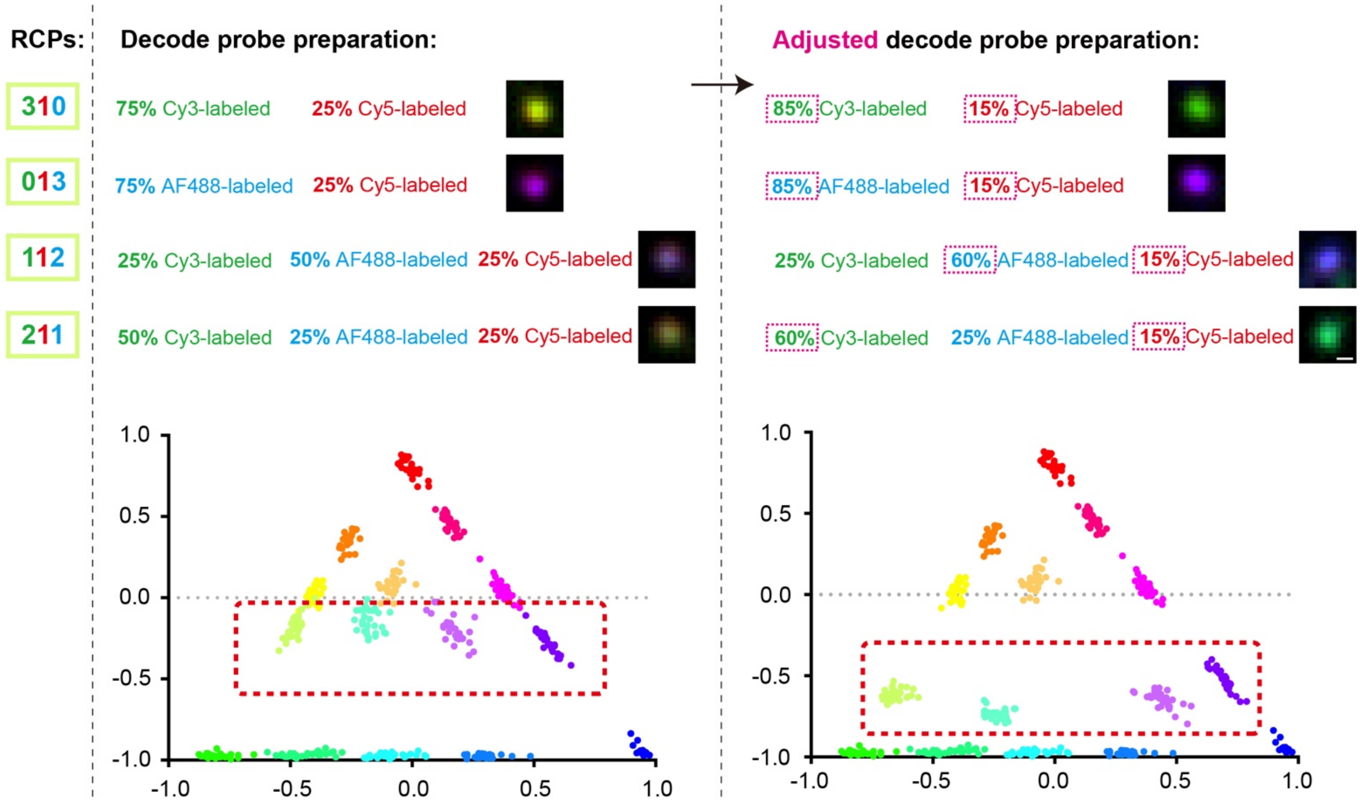


**Supplementary Fig. 38 | Adjustability of PRISM encoding.** Each fluorescent level can be fine-tuned by adjusting the mixing ratio of fluorescent probes to non-fluorescent probes corresponding to each barcode segment in the padlock probes. This adjustability ensures a robust barcode decoding using almost any multi-color imaging instrument, which may have its specific color space. In the first trial (shown left), we assumed a linear relationship between the proportion of fluorophore labeled decode probes and the resulting fluorescent intensity ratio. Certain intensity level clusters exhibit close proximity to adjacent clusters in color space, resulting in a low distinguishability of these barcodes (clusters). Through empirical optimization (reducing Cy5-labeled probes from 25% to 15% for ‘Ch2-level 1’ ), all clusters can be well separated in color space (shown right). Scale bar: 500 nm.

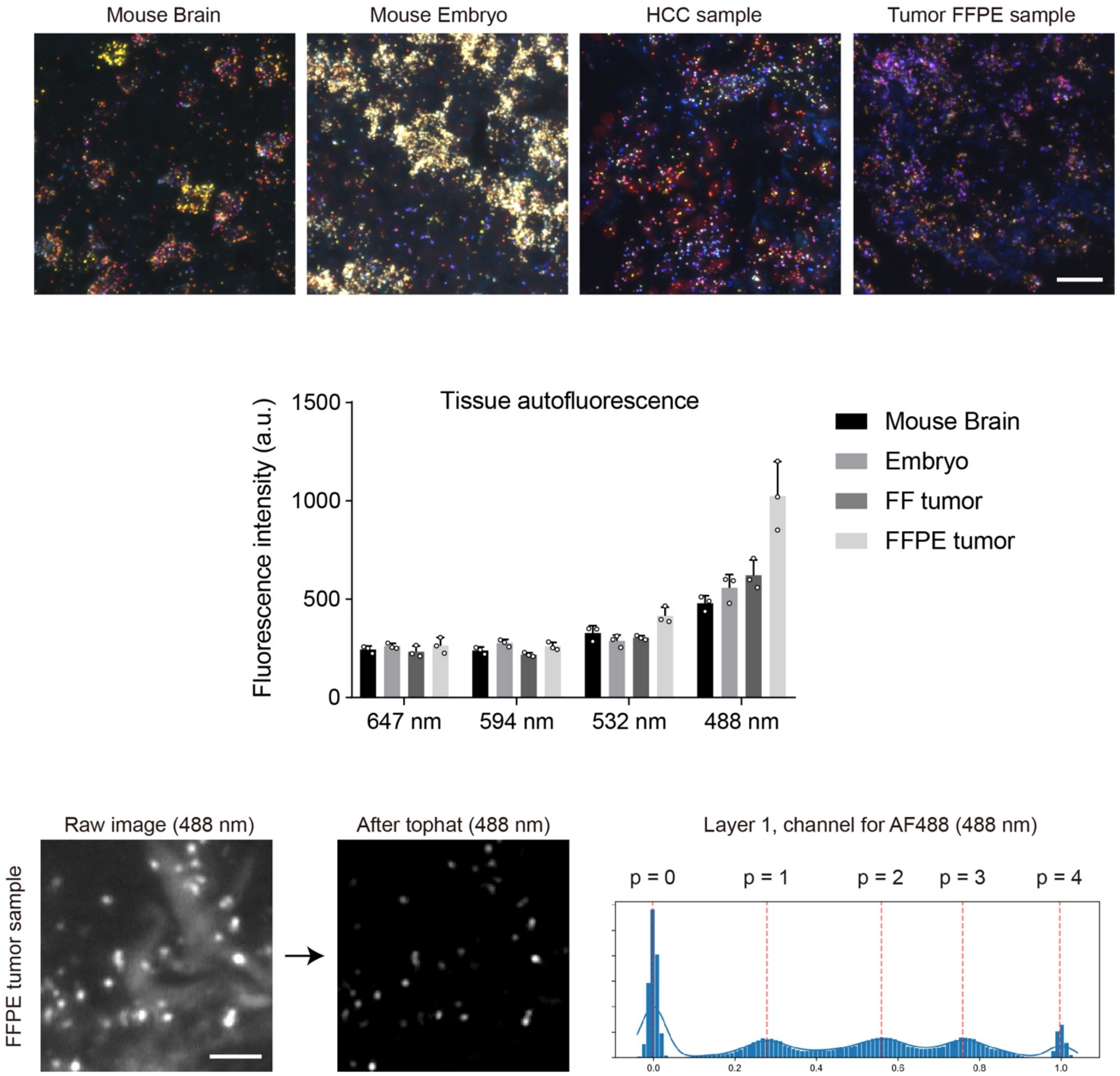


**Supplementary Fig. 39 | PRISM usability across diverse tissue types with different autofluorescence backgrounds.** PRISM's multi-level color coding was validated in diverse samples: mouse brain, mouse embryo and human tumor sample (fresh-frozen/FFPE). Tissue autofluorescence values were quantified across samples (n = 3 biological replicates for each sample), bar plots show mean value ± s.d. Among all channels used during imaging, AF488 channel (excitation wavelength 488 nm) generates the highest level of autofluorescence background. FFPE samples displayed the strongest autofluorescence across all tested samples. PRISM can robustly achieved multi-level color grading in the FFPE samples despite of its high autofluorescence background. Scale bar: 20 μm, above (overview); 5 μm, below (zoom-in image).

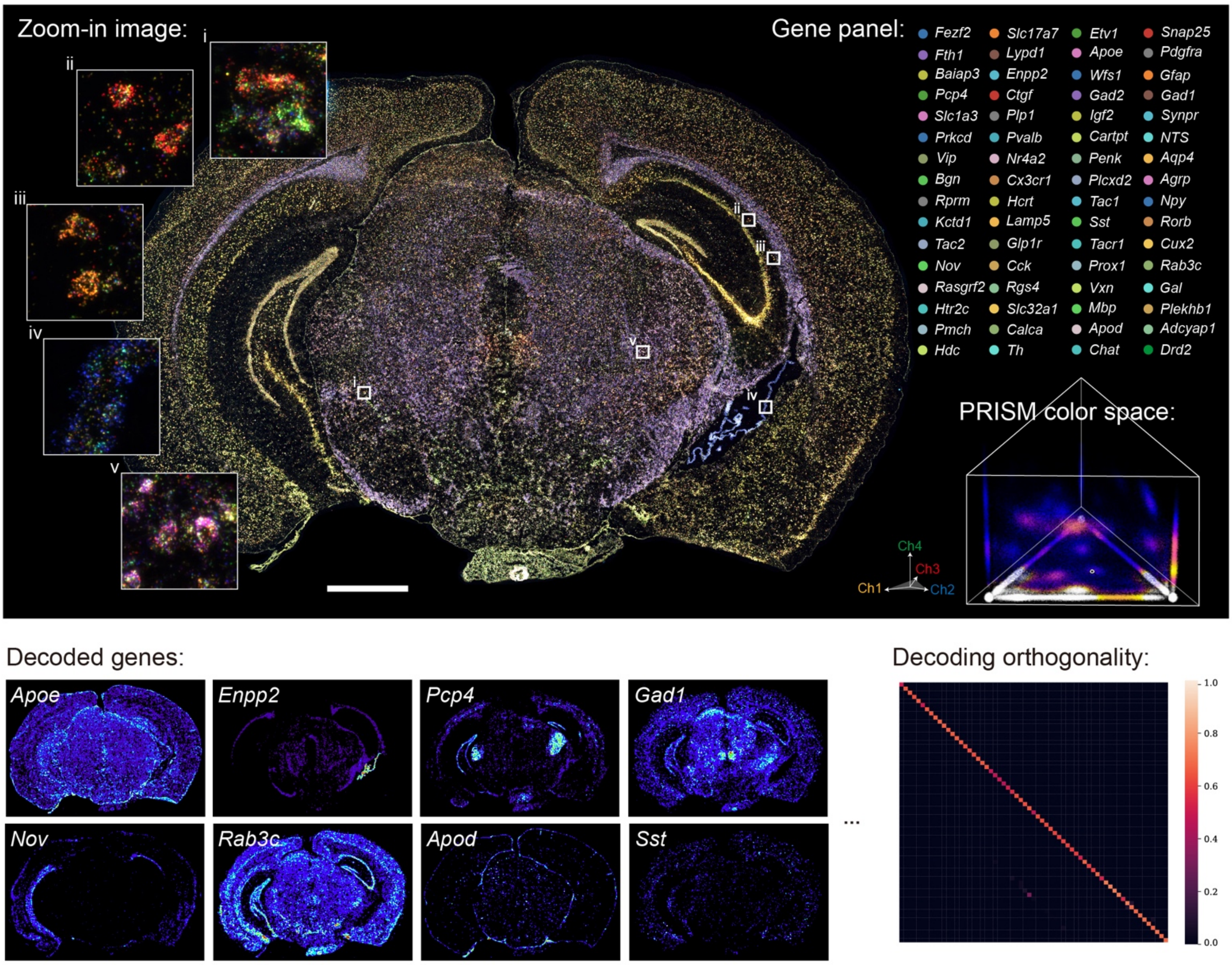


**Supplementary Fig. 40 | 64-plex RNA imaging in mouse brain tissue by PRISM.** Scale bar: 1 mm.

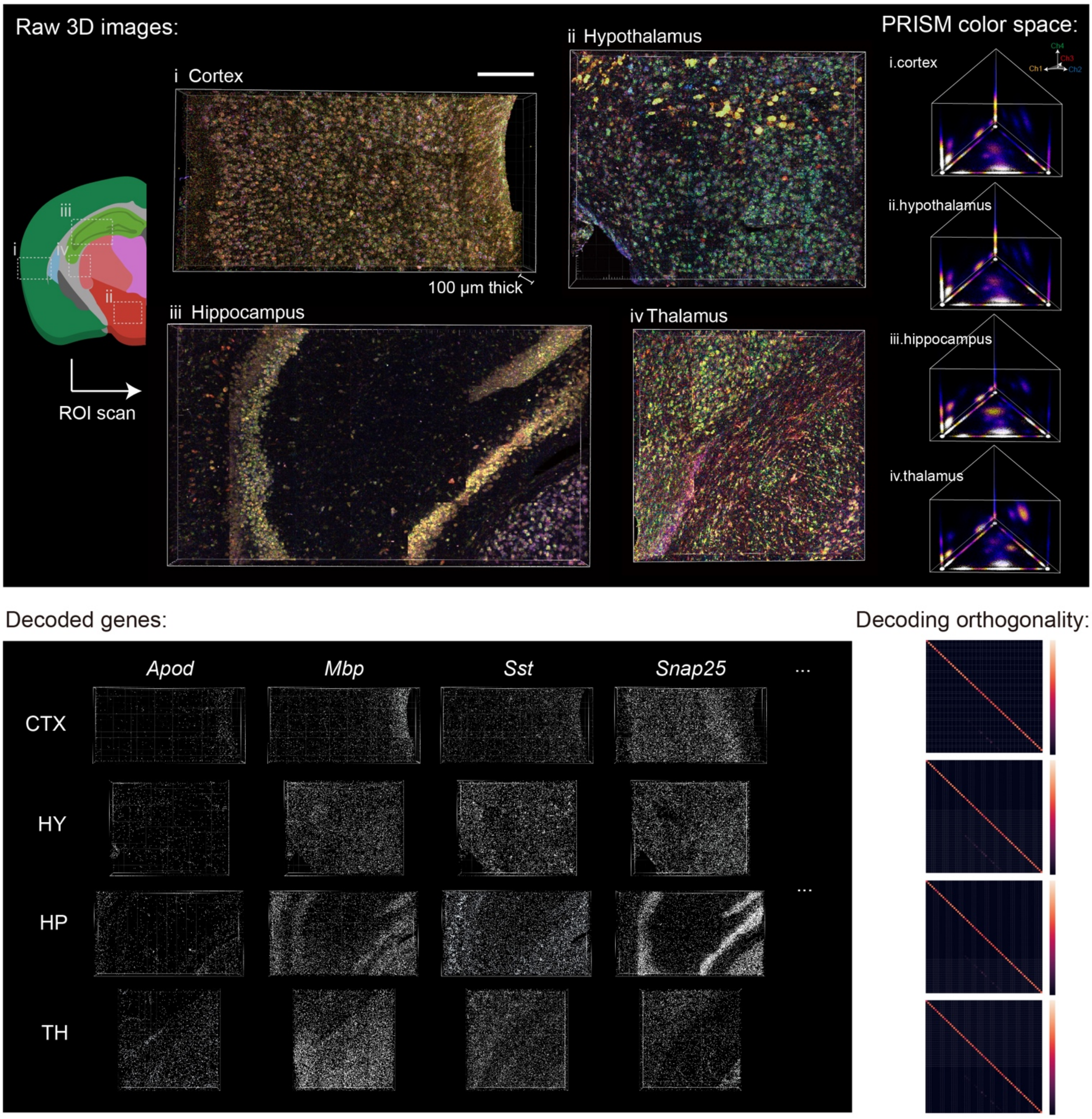


**Supplementary Fig. 41 | 64-plex RNA imaging in 100-µm thick tissue by PRISM.** The experimental workflow and decoding process were performed the same way as in Fig. 6. Imaging was performed with confocal microscopy. The 3D assessment was repeated three times with similar results. Scale bar: 200 µm.
